# Versatile detection of diverse selective sweeps with Flex-sweep

**DOI:** 10.1101/2022.11.15.516494

**Authors:** M. Elise Lauterbur, Kasper Munch, David Enard

**Affiliations:** Department of Ecology and Evolutionary Biology, University of Arizona, Tucson AZ 85719, USA; Bioinformatics Research Centre, Aarhus University, DK-8000 Aarhus C, Denmark

## Abstract

Understanding the impacts of selection pressures influencing modern-day genomic diversity is a major goal of evolutionary genomics. In particular, the contribution of selective sweeps to adaptation remains an open question, with persistent statistical limitations on the power and specificity of sweep detection methods. Sweeps with subtle genomic signals have been particularly challenging to detect. While many existing methods powerfully detect specific types of sweeps and/or those with strong signals, their power comes at the expense of versatility. We present Flex-sweep, a machine learning-based tool designed to detect sweeps with a variety of subtle signals, including those thousands of generations old. It is especially valuable for non-model organisms, for which we have neither expectations about the overall characteristics of sweeps nor outgroups with population-level sequencing to otherwise facilitate detecting very old sweeps. We show that Flex-sweep has the power to detect sweeps with subtle signals, even in the face of demographic model misspecification, recombination rate heterogeneity, and background selection. Flex-sweep detects sweeps up to 0.125 * 4N_e_ generations old, including those that are weak, soft, and/or incomplete; it can also detect strong, complete sweeps up to 0.25 * 4N_e_ generations old. We apply Flex-sweep to the 1000 Genomes Yoruba data set and, in addition to recovering previously identified sweeps, show that sweeps disproportionately occur within genic regions and close to regulatory regions. In addition, we show that virus-interacting proteins (VIPs) are strongly enriched for selective sweeps, recapitulating previous results that demonstrate the importance of viruses as a driver of adaptive evolution in humans.

## INTRODUCTION

The genomic legacy of past natural selection sets the stage for understanding evolutionary trajectories and current diversity. However, establishing the specific genomic contributions of past natural selection to the modern melange of adaptations and their patterns within genomes remains a difficult task. Fully characterizing these patterns requires that we know the genomic contributions of past positive natural selection, either in the form of polygenic adaptation or in the form of selective sweeps. However, in the specific case of sweeps, this presents two primary challenges: First, patterns of positive selection through sweeps tend to fade over time, and it has been particularly challenging to detect older hitchhiking events. Second, since genomic patterns of positive natural selection include diverse types of selective sweeps with varying stages of completeness, varying starting frequency, and varying ages, existing methods often detect a specific type of sweep(s) at the expense of versatility.

Designing detection methods that are capable of detecting diverse sweeps remains a challenge (Kern and Schrider 2018). Thus, not only do the specific targets of old positive selection events remain poorly known even in well studied species such as humans, any effort to characterize sweeps genome-wide is susceptible to bias resulting from the sweep type specialization of many methods that can result in false negatives (Akey 2009). More specifically, the task of pushing back the earliest dates of detectable selective events has recently become an important goal of population geneticists, particularly for humans (Racimo et al. 2014; Key et al. 2016; Racimo 2016; Cheng et al. 2017; Marciniak and Perry 2017; Peyrégne et al. 2017; Cheng et al. 2022), but also for non-model species (eg. Veale and Russello 2017; Hejase et al. 2020). In addition, while many disparate existing methods focus on detecting specific types of selective sweeps, the recent flourishing of machine learning and other composite statistical methods has brought the field to a methodological juncture that allows the development of a “Swiss Army Knife” approach to detect a greater diversity of sweeps with a single method.

A selective sweep describes the process of a beneficial allele increasing in frequency as a result of positive natural selection and the resulting patterns of genetic diversity around the swept locus (Maynard Smith and Haigh 1974). Statistical methods for identifying these patterns and distinguishing them from patterns caused by demographic changes, recombination, or background selection are both numerous and, for sweeps with strong signals, powerful (eg. Kim and Nielsen 2004; Nielsen 2005; Voight et al. 2006). These methods have been used to discover the genomic basis of a variety of adaptations in humans and other species, including adult lactase persistence (Enattah et al. 2002; Bersaglieri et al. 2004; Coelho et al. 2005), high altitude adaptation (Simonson et al. 2012; Hu et al. 2019; Li et al. 2014), and Malaria adaptation through the Duffy null allele (McManus et al. 2017; Hamblin and Di Rienzo 2000; Hamblin et al. 2002). In addition, these methods are used to characterize the nature of selection across the genome, including the relative frequencies of different types of sweeps (Hernandez et al. 2011; Schrider and Kern 2017) and overall strength and effects on diversity (Sattath et al. 2011).

However, as the time between a sweep and sampling grows, the genetic footprint dissipates (Przeworski et al. 2005). This erosion of signal with age makes old sweeps difficult to detect, which has so far limited the development of methods to detect the genomic signatures of selective events in the time frame between old, population-level selection, and speciation. (Here, we use “old” to refer specifically to selection that occurred more than 0.05 * 4N_e_ generations, corresponding roughly to 2,000 generations ago in humans). In principle, these signals should be present thousands of generations after the end of a selective sweep (Bisschop et al. 2021). Existing methods designed to detect sweeps that occurred beyond this time frame are largely based on differences between populations or closely-related species (Prüfer et al. 2014; Racimo et al. 2014; Racimo 2016; Peyrégne et al. 2017; Librado and Orlando 2018; Cheng et al. 2022). While powerful under some conditions, these methods typically rely on correlated allele frequency differences between populations, limiting selective sweep scans to populations with genomic data for outgroups and/or admixture. With these additional populations, selection can be detected in the ancestral branch before the population split(s), and localized to the ancestral or daughter branches. Similar methods can also identify adaptive introgression (Gower et al. 2021). Despite this, these methods do not perform well with soft sweeps (sweeps from standing genetic variation (Hermisson and Pennings 2005)) or weak selection (Peyrégne et al. 2017; Bisschop et al. 2021), and the lack of an appropriate outgroup for many non- model species limits their utility. Ancient DNA is another useful avenue for detecting old sweeps (Key et al. 2016; Marciniak and Perry 2017), but it too is of limited availability, including in humans, especially beyond 10,000 years.

Subtle signals are not limited to old sweeps but can also be caused by very incomplete sweeps, sweeps from standing genetic variation, and sweeps caused by weak selective pressure. Since genomic patterns of sweeps may include sweeps of all types, stages of completeness, and age; and because we don’t typically know what type(s) of sweeps to expect during a targeted analysis (for example, in the case of an older selective pressure such as an ancient epidemic), detection methods need to be able to detect a diversity of different types of sweeps with the highest sensitivity and specificity possible. For example, selection pressure causing an old sweep could have lifted or changed before the swept allele reached fixation, leaving the sweep incomplete but still of interest (eg, Souilmi et al. 2021).

The most power and promise to detect such sweeps comes from machine learning, including methods that combine multiple statistics and, more recently, ancestral recombination graphs. While using ancestral recombination graphs (ARGs) in a deep-learning framework is a promising method to detect sweeps and estimate selection coefficients (Hejase et al. 2022), it is likely to suffer at older time scales as increased uncertainty in ARG reconstruction might begin to limit its power. It is currently uncertain how great this effect is likely to be. Combining multiple statistics in machine learning (Pybus et al. 2015; Kern and Schrider 2018; Mughal and DeGiorgio 2018; Flagel et al. 2019; Mughal et al. 2020), Approximate Bayesian Computation (Racimo et al. 2014), or composite of multiple signals (Grossman et al. 2010; Sugden et al. 2018) approaches takes advantage of the strengths and ameliorates the weaknesses of individual statistics. Similar methods have been developed to distinguish between types of selection (Kern and Schrider 2018; Xue et al. 2021; Isildak et al. 2021; Caldas et al. 2022). Once swept regions have been detected, additional methods can be applied to determine their characteristics (eg. Peter et al. 2012; Kern and Schrider 2018; Sugden et al. 2018; Torada et al. 2019; Xue et al. 2021; Caldas et al. 2022).

In general, summary statistics developed to detect the signatures of selective sweeps fall into one of three categories: examining signatures in haplotype structure, the site frequency spectrum (SFS), and overall genetic diversity. These three types of signatures vary in how they are affected by standing variation, sweep completeness, background selection, recombination rate variation, and time since the beginning and/or end of the sweep (Chen et al. 2010; Ferrer-Admetlla et al. 2014; Racimo 2016). For example, the power of haplotype structure-based statistics is susceptible to the breakdown of linkage disequilibrium (LD) over time, as well as sweeps from standing variation since the sweep does not start from a single haplotype (Chen et al. 2010). In contrast, combining haplotype structure with other diversity metrics increases the power to detect soft and incomplete sweeps (Ferrer-Admetlla et al. 2014). These approaches that combine multiple statistics can thus take advantage of these various complementary strengths and weaknesses.

Haplotype-based statistics (e.g. iHS, Voight et al. 2006; nS_L_, Ferrer-Admetlla et al. 2014; HAF, Ronen et al. 2015; H12, Garud et al. 2015; and iSAFE, Akbari et al. 2018), however, are especially robust to background selection. Background selection, i.e. negative selection against deleterious alleles that reduces genetic diversity at linked sites as they are lost with the deleterious alleles (Charlesworth et al. 1993), can be particularly problematic because it has strong potential to confound selective sweep inference via other types of summary statistics (Stephan 2010). However, background selection does not increase the frequency of long haplotypes, as expected in genetic hitchhiking with a selective sweep (Enard et al. 2014), so haplotype-based statistics are unlikely to be confounded. Thus, their inclusion in a multi-statistic based machine learning approach should increase its robustness to background selection (Schrider 2020).

Here, we develop a convolutional neural network (CNN) based machine learning method that is designed to detect diverse sweeps. This includes sweeps between 0.05 * 4N_e_ generations old (∼2,000 human generations) and 0.125 * 4N_e_ generations old (∼5,000 human generations), as well as those with subtle signals caused by other factors, including those that are very incomplete, those that started from a high level of standing variation, and those that are relatively weak. This method, Flex-sweep, makes this possible through the combination of six existing statistics and five new haplotype-based statistics, all with different strengths to detect selective sweeps with different characteristics. These five new statistics (*hapDAF-o, hapDAF-s, Sratio, lowfreq, and highfreq*) take advantage of the hitchhiking to higher frequency of alleles in regions linked to an adaptive allele by comparing diversity between ancestral and derived haplotypes.

Flex-sweep doubles reliable sweep detection time from 0.05-0.075 * 4N_e_ generations ago (∼2,000 – 3,000 human generations, e.g. Sugden et al. 2018; Harris and DeGiorgio 2020) to at least 0.125 * 4N_e_ generations ago (∼5,000 human generations), and up to 10,000 human generations ago for strong, hard sweeps. In addition, it accurately detects sweeps of a diverse range of time (including recent sweeps), completeness, initial allele frequency, and selection strength, making it a versatile tool. We show its robustness to demographic model misspecification, background selection, and recombination rate variation, all vital considerations for inference in any population genomic context (Johri et al. 2021). This expands our ability to study genome-wide patterns of selective sweeps, as well as the origins of phenotypically important loci, including adaptations to major historical selective pressures such as diet, climate, and ancient epidemics. Flex-sweep requires only one population being sequenced, so we expect it to be particularly useful in the context of non-model species where outgroups with population-level sequencing are often not available.

## NEW APPROACHES

Here, we present a novel selective sweep detection method that provides a versatile tool to identify genomic windows involved in a selective sweep up to 0.125 * 4N_e_ generations old (approximately 5,000 human generations), or a strong, hard sweep up to 0.25 * 4N_e_ generations old (approximately 10,000 human generations). This convolutional neural network-based method, Flex-sweep, uses a variety of types of statistics. In particular, the haplotype-based statistics provide robustness to background selection, a common confounder of sweep detection methods.

We introduce five new statistics designed to detect old, soft, and incomplete sweeps, which have sensitivities that complement those of existing statistics in detecting sweeps of different types. It is this combination of statistics that makes this method versatile in detecting sweeps of a variety of ages, completeness, starting allele frequency, and strengths. The combination of these statistics in a convolutional neural network, with an architecture optimized for versatility in sweep detection, makes this tool an all-purpose sweep detection method that is robust to background selection, recombination rate variation, and demographic model misspecification.

This versatility makes our new approach complementary to existing sweep classifiers and particularly useful for studies of non-model species for which we do not have strong empirical expectations of the characteristics of sweeps present in the genome. It can be used in conjunction with methods designed to classify or quantify sweep characteristics (for example, complete vs. incomplete, hard vs. soft, sweep age, or selection strength): Flex-sweep can first detect sweeps with high confidence, increasing the proportion of all sweeps detected in the genome; then the characteristics of these sweeps can be determined with greater certainty.

Flex-sweep is available at https://github.com/lauterbur/Flex-sweep. A singularity container and pre- trained equilibrium and Yoruba models has been deposited in Zenodo at DOI 10.5281/zenodo.7860595.

## RESULTS

Flex-sweep combines 11 summary statistics, including five newly-defined statistics that we show have good power to detect selective sweeps with certain characteristics that complement the power of existing summary statistics. We show that the application of these statistics in a convolutional neural network framework can detect selective sweeps with subtle signals, and is robust to common confounders such as demographic model misspecification, recombination rate heterogeneity, and background selection.

### New Statistics

We present five new statistics designed to detect the signatures of diverse sweeps and sweeps with attenuated signals designed to complement the published statistics (Table 1). See **supplementary results** for details on the power of each statistic at different combinations of sweep age, starting allele frequency, ending allele frequency, and strength.

**Table 1.** Summary statistics used in Flex-sweep.

| summary statistic | description | summarizes | citation |
| --- | --- | --- | --- |
| iHS | integrated haplotype score | haplotype structure | (Voight et al. 2006) |
| nSL | number of segregating sites by length | haplotype structure | (Ferrer-Admetlla et al. 2014) |
| DIND | derived intra-allelic nucleotide diversity | diversity on derived background | (Barreiro et al. 2009) |
| iSAFE* | integrated selection of allele favored by evolution | haplotype structure | (Akbari et al. 2018) |
| HAF** | haplotype allele frequency | haplotype structure | (Ronen et al. 2015) |
| H12 | frequencies of first and second most common haplotypes, modified to use 80% identity threshold | haplotype structure | (Garud et al. 2015) |
| hapDAF-o | haplotype derived allele frequency (old) | SFS | this paper |
| hapDAF-s | haplotype derived allele frequency (standing) | SFS | this paper |
| Sratio | segregating sites ratio | diversity on derived background | this paper |
| lowfreq | low frequency alleles on derived background | diversity on derived background | this paper |
| highfreq | high frequency alleles on derived background | diversity on derived background | this paper |

\* When a window included fewer than 300 SNPs, the iSAFE statistic was not calculated and the SAFE statistic was used instead. This is in accordance with Akbari et al. 2018's recommendation to use SAFE instead of iSAFE when there are few segregating sites, and SAFE is used in this circumstance for calculating all feature vectors (from the training data as well as from data to be classified).
\*\* Only the top 10% of HAF values are used, as this provides a better signal for incomplete sweeps.

### CNN Structure

The use of Convolutional Neural Networks (CNNs) for sweep detection and other population genetic inference has been comprehensively described by Kern and Schrider 2018 and reviewed in Schrider and Kern 2018 and Flagel et al. 2019. Flex-sweep calculates multiple summary statistics across a locus to create a feature vector, which is then used as input data for the CNN. We explored multiple possible feature vector scaling strategies and CNN architectures to select one with both good performance and evidence that it did not overfit the training data (see **Methods**; Supplementary figures 1 and 2). The final model architecture is described in the Methods, Appendix and available on GitHub (https://github.com/lauterbur/Flex-sweep). In addition, we found that training data sets with 10,000 neutral and 10,000 sweep simulations provided the best power, with little improvement with larger data sets (Supplementary figure 3).

### Performance

Flex-sweep has good power to find sweeps under multiple demographic scenarios when trained on simulations from an equilibrium demography with N_e_ = 10,000 (Figure 1). We trained a model with neutral and sweep data simulated under this scenario (a population with no history of population size change), including sweeps with an age from 0 – 5,000 generations (0.125 * 4N_e_), incomplete sweeps, sweeps from standing variation, and sweeps resulting from weak selection. When used to classify data simulated under the same equilibrium demography, Flex-sweep is able to detect 83.7% of sweeps with a 2.3% false positive rate (FPR; Figure 1a). The FPR increases when this equilibrium demographic-trained model is used to classify data with a history of population decline, expansion, or the complex Yoruba demographic history inferred by (Speidel et al. 2019) (Figure 1b, c, d). Sweep age, completeness, and starting allele frequency have the strongest impacts on power, with power declining as sweep age or starting allele frequency increase or sweep completeness decreases, especially at false positive rates below 1% (Figure 2). In particular, it struggles at the most restrictive false positive rate of 0.1% to detect very incomplete sweeps (20 - 40% complete), very old sweeps (4001 - 5000 generations old), sweeps from high starting allele frequency (7.5 - 10%), and very weak sweeps (0.001 <= *s* <= 0.0028), but is still able to detect most sweeps at a more relaxed false positive rate of 1% or 2% (Figure 2). Calibration analysis suggests that these estimates of power, especially at very low false positive rates, are conservative and that further refinements of the model could improve these figures. See **supplementary results** for details on calibration and additional detail on detecting older sweeps.

**Figure 1.**
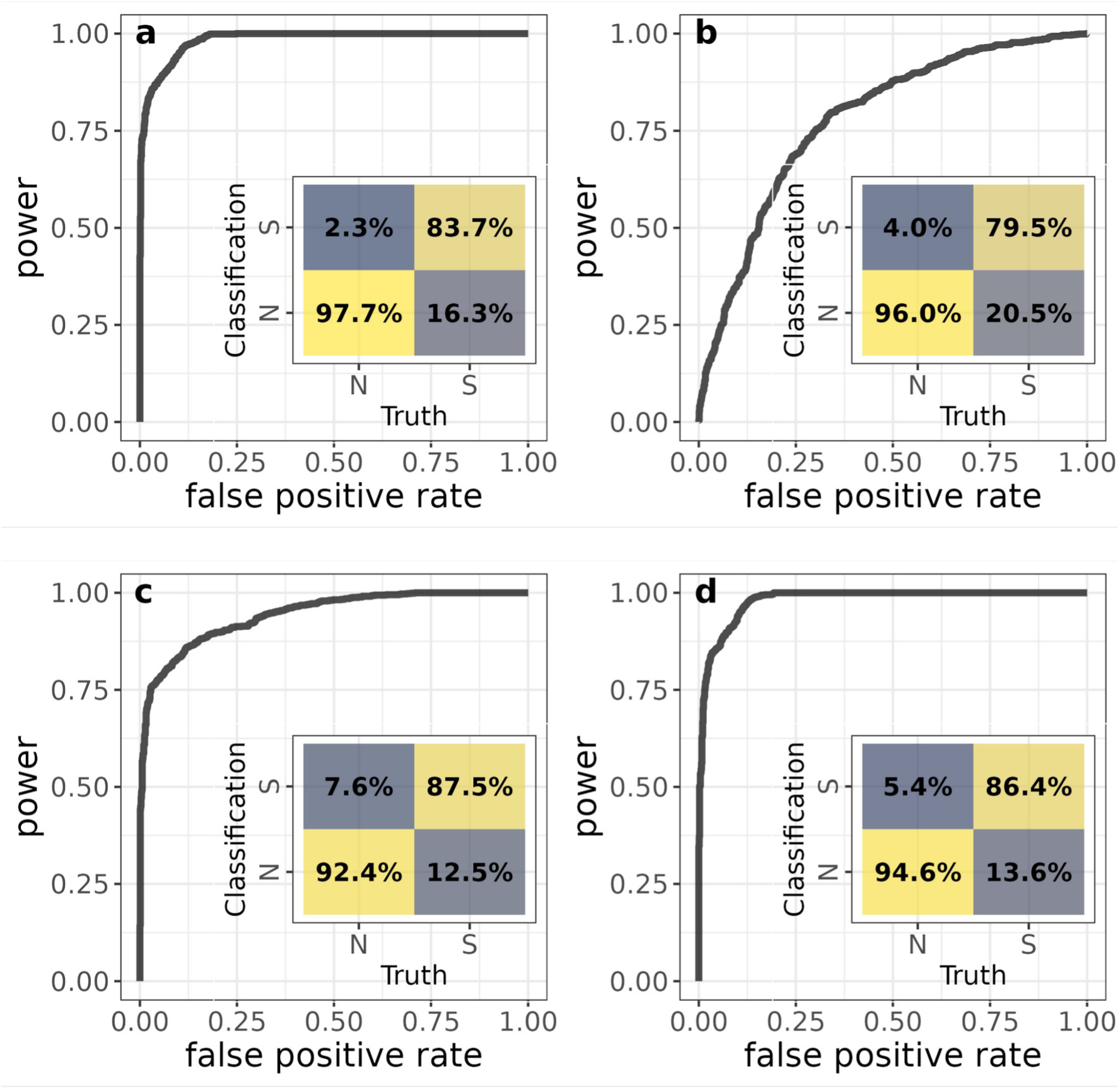
Power and false positive rates of the CNN trained on an equilibrium demographic scenario: (a) Classifying simulations from the same equilibrium demographic scenario, (b) 5x decline, (c) 10x expansion, and (d) Yoruba demographic history after Speidel et al. 2019. The inset confusion matrices show, for each scenario, the classification rate of each region type (neutral, N, or sweep, S, x axis) as neutral or sweep (N or S, y axis), with false positives (neutral regions classified as sweeps) in the upper left, and false negatives (sweep regions classified as neutral) in the bottom right. Darker blue indicates smaller values, brighter yellow indicates larger values.

**Figure 2.**
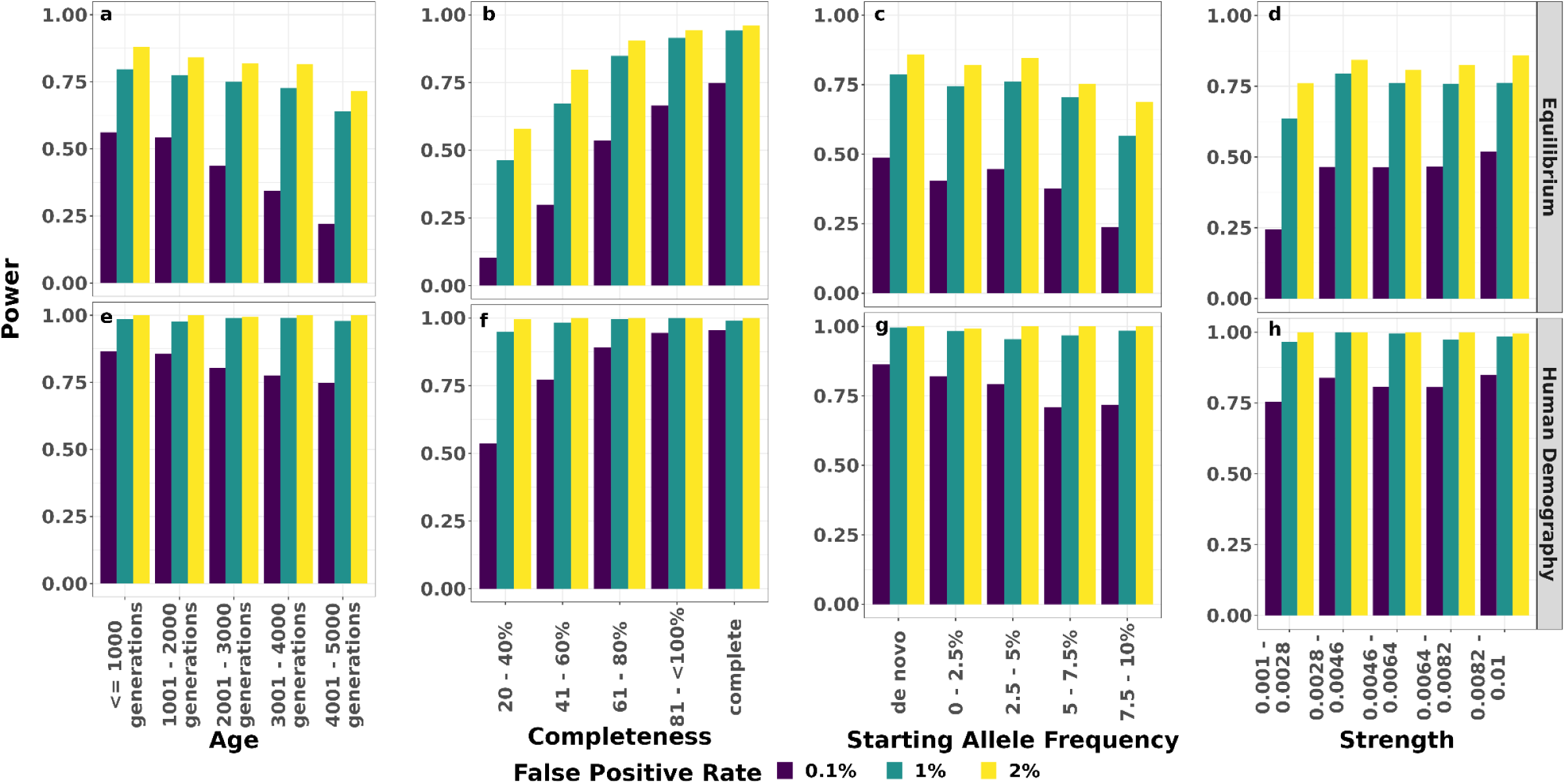
Power to detect sweeps of different types. Power in a correctly specified equilibrium model (top) and the correctly specified human demographic model (bottom). Sweep types vary in age (a, e), completeness (b, f), starting allele frequency (c, g), and strength (d, h) at 0.1% (purple), 1% (teal), and 2% (yellow) false positive rates. In each panel, sweep characteristics otherwise come from the full distributions of each sweep parameter. False positive rates are those presented in the corresponding ROC curve figures.

While the model has good power to detect sweeps even when misspecified with an equilibrium demographic history (Figure 1), some aspects improved when using a generic data set that includes simulations from multiple types of population size history (decline, expansion, and equilibrium), and improved even more when trained with correctly specified demographic models (Figure 3). In particular, training with data simulated under a combination of population histories (Figure 3a) decreases the false negative rate when used to predict data simulated under the Yoruba demographic model, but at the expense of a higher false positive rate. In comparison with a model trained with equilibrium data, the false negative rate for classifying sweeps in a population that underwent a decline decreases when the model is trained with various magnitudes of population decline (Figure 3b) and the false positive rate decreases when training with and classifying expansions (Figure 3c). When a more complex demographic model for the training data is correctly specified, as by using the Yoruba demographic history inferred by Speidel et al. 2019 for both training and testing data (Figure 3d) the false positive and false negative rates decrease. Flex-sweep’s performance suffers with the severe mis-specifiation of using a model trained with population expansions to classify sweeps from a population that underwent a population decline (Supplementary figure 4). Mispolarization of variants is especially likely to be a concern in non-model species with poorly sequenced outgroups, however we find that low rates of mispolarization (0.1%, 1%) have little effect on classification power and false positive rates, while higher rates (5%, 10%) decrease FPR and increase power (Supplementary figure 5).

**Figure 3.**
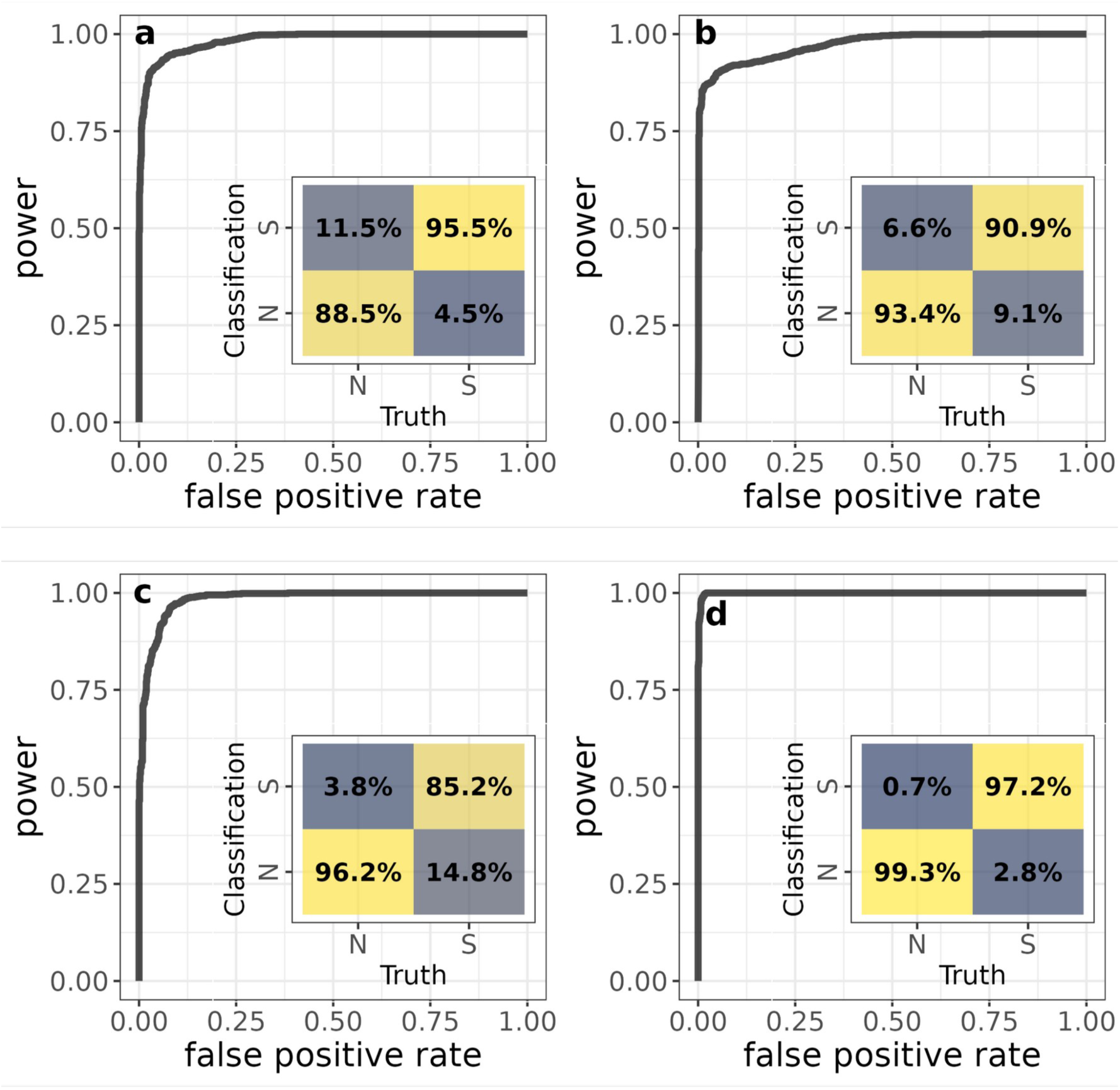
Power and false positive rates for Flex-sweep trained with non-equilibrium demographic scenarios. (a) A combined data set made up of simulations with a history of population declines, expansions, and equilibrium demography, used to classify sweeps on the Yoruba population history (b) population decline (1x, 5x, 10x), used to classify sweeps on a history of 5x population decline (c) expansions (5x, 10x, 50x), used to classify sweeps on a history of 10x population expansion, and (d) the Yoruba demographic history, used to classify sweeps on the Yoruba demographic history.

Sweep time, strength, SAF, and EAF all affect what simulated sweep regions are likely to be misclassified as neutral, and that while mutation rate does not affect which regions are likely to be misclassified, low recombination rates are more likely to generate false positives (t-test, p < 0.001, Supplementary figures 6, 7). In addition, sweeps that are both older and less complete are less likely to be detected than younger, more complete sweeps (Supplementary figure 8). While entire regions of low recombination rate have higher false positive rates, the impact is less when they are heterogenous with regions of high recombination rate (Supplementary figure 9, 10). The impact of low recombination rate can be drastically reduced by increasing the confidence threshold at which a region is considered to contain a sweep (Supplementary figure 10a), or by training with a low recombination rate model (Supplementary figure 11, see **supplementary results**). Furthermore, background selection has little influence on false positive rate (Supplementary figure 9a, b), regardless of exon structure or recombination rate (Supplementary figure 10b).

We also compare Flex-sweep to diploS/HIC, another powerful CNN-based method (Kern and Schrider 2018) and one of the first to popularize CNNs for detecting selective sweeps (see **Methods** and **supplementary methods**). Flex-sweep has greater power and lower false positive rates than this modified, binary (sweep or neutral) diploS/HIC and, as expected because it was not diploS/HIC’s purpose, this difference is more pronounced in data sets that include incomplete sweeps (Figure 4, supplementary figure 12). It would be ideal if Flex-sweep were able to also distinguish between types of sweeps, but we find that attempting to do so decreases power and increases false classifications (Supplementary figure 13). See **supplementary results** for more detail.

**Figure 4.**
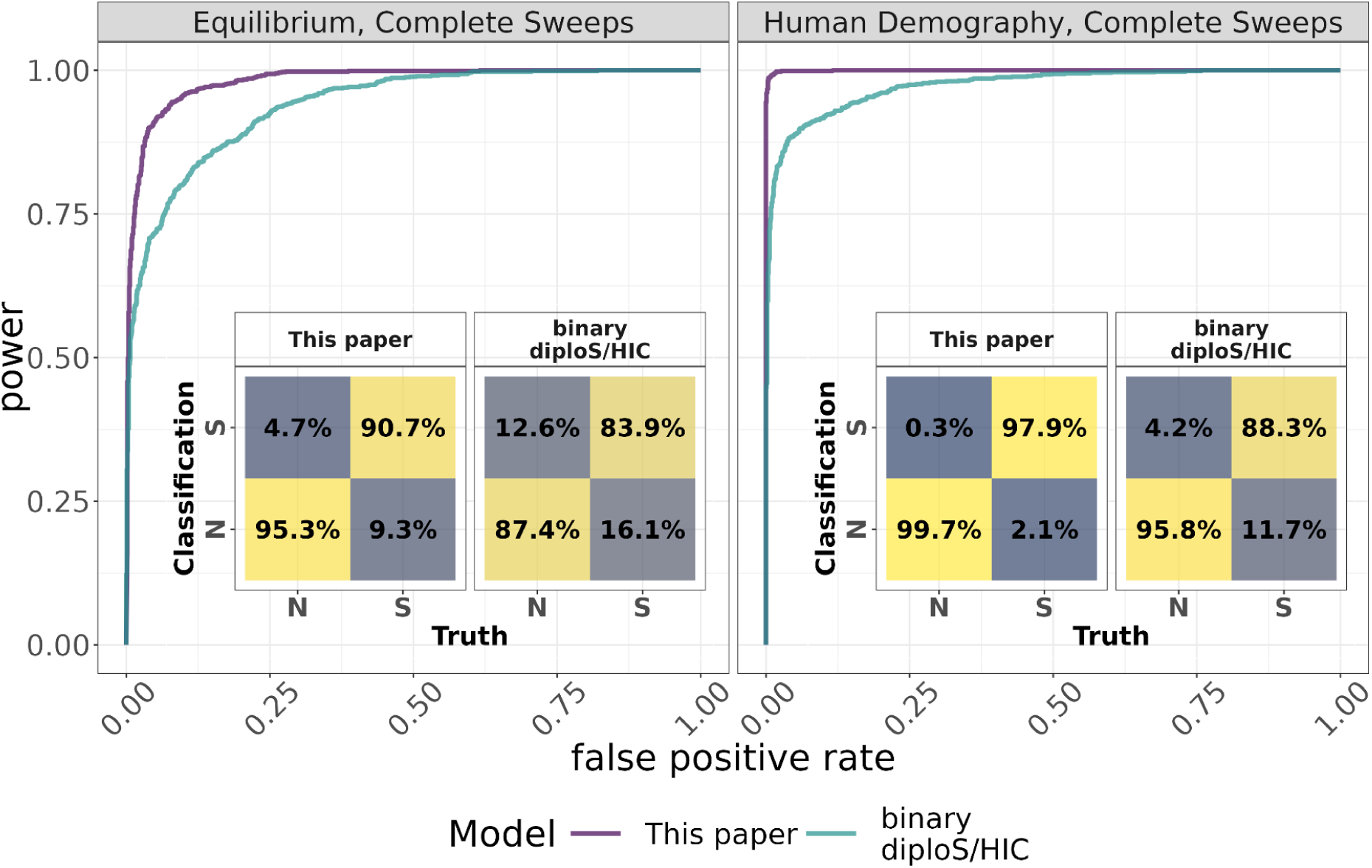
Power and false positive rates comparing Flex-sweep and diploS/HIC. Both were trained and tested on the same data sets using an equilibrium demography with only complete sweeps (left), and the human demography data set with only complete sweeps (right).

To investigate the influence and impact of the different statistics on the model predictions, we compared the classifications results of the network using only the new statistics vs. only the previously published statistics, the order of the statistics in the feature vector, and explored how each statistic contributes to Flex-sweep classification using Deep SHAP (Lundberg and Lee 2017). The combination of all 11 statistics has substantially better power than either group alone (Supplementary figure 14), and ordering the statistics such that those that summarize the data differently are interspersed with each other among the rows improves classification, while grouping them worsens classification (Supplementary figure 15). The most important features influencing classification, regardless of training, are *highfreq* and *hapDAF*-s calculated for the middle as well as the shoulders of the locus (Supplementary figure 16). See **supplementary methods** and **supplementary results** for more detail.

### Application

We applied Flex-sweep to the 1000 Genomes data set of Yoruba in Ibadan, Nigeria (The 1000 Genomes Project Consortium et al. 2015). While Flex-sweep may be of particular practical interest to studies of adaptation in non-model organisms, we chose this data set and population because it has been widely used in previous studies both to identify novel selective sweeps and to benchmark new selective sweep detection methods (eg. Ferrer-Admetlla et al. 2014; Racimo 2016), as well as includes many of the sort of demographic changes that we expect to see in other organisms. Because Flex-sweep is optimized to detect very old sweeps (up to 0.125*4N_e_ generations) of diverse types (both *de novo* and from standing variation, various levels of completeness, and a wide range of strengths, we use it to test the hypotheses that selective sweeps are expected to occur 1. disproportionately in genic regions, and 2. Disproportionately close to regulatory regions.

To understand the patterns of selective sweeps across the genome, we calculated the proportions of swept regions in genes and their distances from sites associated with regulatory activity. We find that selective sweeps are significantly more likely to be found within genes than would be expected by chance at all tested confidence thresholds (one-tailed randomization test (see **Methods**), all p < 0.01, Figure 5, Supplementary figure 17). Across the genome, 54% of sweeps are found within genes, while the expected mean by randomization is 28% at the default confidence threshold (Figure 5, see **Methods**). In addition, the peaks of each sweep region (see **Methods**) are more likely to be closer to the transcription start site at the default confidence threshold (Figure 6, Supplementary figure 18, one-tailed randomization test p = 0.02, see **Methods**). See **supplementary results** for additional detail using higher confidence thresholds.

**Figure 5.**
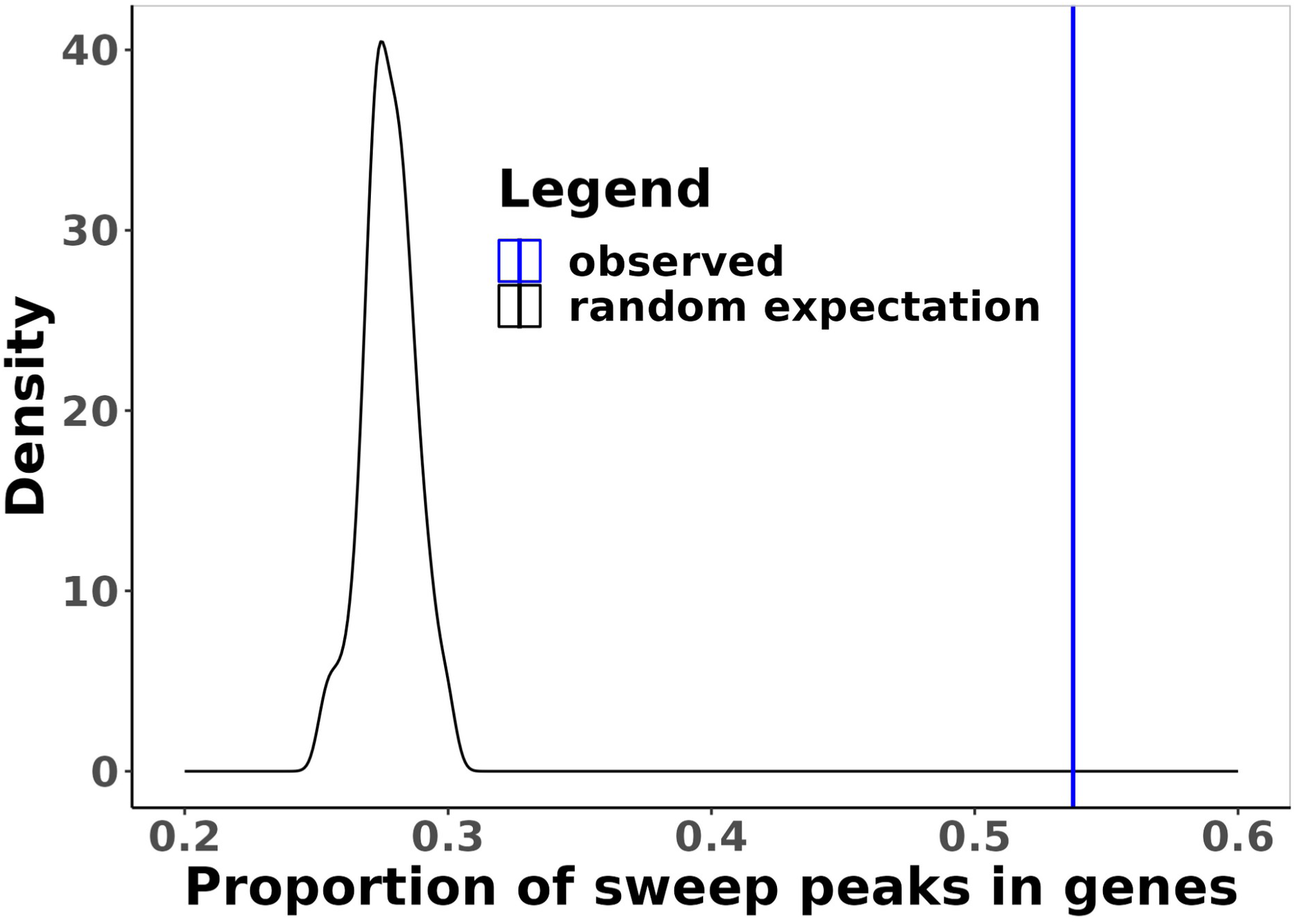
The proportion of sweeps inside genes. The observed proportion of selective sweeps within genes (blue line) compared to the distribution expected by randomization (black line) at the default classification confidence threshold of 0.5.

**Figure 6.**
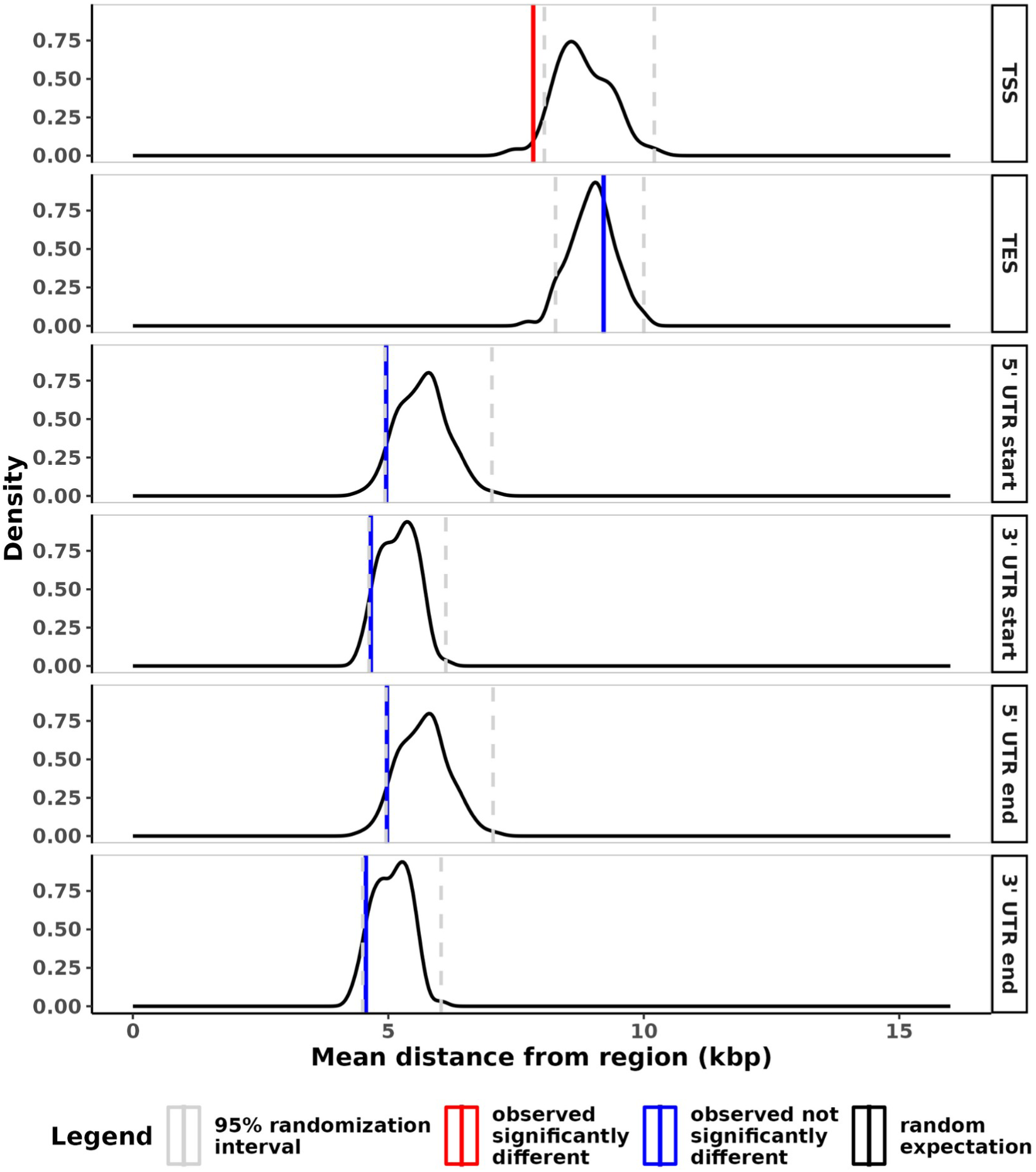
The distance of sweep peaks to regulatory regions. The observed mean distance (red and blue lines) of the peak of each swept region to the nearest start and end site of each type of regulatory region: transcription, 5’ UTR, and 3’ UTR; compared to the distribution expected by randomization (black line) at the default classification confidence threshold of 0.5. Vertical dashed lines indicate the lower and upper 2.5% of the randomized data.

In addition, we estimated a false discovery rate (FDR) metric to demonstrate Flex-sweep’s performance by comparing the false positive rate from the validation tests using the Yoruba demographic model to the proportion of windows classified as sweeps in sets of independent windows 1 Mb apart in the Yoruba data. The FDR is then calculated from these two values as FDR = # false positive windows / total # positively classified windows (see **Methods**). We can use Flex-sweep’s confidence score to determine the threshold that would correspond to a tolerance for a specific false discovery rate. During training on the human demographic model, Flex-sweep had a 0.7% false positive rate (FPR = number of false positives / total number of positive classifications). Averaging over 100 sets of independent windows 1 Mb apart in the Yoruba data, the average FDR is 1.9% (see **Methods** and **supplementary methods**). Obtaining a false discovery rate below 1% would require increasing the confidence threshold to approximately 0.8 (Supplementary figure 19).

We must nevertheless consider the likely possibility that the false discovery rate may well be much higher when scanning actual sequencing data compared to simulation estimates. The proportion of the human genome classified in a sweep is indeed arguably high, but this may reflect the increased power to detect regions of the genome that overlap diverse types of sweeps, with more sensitivity and the ability to detect sweeps further back in time. Many windows also likely represent the impact of hitchhiking further away from the actually selected variants due to the increased detection power. When looking only at windows with the strongest possible sweep classification score of one, the percentage of genomic windows with such decisive sweep evidence falls from 28% to 1.9% (Supplementary figure 20).

To further test if Flex-sweep does identify true positive sweeps in the Yoruba genome, given the high proportion of the genome classified in sweeps, we compared the occurrence of sweeps at genes that interact with viruses (noted VIPs for Virus-Interacting-Proteins), where a strong excess of sweeps has previously been reported (Schrider and Kern 2017; Enard and Petrov 2020). If the sweep classifications in the Yoruba genome are over-run with false positives, we should not be able to detect an excess of sweeps at VIPs, because a high false positive rate would erase the true difference between VIPs and the rest of the genome. Using the same approach as reported in Enard and Petrov 2020, we find a very substantial sweep enrichment at VIPs (Figure 7). The observed enrichment is highly unlikely, with an estimated false discovery rate of 0.0006. The FDR estimation approach was previously described in Enard and Petrov 2020; Di et al. 2021. The enrichment is in particular greater than three-fold for sweeps with a very strong confidence score equal to one. This shows that, even if the false discovery rate may well be higher than estimated from simulation-based false positive rates, a substantial proportion of the sweep classifications have to be true positives. The very strong enrichment of VIPs in high-confidence sweep classifications is also in line with previous observations that viruses tend to drive strong selective sweeps reflecting strong selection (Enard and Petrov 2020; Souilmi et al. 2021).

**Figure 7.**
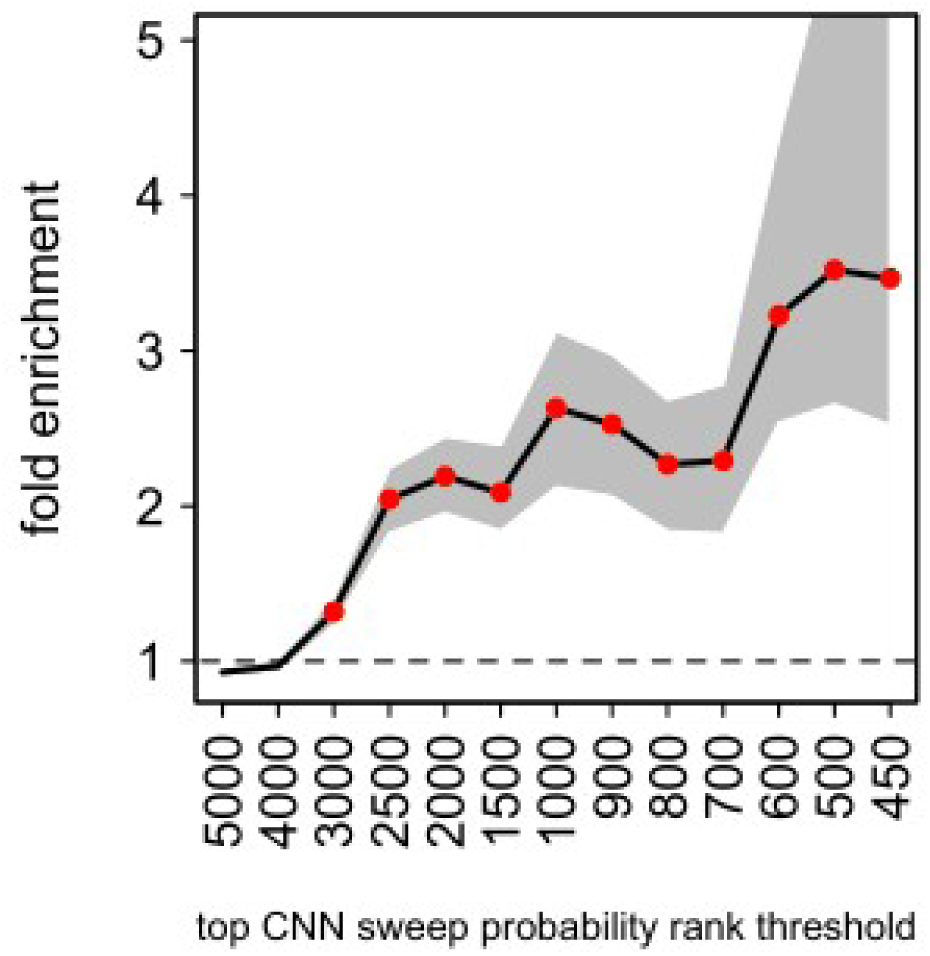
Flex-sweep sweep enrichment at VIPs relative to control non-VIPs. The number of VIPs in sweeps is compared to the average number in random control sets of non-VIPs for increasing CNN sweep confidence rank thresholds. A gene (VIP or non-VIP) was considered to be subject to a sweep if its genomic center of the gene was covered by at least one window classified as a sweep. The x axis represents the stringency of candidate genes considered, from lower stringency (more genes considered) on the left to higher stringency (fewer genes considered) on the right. Stringency is determined by ranking the genes in order of sweep confidence and choosing the top X ranked. The y axis represents the fold enrichment of VIPs compared to control non-VIPs in group of candidate genes, on average across all sets of non-VIPs, as described in Enard and Petrov 2020. Multiple neighboring VIPs in the top X may only represent one sweep, but this clustering is accounted for in the false discovery rate estimation as previously described (Di et al. 2021). Grey area: 95% confidence interval of the fold enrichment. Red dots: fold enrichment test P<0.001.

Furthermore, we recover multiple regions that have been previously implicated in sweeps with functional support. These include the known sweep at the high recombination locus of the Duffy Antigen Receptor for Chemokines gene (*DARC*), which is also the receptor for the malaria parasite *Plasmodium vivax*. This gene has been previously implicated as the target of a near-complete selective sweep, with the null (nonexpression) allele conferring malaria resistance (McManus et al. 2017). We find that a 430 kbp region surrounding the *DARC* locus and the causative promoter SNP rs2814778 is affected by a selective sweep at the default sweep confidence threshold of 0.5. At the more stringent threshold of 0.99, the region narrows to 370 kbp and still includes the *DARC* locus and rs2814778 (Figure 8). The edges of the sweep correspond to recombination hotspots 5’ and 3’ from the *DARC* locus. In addition, we find evidence of sweeps with high confidence at multiple genes with previous genomic and functional evidence of selection. Those supported by both lines of evidence include genes associated with responses to infectious diseases and pathogens, see **supplementary results** for more detail.

**Figure 8.** The region around the DARC gene on chromosome 1, using hg19 coordinates. The DARC gene is shown in black, with the causative polymorphism site rs2814778 shown as a red point. The entirety of the DARC gene, including rs2814778, lies within the swept region classified with high confidence by Flex-sweep. The grey line shows recombination rates in cM/Mb.

## DISCUSSION

Flex-sweep is a new and versatile deep-learning based method for detecting diverse selective sweeps of particular practical interest to studies of adaptation in non-model organisms. Because Flex-sweep is designed to detect sweeps of diverse types and characteristics, it is ideal for species for which we have few empirical expectations for the nature and types of selective sweeps to be detected. We show its power to detect sweeps under multiple demographic models, including demographic model misspecification, as well as its robustness to background selection and recombination rate heterogeneity. In addition to its broad versatility, it can detect sweeps at least 0.125 * 4N_e_ generations old (∼5,000 human generations), greatly expanding the potential to study the genomic legacy of ancient natural selection. We demonstrate Flex-sweep by identifying sweeps genome-wide using the 1000 Genomes project data set of Yoruba in Ibadan, Nigeria (The 1000 Genomes Project Consortium et al. 2015) and characterize the patterns of sweeps relative to genic and gene regulatory regions, and at virus-interacting proteins (VIPs). While Flex-sweep may be of particular practical interest to studies of adaptation in non-model organisms, this data set and population have been widely studied and includes many demographic changes that we expect to see in other organisms, so it provides a useful test case. We recapitulate previous results that sweeps are likely to occur in genic regions (eg. Fagny et al. 2014) and close to gene regulatory regions (eg. Peyrégne et al. 2017), as well as in VIPs (eg. Barreiro et al. 2008, Enard et al. 2016).

Supervised machine learning methods have become a popular approach for detecting patterns in population genetic data, including selective sweeps (Kern and Schrider 2018; Torada et al. 2019 ; Xue et al. 2021), recombination (Gao et al. 2016; Adrion et al.2020), population demographic history and structure (Wang et al. 2021), population assignment (Chen et al. 2018; Sylvester et al. 2018), and assignment to continuous geographic regions (Battey et al. 2020). These methods typically benefit from the use of multiple powerful summary statistics to take advantage of their various complementary strengths and weaknesses. This gives such methods the particular potential to detect diverse patterns, including the broad diversity of selective sweeps present throughout the genome. However, most methods to detect sweeps focus on one or a few types, for example, hard vs. soft sweeps or partial sweeps (Kern and Schrider 2018; Xue et al. 2021). While distinguishing between sweep characteristics provides useful information, it may reduce the power to detect sweeps with attenuated signals, as we show here.

We show that Flex-sweep is versatile in terms of the types and ages of sweeps it is able to detect, at the expense of determining the qualities of the sweep. To determine if a locus contains signatures of a sweep, it uses 11 statistics, including five new statistics, calculated over nested windows of five sizes (see **Methods**). These five new statistics (*hapDAF-o, hapDAF-s, Sratio, lowfreq,* and *highfreq*) have good power to fill in some detection gaps of existing statistics, and otherwise complement these statistics in the context of composite, multi-statistic detection methods such as machine learning. Calculating the statistics over nested windows of multiple sizes allows the neural network to take advantage of the differences in power statistics can have depending on window size. For example, Shapely analyses reveal that the importance of windows of different sizes can vary depending on the data used to train and test the model. Since we do not attempt to distinguish between sweep types or regions linked to selected loci in order to maintain the most power to detect a diversity of sweeps, other methods can be applied subsequently to determine sweeps’ nature (Kern and Schrider 2018; Xue et al. 2021; Caldas et al. 2022), location (Akbari et al. 2018; Sugden et al. 2018), strength (Torada et al. 2019; Caldas et al. 2022), and/or timing (Speidel et al. 2019). Flex-sweep may be combined with one or more of these methods by first identifying a swept region using Flex-sweep and then applying a more targeted method on the resulting candidate regions to further investigate their qualities. This has the potential to increase our overall power to understand these qualities of old sweeps and others with subtle signals.

A potential avenue of improvement in the future would be to include ancestral recombination graphs (ARGs) as an element of Flex-sweep, or to combine the Flex-sweep convolutional neural network (CNN) with another deep-learning approach to sweep inference through ARGs. It is possible, for example, to combine CNNs with recurrent neural networks (RNNs) such as the ARG-based RNN developed by Hejase et al. 2022. Such a combination would have the advantage of taking into account both current patterns of diversity and the historical patterns they resulted from, and typically shows improved performance over CNNs or RNNs alone (as demonstrated by Gheisari et al. 2021).

An important feature of any sweep detection method is robustness to non-equilibrium demographic scenarios. Flex-sweep is trained using user-specified neutral and sweep simulations, allowing it to account for variations in demographic and selective sweep parameters. Even in populations with complex demographic histories, such as the sequence of expansions and bottlenecks in the Yoruba demographic history, it has good power to detect sweeps up to 5,000 generations old. We also demonstrate its robustness to demographic misspecification as well as recombination rate heterogeneity and background selection. Despite this, in a scenario in which the true population history is completely unknown, as might be the case in a non-model species, its performance suffers when trained under a very different demographic scenario (eg. expansion) than the true history (eg. decline). In this circumstance it would be beneficial to first infer the general trend of the population history using the available data and a method such as ∂a∂i (Gutenkunst et al. 2010), PSMC (Li and Durbin 2011), MSMC (Schiffels and Durbin 2014), Relate (Speidel 2019), or StairwayPlot2 (Liu and Fu 2020), and the data needed to find sweeps, especially depth of population sequencing, will allow good demographic inference. It has good power at false positive rates of less than 1% in simulations, though this is reduced by background selection. Moreover, while this method does require knowledge of derived and ancestral alleles, it is not dependent on having an outgroup sequenced population nor admixture within the population being studied.

As expected, there is less power to detect sweeps with extremely subtle signals, including very old, weak, or incomplete sweeps and sweeps from higher amounts of standing variation. However, we test Flex-sweep under a broader parameter space than commonly used in evaluating the performance of other methods, especially with respect to sweep age (up to 0.125*4N_e_ generations) and strength (as low as *s* = 0.001, a selection strength of γ = 20 at *N_e_* = 10,000). While it has lower power to detect such weak sweeps under a very stringent false positive rate of 0.1%, its performance improves at a more relaxed false positive rate of 1% (Figure 2). In addition, the presence of population declines reduces the power to detect sweeps because bottlenecks can weaken signatures of past sweeps and are well-known to create patterns of variation that mimic selective sweeps. This makes it challenging to distinguish between a real sweep and a rapid frequency increase due to sampling noise during the bottleneck. Training Flex-sweep under a demographic model that includes population declines partially ameliorates this effect. Conversely, it is easier to detect sweeps that happened before, during, or shortly after a population expansion because the expansion decreases drift, preserving the signatures of previous sweeps. Simulations that include expansions have less variation in genetic diversity overall, and stronger differences between neutral and sweep regions, than do simulations of equilibrium demography. This is likely related to the increase in the number of haplotypes carrying the swept allele that occurs during an expansion, which maintains the signal of an old sweep longer than in an equilibrium simulation. In addition, the larger population size will limit the impact of genetic drift, so there is less variation among neutral simulations. This amplifies the differences between neutral and swept regions. These effects too are apparent in Flex-sweep’s increased power to detect sweeps under conditions of population expansion.

Flex-sweep is capable of detecting some sweeps up to 10,000 generations old in the human demographic model. This corresponds to the approximate time period of the emergence of modern humans (Schlebusch et al. 2017). As a result, there is the potential to detect adaptations that could correspond to shared traits among the ancestors of all modern humans. However, this is limited to strong, hard sweeps that are complete, or nearly so. Signals from sweeps in which the beneficial allele has gone to fixation will persist for a much longer time, as there are fewer ancestral variants remaining on the derived haplotype which will degrade signal through the action of recombination. The *hapDAF-o* statistic in particular will be particularly influential in detecting such sweeps. It is important that Flex-sweep is both able to detect these sweeps and robust to background selection because other statistics that may be able to detect old statistics (such as π) are also sensitive to background selection. We note that the CNN model used to classify the 10,000 generation old sweeps was the same Yoruba model described above, and was trained only with sweeps up to 5,000 generations old. It is possible that training with even older sweeps would increase this power.

We also compare Flex-sweep to diploS/HIC, another powerful CNN-based method that popularized deep machine-learning based sweep detection (Kern and Schrider 2018). Flex-sweep has greater power and lower false positive rates when trained and tested on data sets including only complete sweeps, which diploS/HIC is designed to classify. Flex-sweep should be a further improvement when attempting to detect sweeps genome-wide because of its ability to detect a greater diversity of sweeps with more power. However, Flex-sweep does not attempt to classify sweep type. In contrast, diploS/HIC (and its extension partialS/HIC, Xue et al. 2021) classify sweeps by hard or soft, complete or incomplete, and whether the window includes the swept locus itself or is linked to the swept locus. It would be advantageous to use these tools in succession: Flex-sweep allows first detecting diverse sweeps with high confidence, and diploS/HIC, and/or other methods (such as those of Peter et al. 2012; Akbari et al. 2018; Sugden et al. 2018; Speidel et al. 2019; Torada et al. 2019; and Caldas et al. 2022) meant to determine the properties or specific locus of a selective sweep can then be applied to the high confidence regions where sweeps have already been detected to determine the properties of these selective sweeps with greater power. In these cases, if the successor tool classifies an entire high-confidence region as neutral, we could infer that it is more likely to be a region with a sweep that the tool has difficulty detecting. If the successor tool classifies partial sections on the ends of a high confidence region as neutral, we might instead infer that it is narrowing down the location of the specific swept locus.

In contrast to previous work (Kern and Schrider 2018), we find that the order of the summary statistics in the input tensor does have a modest effect on power. An order that intersperses statistics that summarize different aspects of the genetic variation (Table 1), rather than grouping statistics by the type of data they summarize, increases power. This may be a result of the convolutional filters that span incorporating multiple types of information earlier in the process, thus increasing the information content of each layer. This is an avenue that could be further explored in the future.

While background selection reduces Flex-sweep’s ability to identify swept regions, it does not cause the CNN to generate false positives, regardless of the density of coding regions. This is congruent with findings that selective sweeps and background selection cause different patterns of genetic diversity, especially when haplotype structure is taken into account (Enard et al. 2014; Schrider 2020). In addition, we also find the surprising result that background selection causes false negatives for swept regions. This was also found by (Schrider 2020), who proposed that it may be the result of Hill-Robertson interference (Hill and Robertson 1966) and/or reduction in diversity locus-wide, including in the regions around selective sweeps that the CNN may use as distinguishing features. It may be valuable to explore the potential of using a training data set that incorporates background selection with an appropriate DFE to increase power.

On the other hand, regions of very low recombination rate do increase Flex-sweep’s false positive rate, as expected from previous work (O’Reilly et al. 2008; Ferrer-Admetlla et al. 2014; Lotterhos 2019; Booker et al. 2020). However, while regions of low recombination rate and selective sweeps both reduce local diversity, the overall patterns of genetic variation they create are different (McVean 2007). Since the CNN calculates statistics in windows of five sizes to incorporate information from a range of distances from the site under selection, the CNN is able to take advantage of these differences in overall patterns to distinguish sweeps from recombination deserts. This can be seen by increasing the confidence threshold for making a determination of whether a region contains a sweep, which reduces the false positive rate. When the recombination map of the species being analyzed can be estimated, for example through LD analyses (eg. LDhelmet, Chan et al. 2012) or machine learning (eg. ReLRNN, Adrion et al. 2020) with the same data with which sweeps are inferred, we recommend a two-step process for refining predictions in low recombination rate regions: 1. Predict sweep windows using the model trained on data generated with a distribution of recombination rates centered around the average recombination rate; 2. Retrain the model using a distribution of recombination rates centered around a lower recombination rate, then use this retrained model to predict those windows in low recombination rate regions that were previously identified as sweeps. This “clean-up” step thus removes many of the false positives caused by low recombination rates.

In addition, mispolarization of variants is especially likely to be a concern in non-model species with poorly sequenced outgroups. However, we find that low rates of mispolarization (0.1%, 1%) have little effect on classification power and false positive rates. Contrary to expectations, higher rates of mispolarization (5%, 10%) decrease FPR and increase power. We speculate that this is because many of the statistics in Flex-sweep use the frequency of shared flanking variants on ancestral vs derived backgrounds (eg. DIND and *hapDAF*) in which a high frequency of shared flanking variants on a derived background relative to the ancestral background suggests a selective sweep. Most frequently, abundant low frequency derived alleles are mispolarized as ancestral, and the corresponding high frequency true ancestral allele is mispolarized as derived. High frequency ancestral alleles can reach high frequencies when hitchhiking with a nearby selected allele, a sweep signature that we have not exploited with our current statistics. However, this hitchhiking of ancestral alleles may likely contribute to added power by being at least partially captured by statistics such as hapDAF, when these alleles are mispolarized as derived alleles. Supporting this, we see that *hapDAF-s* is one of the most important statistics influencing Flex-sweep’s classifications (see below).

Statistics vary in their importance in contributing to the model’s classifiction depending on training and testing data, based on SHAP analyses (Lundberg and Lee 2017). Both *highfreq* and *hapDAF*-s, calculated for the middle as well as the shoulders of the locus at various window sizes, make up most of the top 10 most important features for models trained and tested with simulations generated under the human demographic model, equilibrium demographic model, and demographic model with a population decline. iHS, nSL, and H12 were consistently among the least important features for these three data sets. This is likely because these statistics only have good power to detect recent sweeps, and our simulated data sets also use very old sweeps for which they have little to no power. It is important to note, however, that this does not mean that these three statistics do not contribute to the overall model, nor that they are not potentially important with other training and testing data. In particular, they likely still contribute to detecting very recent sweeps, and may be more important in data sets focusing on these.

Among other loci previously identified as being subject to a selective sweep, Flex-sweep classified a large swept region around the Duffy antigen/chemokine receptor (DARC) gene, which has a null allele that is partially responsible for malaria resistance in humans (McManus et al. 2017). This region contains three other genes that have previously been identified as candidates for positive selection in humans: high affinity immunoglobulin epsilon receptor subunit alpha (FCER1a) (Reiner et al. 2012), and two olfactory receptor genes (OR10J3 and OR10J1) (Reiner et al. 2012; Fernandes et al. 2019). Despite occurring in a region of relatively high recombination (up to 4 cM/Mbp), the length of this swept region suggests that Flex-sweep retains good power in high recombination regions, and that selection in this region was likely very strong (McManus et al. 2017; Hamid et al. 2021).

Flex-sweep also identified sweep windows in TLR5 (Toll Like Receptor 5) (Hawn et al. 2003; Abu-Maziad et al. 2010; Grossman et al. 2013), ITGAE (Integrin Subunit Alpha E) (Grossman et al. 2013; Triska et al. 2015; Ravenhall et al. 2018; Harris and DeGiorgio 2020), and APOL1 (Apolipoprotein L1) (Mizuno et al. 2010; Ko et al. 2013; Thomson et al. 2014), all previously identified as sweeps associated with infectious diseases or pathogen response by both genomic and functional studies. A selective sweep in APOL1 is of particular interest because two allele variants have been implicated in differential outcomes of kidney donation between African American donors and European American donors and it is a current target of research for improving outcomes of kidney donations and end-stage renal disease in African Americans (Freedman et al. 2016; Freedman et al. 2020). We also found evidence of sweeps at genes previously suggested to be sweep targets by genomic evidence alone, including CADM3 (de Magalhães and Matsuda 2012; Grossman et al. 2013; Sugden et al. 2018), CTNS (Grossman et al. 2013), DOCK3 (Higasa et al. 2009; Sugden et al. 2018), P2RX5 (Mizuno et al. 2010; Sugden et al. 2018), and SHPK (Barreiro et al. 2008 ; Kudaravalli et al. 2008; Grossman et al. 2013; Sugden et al. 2018), among others (Supplementary table 1).

Since it is designed to be versatile in terms of the types, strengths, and ages of sweeps it can detect, Flex-sweep will be particularly useful for scanning the genomes of non-model species for which we have few expectations for the nature of selection. Since it requires only a single species, requiring no outgroups save to polarize ancestral and derived alleles, it is also useful for species that have limited data. At present, Flex-sweep requires good genome assemblies and phased haplotypes, which is currently a limitation for non-model species. However, we are in the midst of an accelerating explosion of both available genomic data (through efforts such as the Earth Biogenome Project (Lewin et al. 2018) and the Darwin Tree of Life Project (The Darwin Tree of Life Project Consortium et al. 2022) and of long-read sequencing technologies (Amarasinghe et al. 2020; Amarasinghe et al. 2021). Together, these are lowering the financial and technical costs of sequencing and genome assembly, making genomic data of non-model species more attainable than ever. It will be possible to extend Flex-sweep to use unphased versions of the haplotype-based statistics, such as iHS (Voight et al. 2006) and nSL (Ferrer- Admetlla et al. 2014), as demonstrated and tested in Harris et al. 2018; Mughal and DeGiorgio 2018; and Klassmann and Gautier 2022.

In the future, loci identified using Flex-sweep can be examined with existing or new methods to identify the timing of these sweeps, further illuminating the history of natural selection in humans and model species, but especially of non-model species that are more challenging to analyze with existing methods. This information can be combined with data about the timing of climate, diet, and biogeographic shifts, among other events of interest, to identify important selective pressures that impact modern diversity.

## METHODS

We present Flex-sweep, a convolutional neural network-based sweep detection method, and describe its performance and application to the 1000 Genomes Project Yoruba population data (The 1000 Genomes Project Consortium et al. 2015). Flex-sweep uses six existing and five new (described below) summary statistics to quantify signals of selective sweeps (Table 1).

### Definition of New Statistics

Here, we present five new statistics designed to detect the signatures of diverse sweeps and sweeps with attenuated signals designed to complement the published statistics. This includes incomplete sweeps from standing variation and incomplete sweeps older than can be detected with existing statistics. These statistics detect the signatures of positive selection using alterations in the frequency of variants on derived vs. ancestral backgrounds. In this respect, they are inspired by the DIND statistic (Derived Intra-allelic Nucleotide Diversity, Barreiro et al. 2009) that compares π (pairwise genetic diversity) within a fixed window size around a focal variant between the ancestral and derived backgrounds. The technique of comparing derived and ancestral backgrounds with a focal variant originates with the EHH statistics (Sabeti et al. 2002), from which other statistics such as iHS (Voight et al. 2006) and nS_L_ (Ferrer-Admetlla et al. 2014) are derived.

During a selective sweep, the haplotype containing the adaptive allele increases in frequency as it hitchhikes with the adaptive allele. This increases the frequency of variants linked to the adaptive allele, thus reducing variation in the derived background relative to the ancestral background. These changes can be measured by comparing the frequencies of variants present on both the derived and the ancestral backgrounds (*hapDAF*), comparing the total number of variants on the derived and ancestral backgrounds (*Sratio*), and looking for an excess of low or high frequency alleles on the derived background (*lowfreq* and *highfreq*). Each statistic is standardized against the genome-wide distribution of focal variants with similar derived allele frequencies, divided into bins with ranges of 2%:

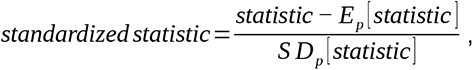

with E_p_[statistic] and SD_p_[statistic] estimated from an empirical distribution of variants with the same derived allele frequency bin (p, bins have ranges of 2% so there are 50 bins total) as the focal variant. This standardization has the effect of making each statistic comparable across focal variants regardless of focal allele frequencies.

#### hapDAF

The haplotype derived allele frequency (*hapDAF*) statistic compares the frequency of shared variants on ancestral and derived backgrounds in a fixed window size around a focal variant. It is defined as follows: A focal variant with frequency between 0.25 and 0.95 is identified, and loci are separated into those containing the derived allele and those containing the ancestral allele. Shared flanking variants that meet four criteria are identified in a window around the focal variant. These criteria are 1. the flanking variant is present on at least one derived background and at least one ancestral background; 2. the flanking variant is more common on the derived background; 3. the frequency of the flanking variant on the ancestral background is below a set threshold (see below); and 4, the total frequency of the variant on both ancestral and derived backgrounds is greater than a set threshold (see below). The frequency thresholds are determined by the goal of the scan, to identify older, incomplete sweeps (*hapDAF-o*), or more recent incomplete sweeps from standing variation (*hapDAF-s*).

The frequency of shared flanking variants is compared for shared flanking variants *i∈*{1, 2, 3, … *, k* }, following the equations

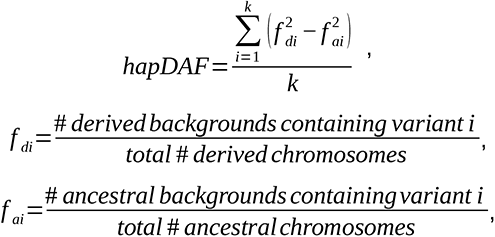

given *f _di_*> *f _ai_* (Supplementary figure 21). When there are no variants that meet the criteria (k = 0), *hapDAF* is set to 0.

*hapDAF-o*: This version of the *hapDAF* statistic is optimized to identify older, incomplete sweeps by tuning the frequency thresholds of the shared flanking variants such that *f _di_*+ *f _ai_* >0.25 and *f _ai_*< 0.25.

*hapDAF-s*: This version of the *hapDAF* statistic is optimized to identify recent, incomplete sweeps from standing variation, by tuning the frequency thresholds such that *f _di_*+ *f _ai_* >0.1 and *f _ai_*< 0.1.

#### Sratio

The segregating sites ratio (*Sratio*) compares the number of variable sites in a window of fixed size around a focal derived allele with the number of variable sites around the corresponding ancestral allele. A focal variant with a frequency between 0.25 and 0.95 is identified, and loci are separated into those containing the derived allele and those containing the ancestral allele. Variable sites are identified in a window around the focal variant, and a ratio is calculated between the number of variable sites around the ancestral focal allele and the number around the derived focal allele: 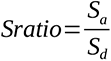 (Supplementary figure 22). *Sratio* thus detects incomplete and soft sweeps by identifying sites with fewer variable sites near derived vs. ancestral alleles.

#### freq

The two *freq* statistics look for an excess of low or high-frequency alleles on the derived background by calculating the frequencies of derived alleles in a window of fixed size around a focal variant. After defining a focal variant with a frequency between 0.25 and 0.95, all loci containing the derived allele of the focal variant (derived backgrounds) are retained. We expect little power to detect sweeps at variants with a frequency less than 0.25. Variable sites are identified in a window around the focal variant, and each site is classified as having a derived allele or an ancestral allele. The frequency of the derived allele of each variant is calculated and these are averaged over the window around the focal variant.

These two statistics complement each other by focusing on different ends of the site frequency spectrum (SFS) to quantify the excess of low and high-frequency alleles relative to expectations et al. These excesses are expected to be most pronounced after a complete, hard sweep. Restricting the calculation of the *freq* statistics to only flanking variant frequencies on the derived background approximates this expectation by treating the derived backgrounds as if they represent a complete sweep in which the derived focal allele has been fixed.

*lowfreq*: This statistic is calculated for flanking variants with the derived allele in the derived background *i∈* {1, 2, 3, … *, k* }, following the equations

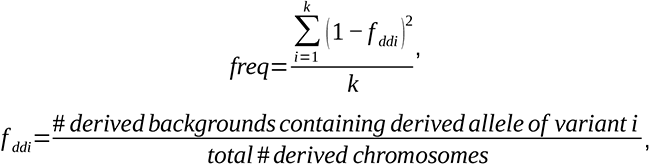

given *f_ddi_* <0.25. This statistic takes advantage of the same expected excess of rare variants as does Tajima’s D (Tajima 1989). Regions linked to a selected, derived allele that has gone to fixation (complete sweep) are expected to have an excess of rare alleles, and we approximate that by examining only loci with the derived background in which the derived focal allele is “fixed.” Because *lowfreq* is standardized by genome-wide variation data, it does not require the same theory-based standardization as Tajima’s D, which was designed at a time when genome-wide variation data was not available.

*highfreq*: This statistic is calculated for flanking variants with the derived allele in the derived background *i∈* {1, 2, 3, … *, k* }, following the equations

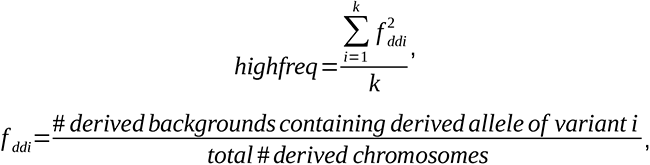

given *f _ddi_*>0.25. This statistic takes advantage of the same expected excess of high frequency variants as does Fay and Wu’s H (Fay and Wu 2000) (Supplementary figure 23). Because *highfreq* is standardized by genome-wide variation data, it does not require the same theory-based standardization as Fay & Wu’s H, which was designed at a time when genome-wide variation data was not available.

### Power of New Statistics

The power of these new statistics was tested by simulating sweeps using discoal (Kern and Schrider 2016) for a 1.2 Mb locus at several values of sweep age (scaled by 4N generations, τ ∈ {0, 0.05, 0.1, 0.15, 0.2}), selection strength (scaled by 2*Ns*, *s* ∈ {0.0025, 0.005, 0.01, 0.05}), starting allele frequency (*f*_start_ ∈ {0, 0.01, 0.05, 0.1}), and ending allele frequency (*f*_end_ ∈ {0.2, 0.3, 0.4, 0.5, 0.6, 0.7, 0.8, 0.9, 1}), see **supplementary methods**.

### Convolutional Neural Network for Sweep Detection

The use of Convolutional Neural Networks (CNNs) for sweep detection and other population genetic inference has been comprehensively described in (Kern and Schrider 2018) and reviewed in (Schrider and Kern 2018) and (Flagel et al. 2019), but we give a brief description in the **supplementary methods**.

### Simulations

Flex-sweep is trained using simulated data. While the simulations for the analyses presented were performed using discoal (Kern and Schrider 2016) and SLiM 3 (Haller and Messer 2019), training data could be generated by any simulation method capable of simulating the sweep scenarios the user wishes to include in the training data set, provided it can be converted to ms- style output. We simulated neutral and sweep data at a single 1.2 Mb locus, with the selected site at the center of the locus, under a broad parameter space of selective and demographic scenarios. One hundred chromosomes (50 diploid individuals) were sampled for each scenario.

We generated data sets for demographic equilibrium, population decline, population expansion, population bottleneck with recovery, and a complex scenario following the Yoruba population history inferred by (Speidel et al. 2019). Each data set had 10,000 neutral and 10,000 sweeps for training, and an additional 1,000 neutral and 1,000 sweep simulations for testing, with mutation and recombination rates as described in Supplementary table 2. Mutation and recombination rates for the human demographic simulations were based on (Speidel et al. 2019). The starting and ending frequencies for the swept allele and the selection strength were also drawn from the distributions described in Supplementary table 2. In the demographic equilibrium simulation set, the sweep timing was drawn from the distribution as described in Supplementary table 2 while in the simulations with population size changes, the sweep timing was drawn from the distributions described in Supplementary table 3. Simulations were also conducted to vary the time of sampling after the decline, expansion, or recovery, as well as the selection timing with respect to the demographic event(s) (before or during). To test for the impacts of background selection and recombination rate heterogeneity, we also simulated data sets with 100 neutral and 100 sweep simulations in SLiM 3 (Haller and Messer 2019) under scenarios with recombination rate heterogeneity mimicking the deCODE map (Halldorsson et al. 2019) and background selection (DFE ∼ γ(mean = -0.030, shape = 0.206), Boyko et al. 2008). See supplementary methods for more detail.

### Feature Vectors

Feature vectors represent the summarized variation at each simulated locus, with one vector per simulation. We divide each simulated 1.2 Mb locus into a series of 21 center points spaced at 10kb intervals, each with nested windows of five different sizes ({50kb, 100kb, 200kb, 500kb, 1000kb}) around the center point. We then calculate 11 summary statistics to summarize the characteristics of the sample with respect to site frequency spectrum (SFS), haplotype structure, and diversity on the derived background (Table 1) in each of the five nested windows, and normalize against the genome-wide distribution of focal variants with similar derived allele frequencies (as described in Voight et al. 2006) using the neutral simulations, with the exception of *HAF* and *H12* which take a single value across the entire locus. See **supplementary methods** for more detail.

### CNN Structure and Testing

The characteristics of the feature vector, such as its variance among and between features, can have a significant effect on the ultimate performance of the CNN. We tested multiple preprocessing strategies to eplore the impact of different types of feature vector rescaling and standardization. See **supplementary methods** for more detail.

To develop the final model, we tested model architectures using the simulations described above with a variety of combinations of hyperparameters. See **supplementary methods** for more detail. We trained each model using sets of simulations as described above and tested them with balanced data sets of 1,000 neutral and 1,000 sweep simulations simulated under the same conditions (except for simulations for recombination rate heterogeneity and background selection, for which only 100 neutral and 100 sweep simulations were generated due to computational limitations). Models were tested under conditions of correct demographic specification (same demographic model for training and testing data), demographic misspecification (model tested with data generated under a different demographic scenario than it was trained on), and mixed demographic specification (model trained on data generated from a combination of demographic scenarios). We also examined the effect of misspecification of the timing of the sweep relative to the demographic event by testing models with sweeps occurring during or after a demographic event. Models were trained with 20% of the data held back for in-model validation. Once trained, the model was used to classify data from each testing data set. See **supplementary methods** for evaluation details.

All CNN code was implemented in Python (3.6) using Tensorflow 2.4 and Keras 2.1.3. A singularity container and two pre-trained models (equilibrium and Yoruba) will be available at Zenodo, DOI 10.5281/zenodo.7860595.

### Training set size

The size of the training data set can have a strong effect on CNN performance, but there is a trade-off with training time, so the marginal benefit of a larger training data set size may not be worth the increased cost of training time and computational resources. We compared the method’s performance when trained on data sets of four different sizes, from 2,000 neutral and 2,000 sweep simulations to 20,000 of each. The false positive rate continues to improve to 20,000 simulations, but the biggest improvement is between 2,000 and 5,000 simulations.

### Validation

Models were tested and compared, as described above, by training on a data set and then used to predict the held back test data sets, including those with recombination rate heterogeneity, background selection, and the human demographic model. ROC curves and AUC, false positive rates (FPR), accuracy, and precision were compared to determine the best network architecture. We explored a number of additional factors that could influence the model predictions and accuracy results, including the power added by the new statistics, the impact of the order of the statistics in the feature vector, the impact of mispolarization of alleles, the impact of extremely misspecified demographic models, and the impact of simulation parameters such as time since sweep, starting and ending allele frequencies, sweep strength, mutation rate, and especially recombination rate. We also investigated whether Flex-sweep is able to classify the types of sweeps without losing power and compared the success of Flex-sweep to diploS/HIC, one of the state-of-the-art machine learning methods designed to detect selective sweeps (Kern and Schrider 2018), see **supplementary methods** for more detail.

To explore how each statistic contributes to Flex-sweep classification, we used Deep SHAP (SHapley Additive exPlanations) to interpret the contribution of each feature (a statistic calculated at a region in the locus over a specific window size) (Lundberg and Lee 2017). See **supplementary methods** for more detail. We calculated SHAP values (Lundberg and Lee 2017) for the equilibrium-trained, YRI demography-trained, and decline-trained models and interpreted the results in their corresponding testing data following two approaches: 1. clustering features by similarity, and 2. averaging the importance across each statistic at small (50kb- 100kb), medium (200kb-500kb) and large (1Mb) window sizes in the middle (the 7 windows in the center of the locus) and shoulders (the leftmost and rightmost 7 windows of the locus) of each region.

### Application

We applied our method to data from the Yoruba population data in the 1000 Genomes Project (The 1000 Genomes Project Consortium et al. 2015). We divided each autosome into 1.2 Mb overlapping windows with a step size of 10kb and calculated a feature vector for each window using the deCODE recombination map (Halldorsson et al. 2019). We used Flex-sweep, trained on the Yoruba demographic model, to classify each window as neutral or containing a sweep. Furthermore, swept regions spanning multiple windows were identified as streaks of windows classified as sweeps. We developed a false discovery rate (FDR) metric to demonstrate its actual performance in this real-world data set by comparing the proportion of windows classified as sweeps in the Yoruba data to the false positive rate from the validation tests using the Yoruba demographic model. See **supplementary methods** for more detail.

To understand the patterns of selective sweeps across the genome, we calculated the proportion of swept regions localized within genes, as well as the distances of swept regions from transcription start and sites, as well as 5’ and 3’ UTR start and end sites and tested for statistical significance using a randomization procedure described in the **supplementary methods**. To understand the relationship of sweeps to virus-interacting proteins (VIPs) specifically, we compared the number of VIPs under selection to the number of control non-VIPs under selection, as described in (Enard and Petrov 2020), notably using the same controls for potential confounding factors. A gene (VIP or non-VIP) was considered to be subject to a sweep if the genomic center of the gene (as calculated by the midpoint of the most 5’ start and the most 3’ end) was covered by at least one window classified as a sweep.

### Data and Code Availability

Flex-sweep is available at https://github.com/lauterbur/Flex-sweep, and a singularity container with all dependencies, as well as two pre-trained Flex-sweep models for the equilibrium and Yoruba demographics described here are also provided at Zenodo (DOI 10.5281/zenodo.7860595). Code to replicate simulations and analyses presented here, and for the five new statistics presented, is also available at the above repository.

## Supporting information

Supplementary methods and results

## Acknowledgments

M. Elise Lauterbur is supported by an NSF Postdoctoral Research Fellowship #2010884. David Enard is supported by NIH NIGMS MIRA R35GM142677. This material is based upon High Performance Computing (HPC) resources supported by the University of Arizona TRIF, UITS, and Research, Innovation, and Impact (RII) and maintained by the UArizona Research Technologies department. We thank Dan Schrider, Parul Johri, the editor, and two anonymous reviewers for valuable comments, advice, and discussions.

