## Supplementary figures for "Versatile detection of diverse selective sweeps with Flex-sweep"

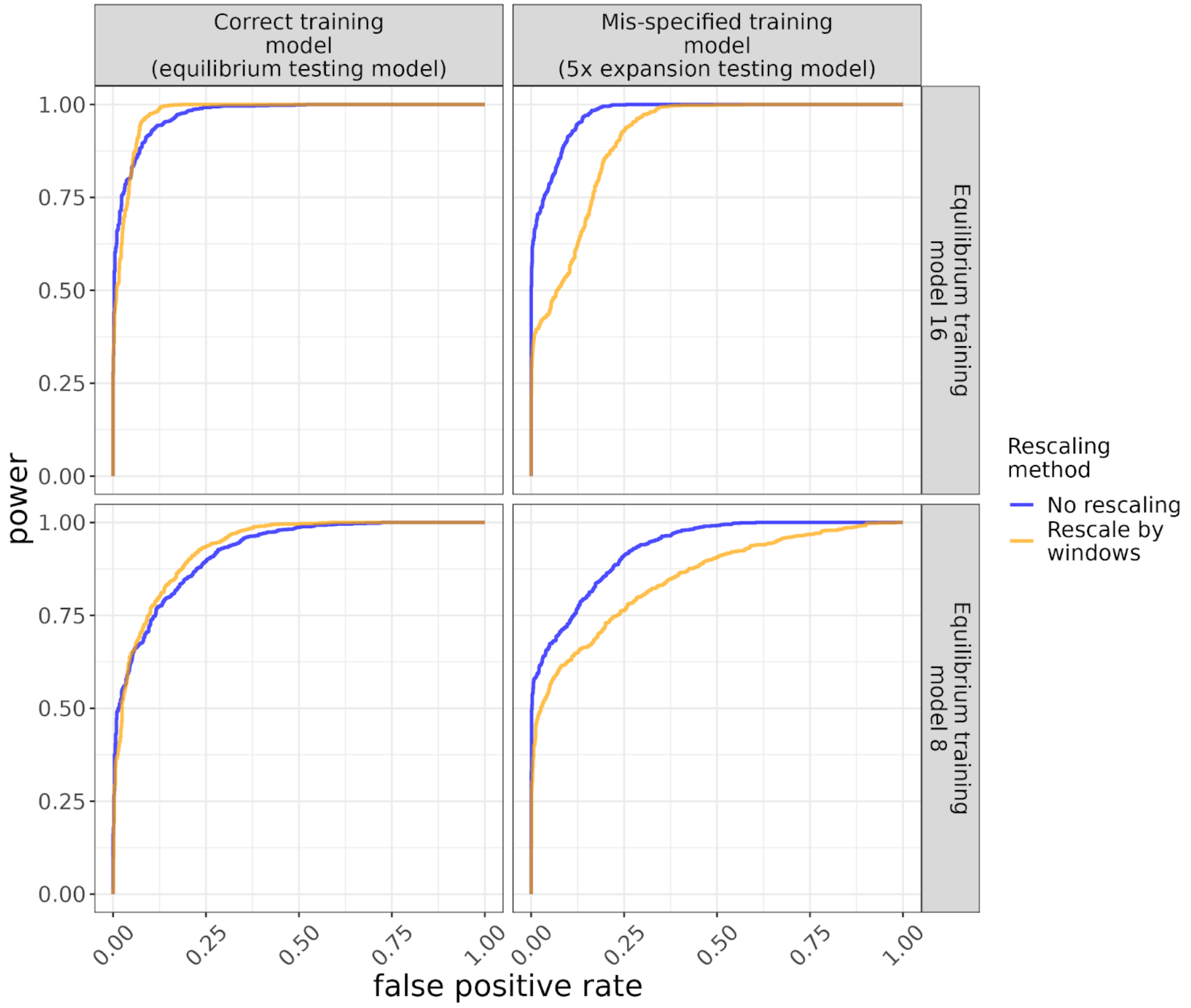


**Supplementary figure 1** ROC curves for two rescaling strategies (no rescaling vs rescaling by windows) applied to feature vectors before training a model with a correctly specified equilibrium demographic scenario (left) or a model with a mis-specified equilibrium demographic scenario predicting data simulated with a 5x expansion (right). The top row uses the model architecture implemented in Flex-sweep, the bottom row uses an alternative tested model architecture.

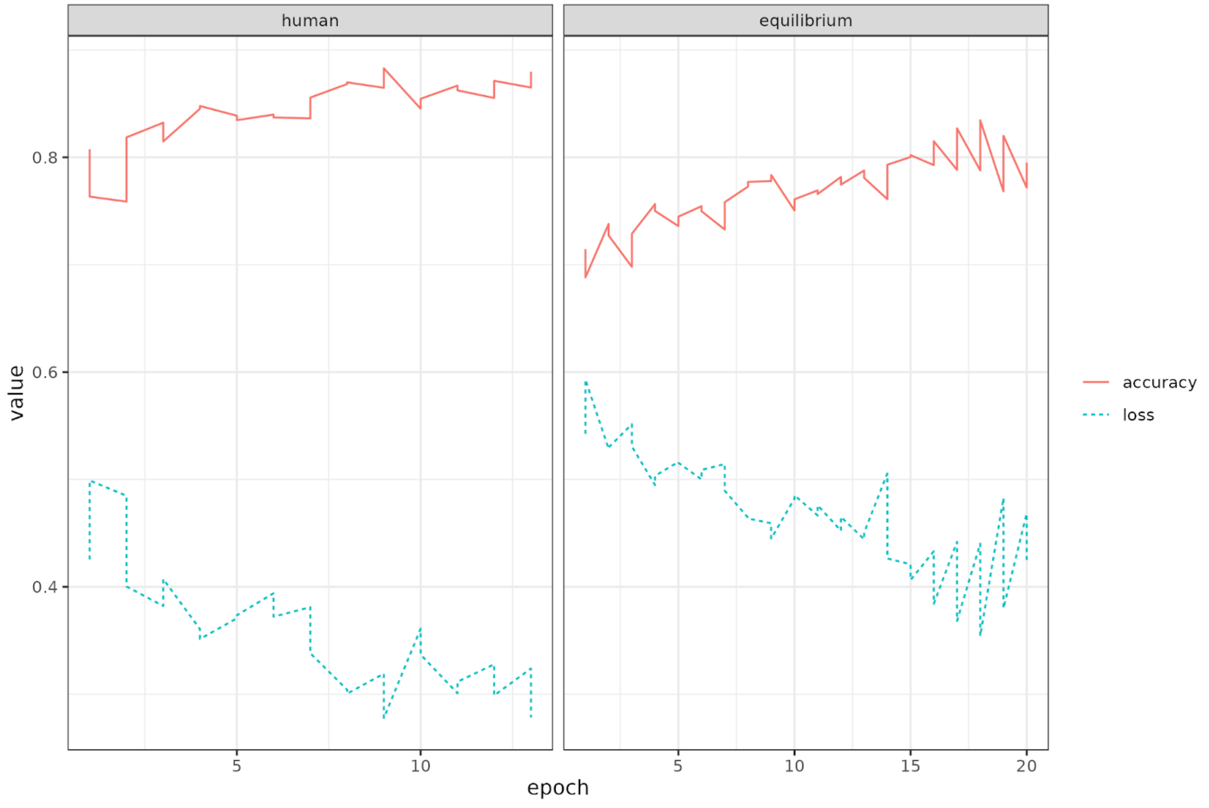


**Supplementary figure 2** Accuracy and loss over the training epochs of models trained on human demography (left) and equilibrium demography (right) showing overall increase in training accuracy and decrease in training loss over the course of the training periods.

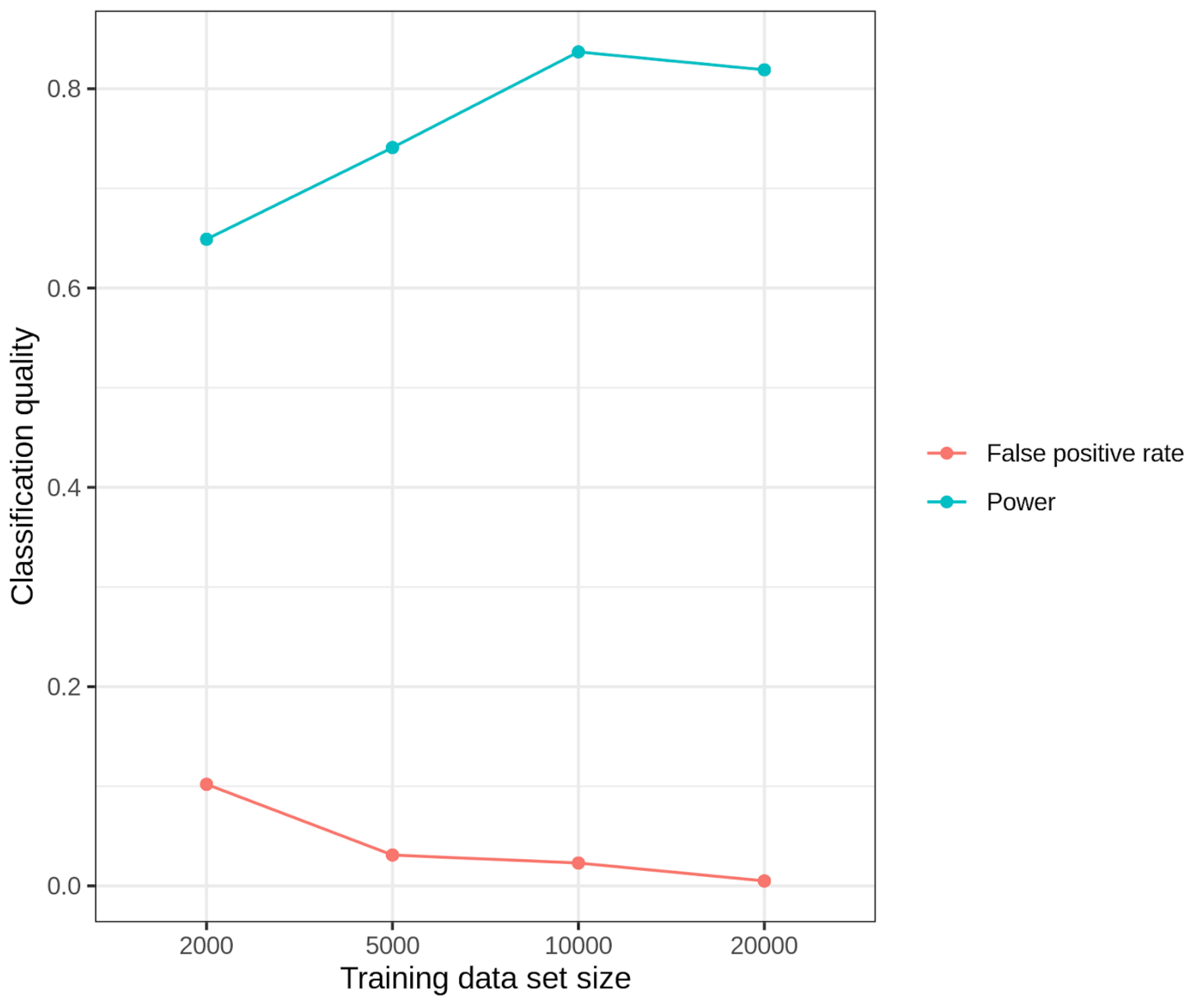


**Supplementary figure 3** The false positive rate (pink) and power (blue) of the network depending on the number each of neutral and sweep simulations in the training data set. False positive rate increases through 20,000 simulations of each, but power levels off after 10,000.

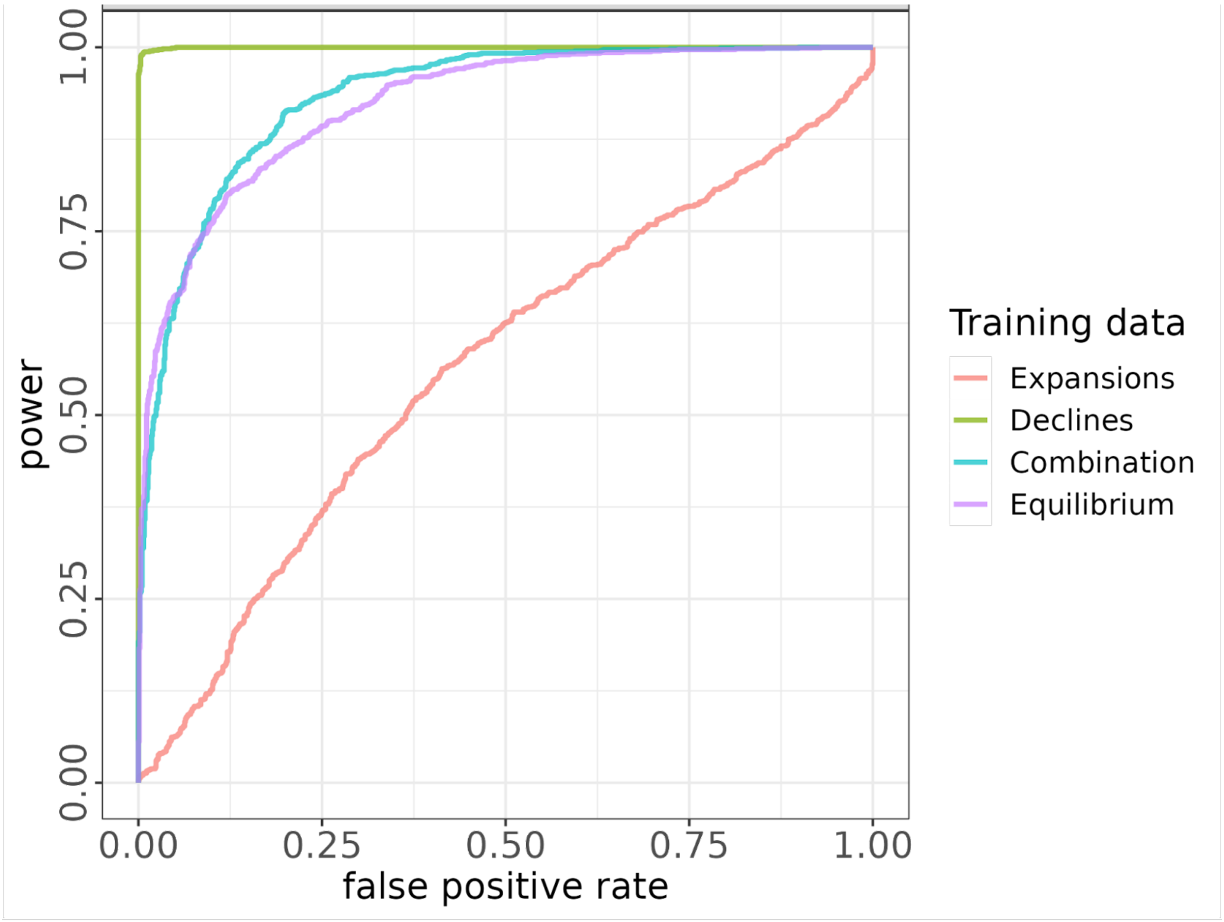


**Supplementary Figure 4.** ROC curves comparing the effect of the training data on the predictive power of data simulated under a 5x bottleneck.

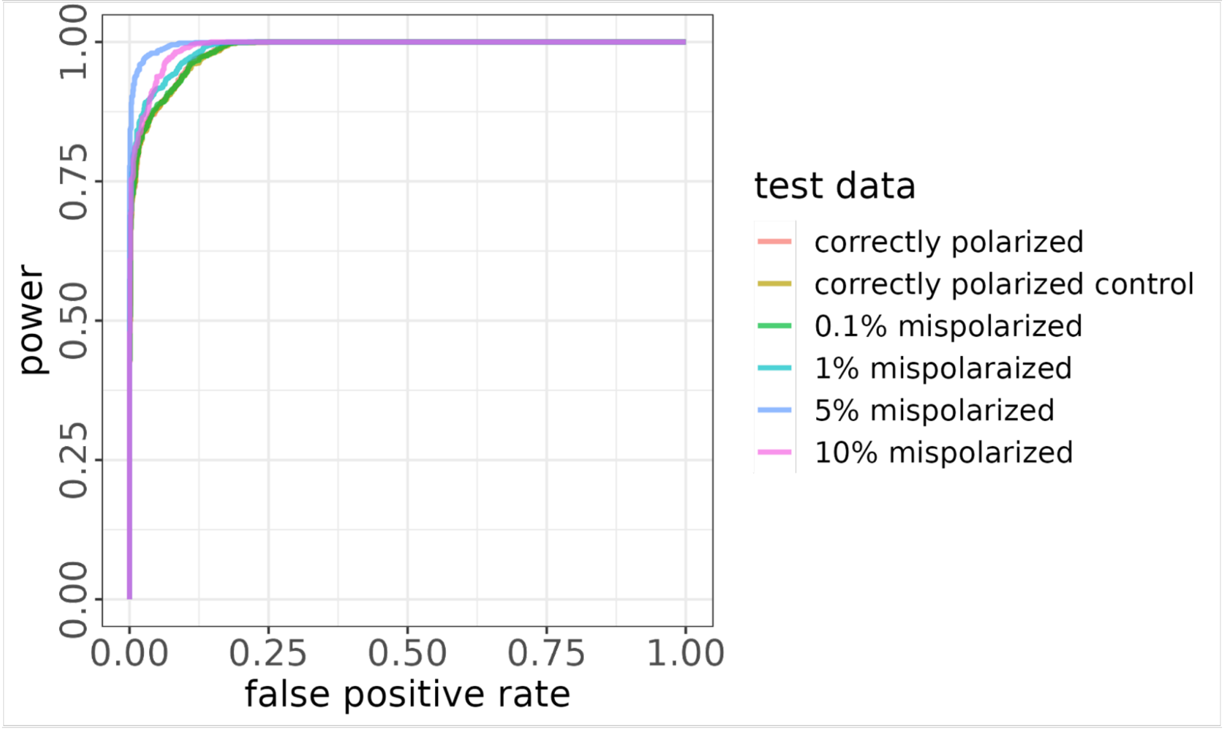


**Supplementary Figure 5** ROC curves comparing mispolarized data sets using the equilibrium demography training and test data. Correctly polarized test data is the original test data set, with each mispolarization test data set created by switching randomly chosen X% of variants from derived to ancestral states and vice versa. The correctly polarized control data set was produced by running the mispolarization script with a mispolarization rate of 0%.

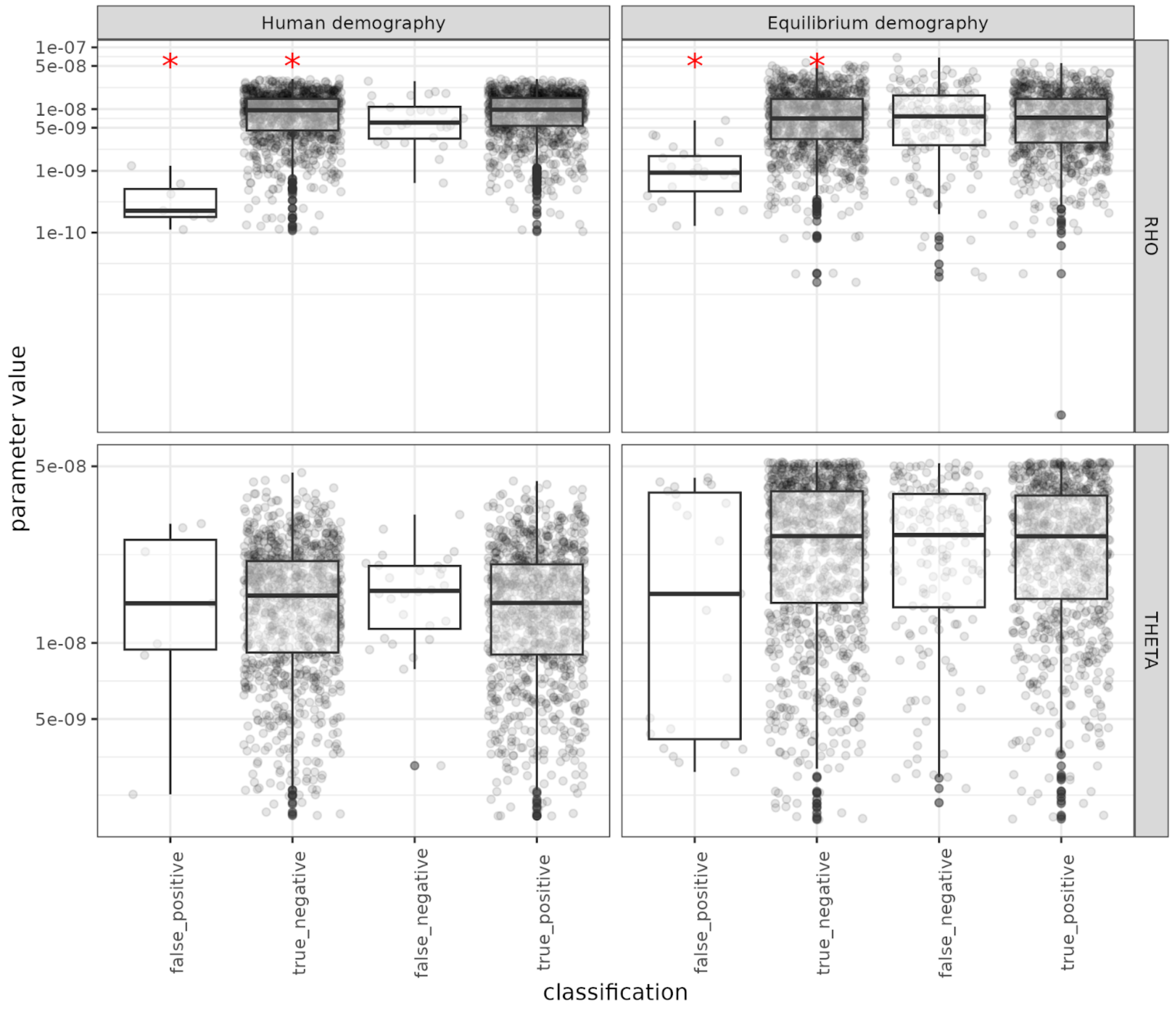


**Supplemental Figure 6** Association between simulation parameter values for recombination rate (RHO, top) and mutation rate (THETA, bottom) and false positive and false negative rates in the human demographic model and equilibrium demographic model scenarios. Boxes indicate the 25% - 75% quantiles, grey points represent the parameter value and classification of each simulation, and red asterisks indicate true/false categories that had significantly different parameter values (t-testss, α = 0.05).

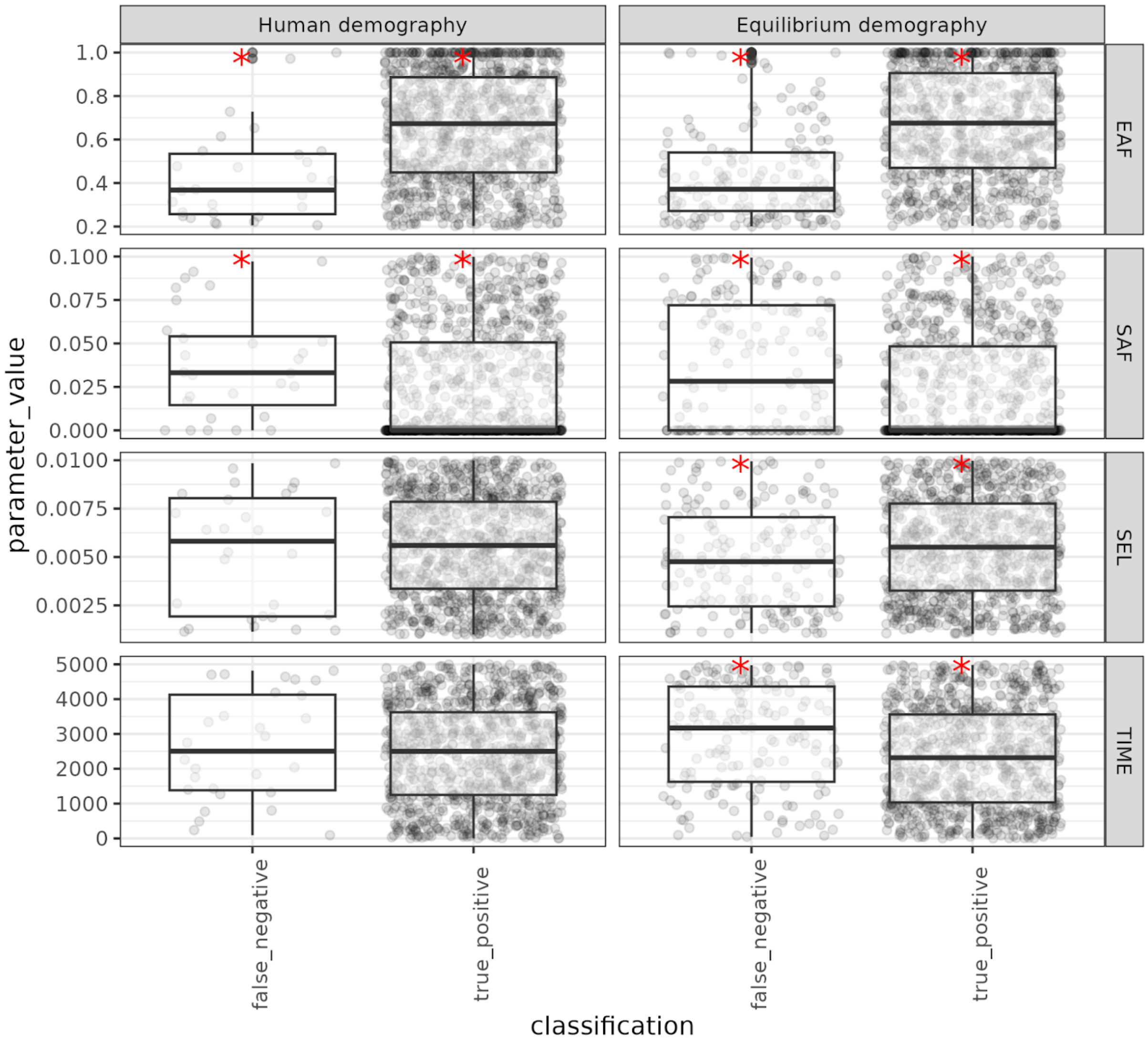


**Supplemental Figure 7** Association between simulation parameter values for ending allele frequency (EAF), starting allele frequency (SAF), selection strength (SEL), and sweep time (TIME) and false positive and true negative rates in the human demographic model and equilibrium demographic model scenarios. Boxes indicate the 25% - 75% quantiles, grey points represent the parameter value and classification of each simulation, and red asterisks indicate true/false categories that had significantly different parameter values (t-tests, α = 0.05).

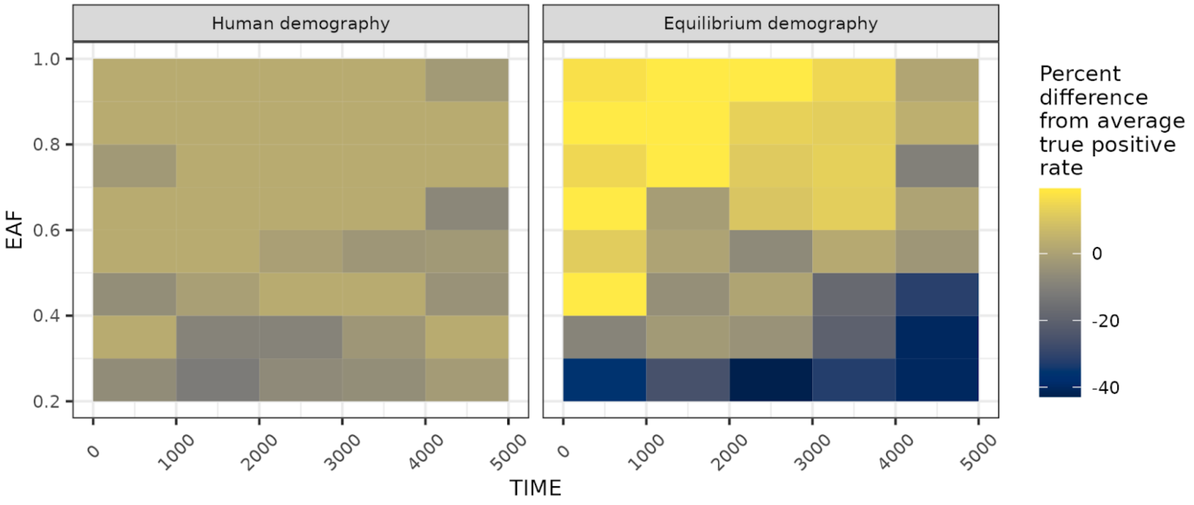


**Supplemental Figure 8** Relative difference in true positive rate at different combinations of sweep age (TIME in generations) and completeness (EAF, ending allele frequency) from the overall true positive rate for the entire data set.

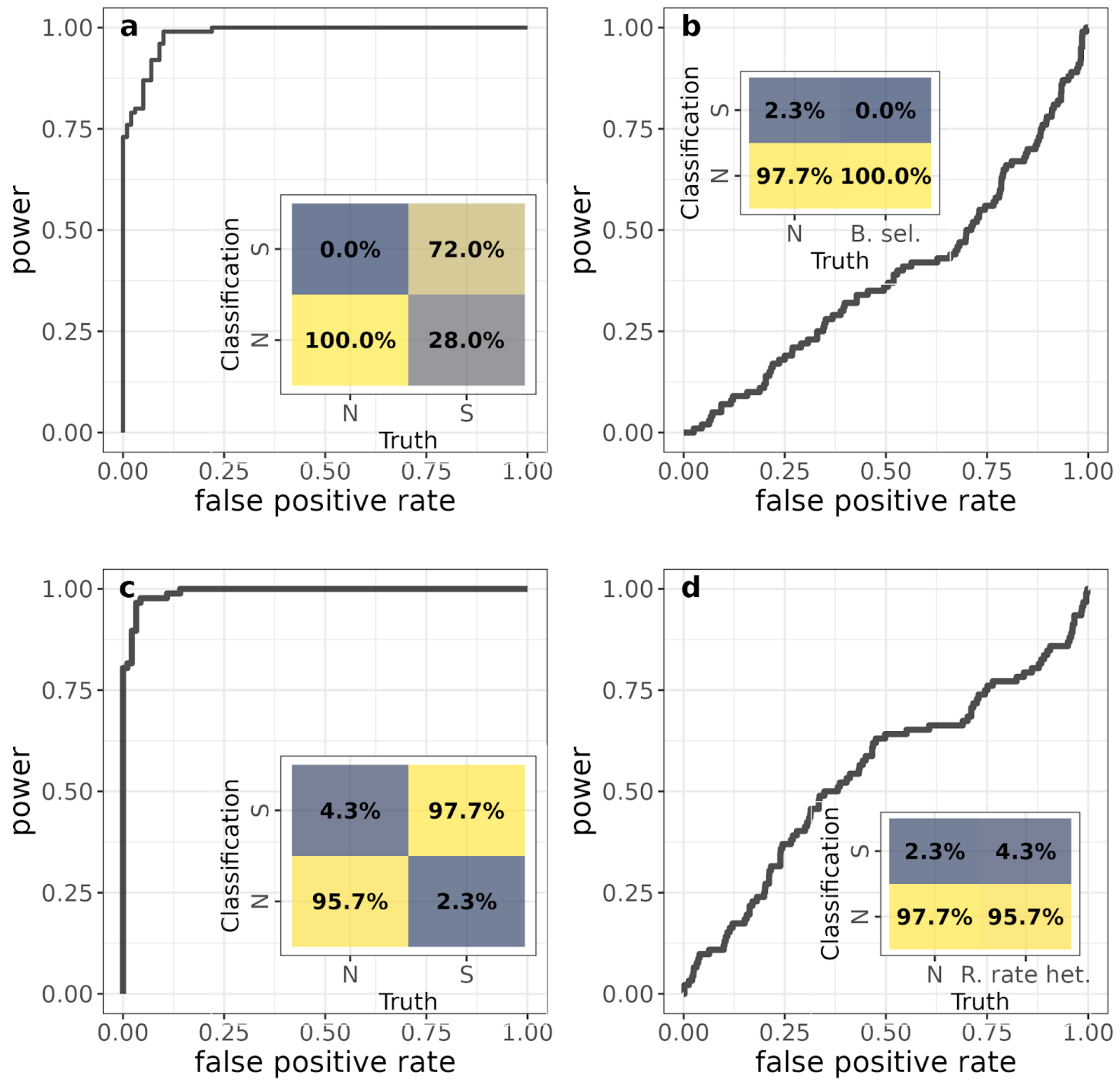


**Supplementary Figure 9** (a) Power to detect sweeps in the presence of background selection ( DFE ~ $\gamma$(mean = -0.030, shape = 0.206)) and (b) the effect of background selection on false positive rate by using the CNN to classify neutral simulations with and without background selection; c) The power to detect sweeps with recombination rate heterogeneity mimicking human chromosome 1 and (d) and the effect on false positive rate by using the CNN to classify neutral simulations with and without recombination rate heterogeneity. The diagonal ROC curves in (b) and (d) show that the CNN does not classify neutral simulations with background selection (b) or recombination rate heterogeneity (d) as sweeps more often that it does neutral simulations without background selection or recombination rate heterogeneity.

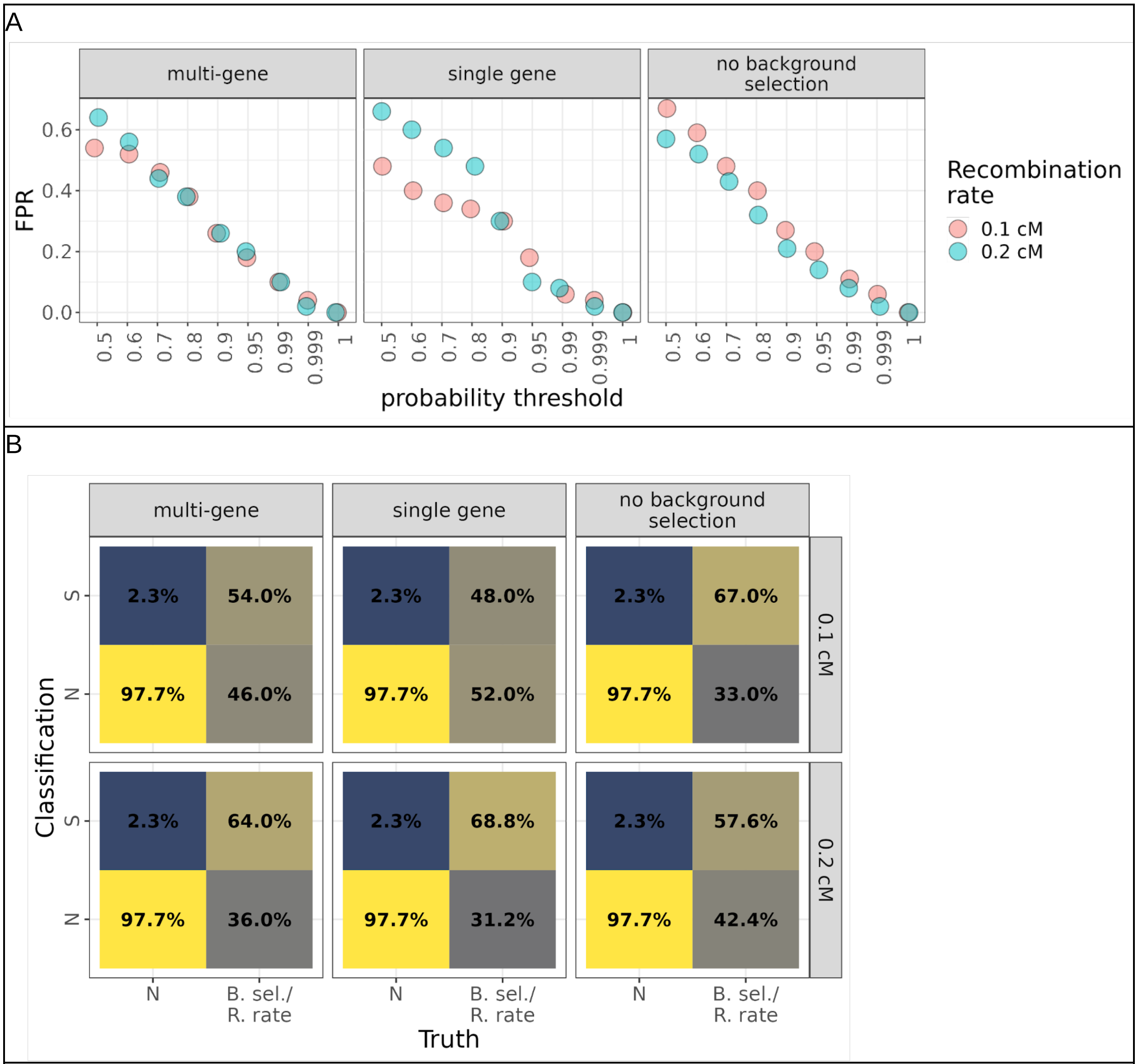


**Supplementary Figure 10** The effects of recombination rate and background selection on classification accuracy, a. the relationship of the probability threshold on false positive rate at different recombination rates, with and without background selection and b. confusion matrices showing the effect of recombination rate and exonic structure with background selection at different recombination rates vs variation in recombination rate only on false positive rate by using the CNN to classify neutral simulations with and without background selection.

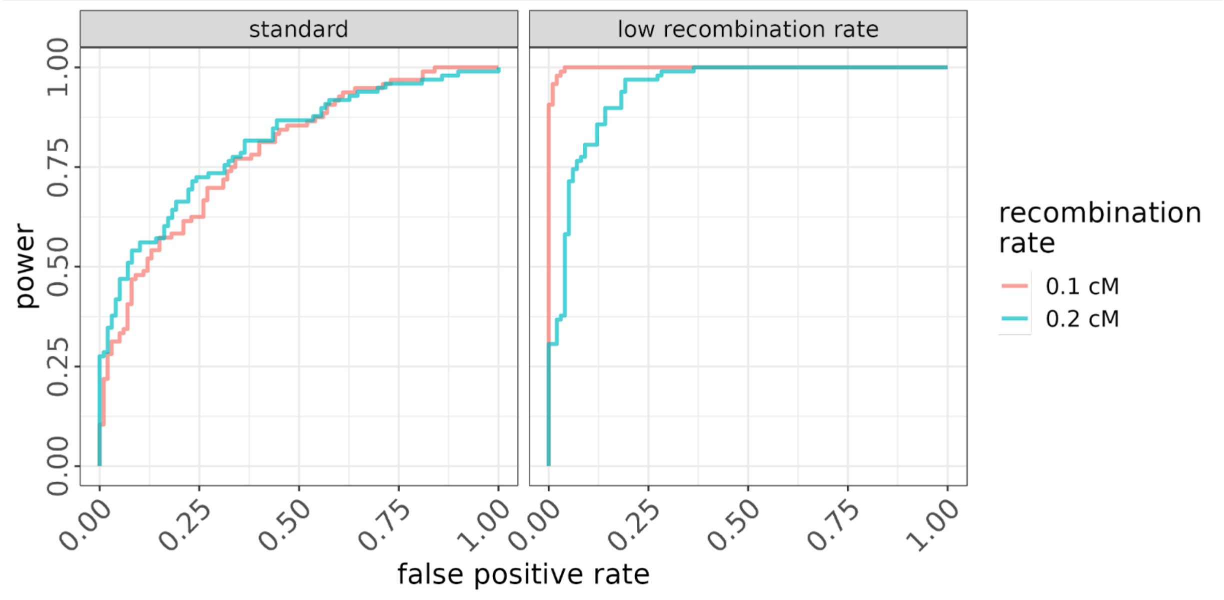


**Supplementary Figure 11** ROC curves showing power and false positive rate at different recombination rates of the model trained under the equilibrium demographic scenario with the standard distribution of recombination rates vs trained under the equilibrium demographic scenario with the distribution of recombination rates an order of magnitude lower, to predict simulations with recombination rates of 0.1 and 0.2 cM.

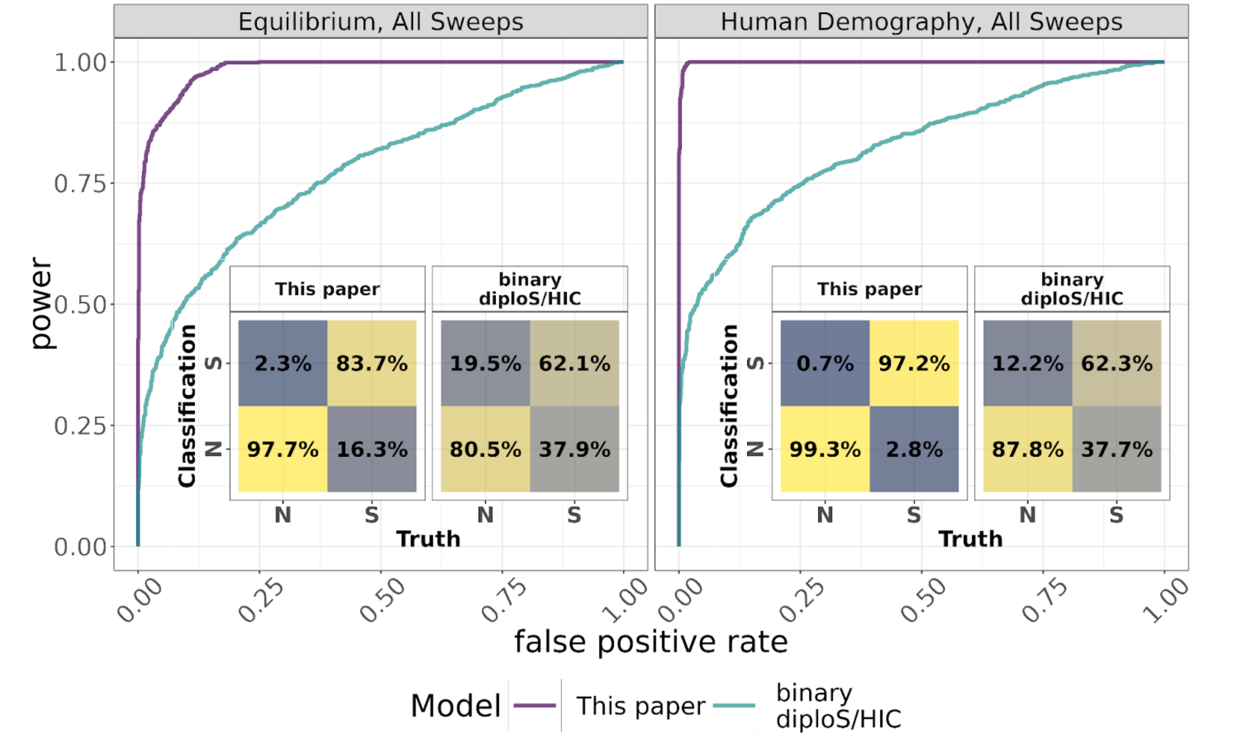


**Supplementary Figure 12** Power and false positive rates comparing Flex-sweep and diploS/HIC trained and tested on data under an equilibrium demography including incomplete and complete sweeps (left), and the human demography data set including incomplete and complete sweeps (right).

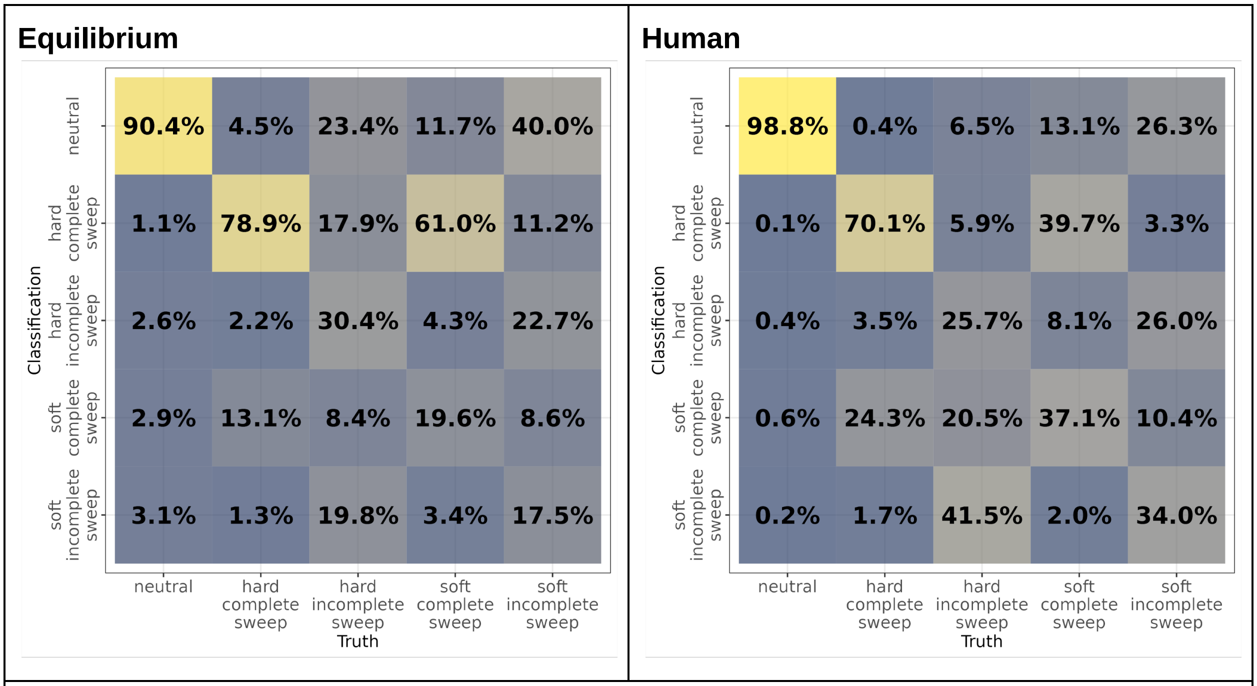


**Supplementary Figure 13** Confusion matrices describing the power to classify sweeps divided into four types: hard complete, hard incomplete, soft complete, and soft incomplete, under the equilibrium demographic scenario (left) and the YRI demographic model (right). The false positive rate (identifying a neutral region as any kind of sweep) is somewhat increased, and sweep types are frequently misclassified.

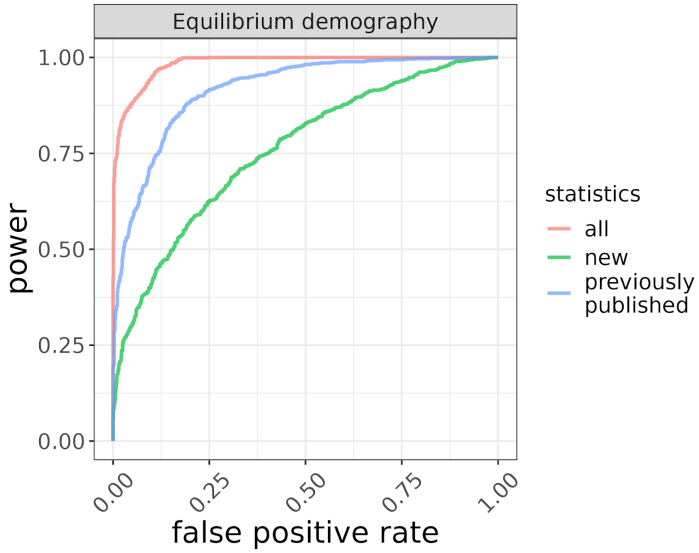


**Supplementary Figure 14** ROC curves comparing the CNN when using only the previously published summary statistics (blue line), the summary statistics described in this paper (green line), or all together (pink line). The full combination of 11 statistics has greater power to classify sweeps than either sub-group.

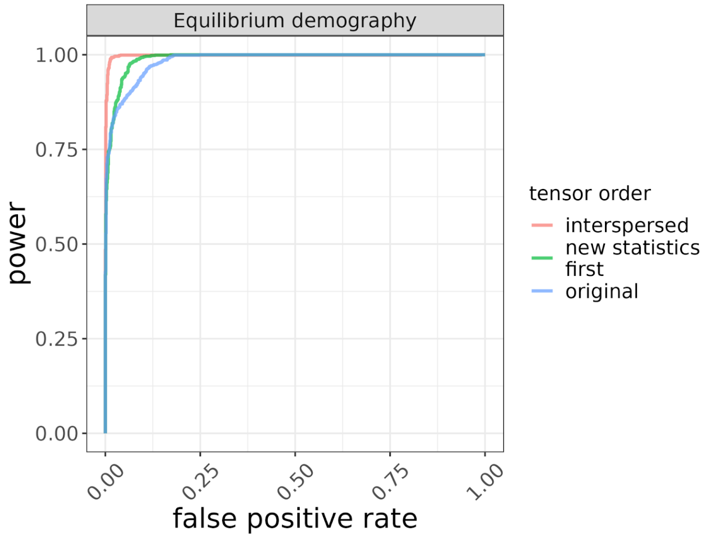


**Supplemental Figure 15**. The effect of the order of the statistics in the input tensor under the equilibrium demographic scenario. The original order is shown in green, grouping the new statistics in pink, and randomized in blue.

A

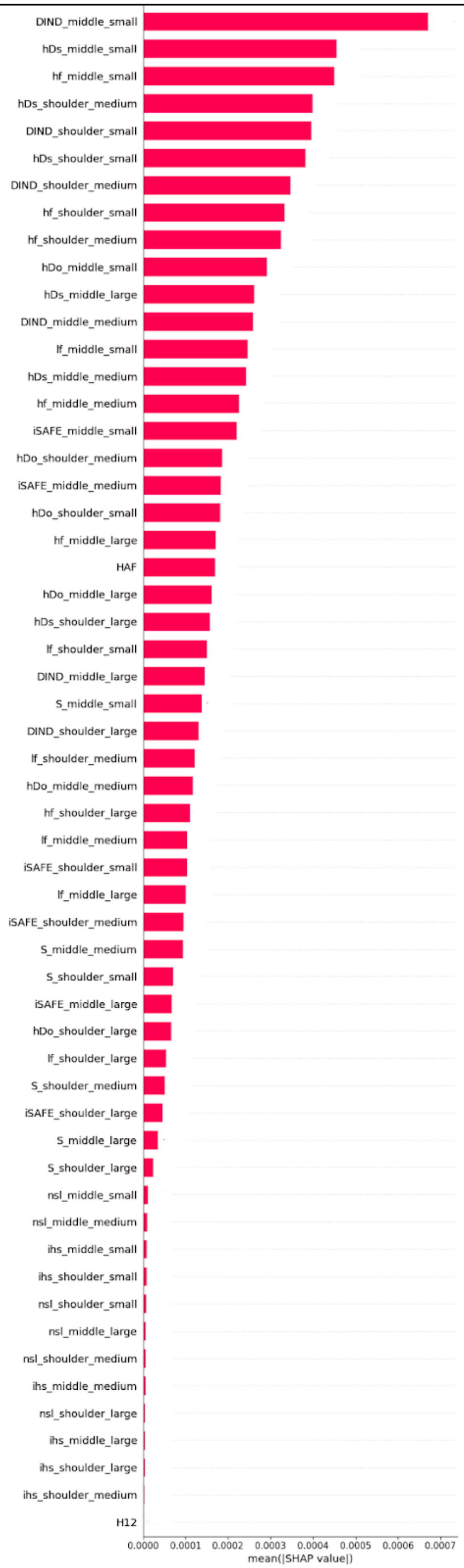

DIND_middle_small
hDs_middle_small
hf_middle_small
hDs_shoulder_medium
DIND_shoulder_small
hDs_shoulder_small
DIND_shoulder_medium
hf_shoulder_small
hf_shoulder_medium
hDo_middle_small
hDs_middle_large
DIND_middle_medium
lf_middle_small
hDs_middle_medium
hf_middle_medium
iSAFE_middle_small
hDo_shoulder_medium
iSAFE_middle_medium
hDo_shoulder_small
hf_middle_large
HAF
hDo_middle_large
hDs_shoulder_large
lf_shoulder_small
DIND_middle_large
S_middle_small
DIND_shoulder_large
lf_shoulder_medium
hDo_middle_medium
hf_shoulder_large
lf_middle_medium
iSAFE_shoulder_small
lf_middle_large
iSAFE_shoulder_medium
S_middle_medium
S_shoulder_small
iSAFE_middle_large
hDo_shoulder_large
lf_shoulder_large
S_shoulder_medium
iSAFE_shoulder_large
S_middle_large
S_shoulder_large
nsl_middle_small
nsl_middle_medium
ihs_middle_small
ihs_shoulder_small
nsl_shoulder_small
nsl_middle_large
nsl_shoulder_medium
ihs_middle_medium
nsl_shoulder_large
ihs_middle_large
ihs_shoulder_large
ihs_shoulder_medium
H12
0.0000
0.0001
0.0002
0.0003
0.0004
0.0005
0.0006
0.0007
mean(|SHAP value|)

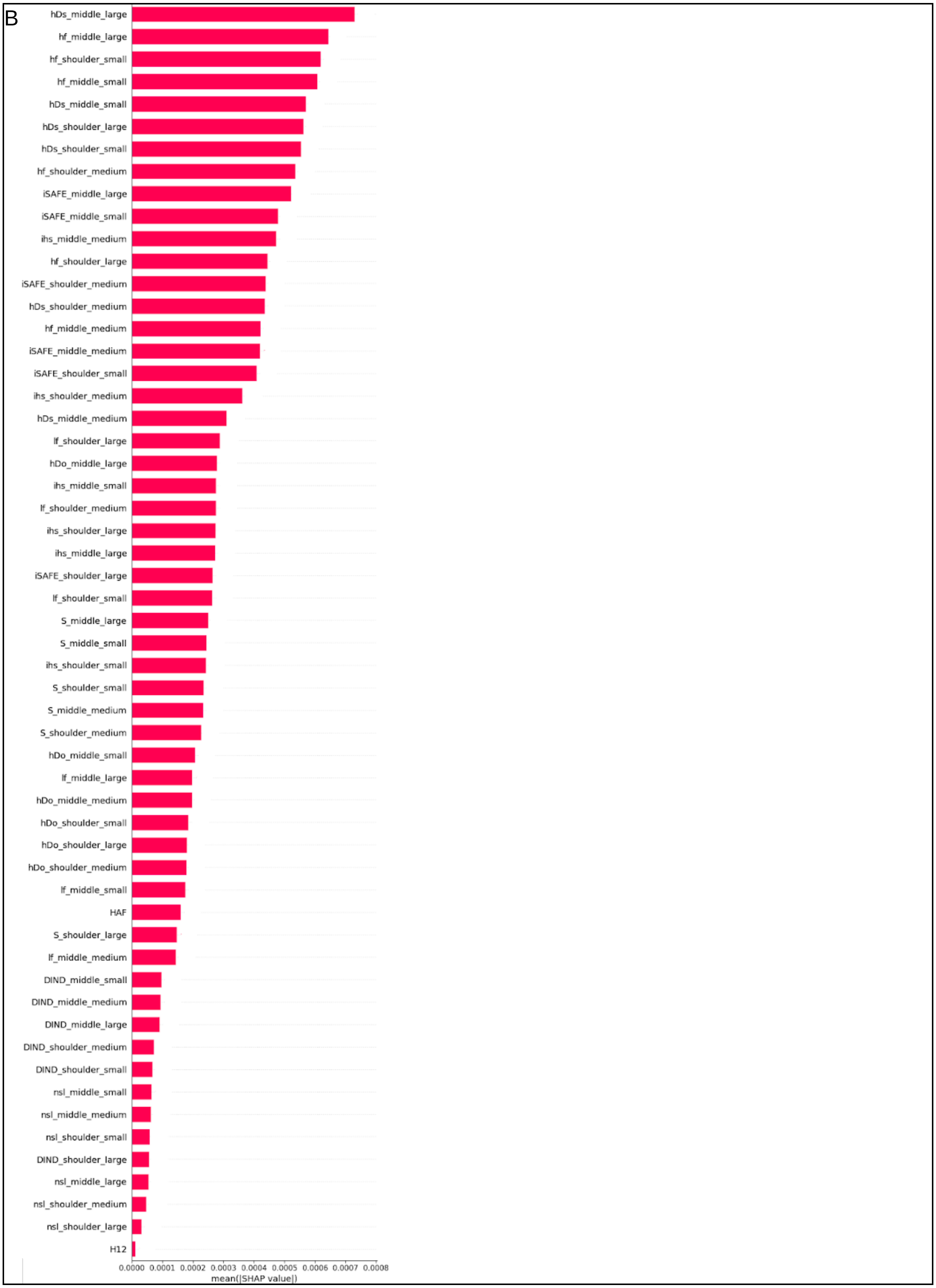
B
hDs_middle_large
hf_middle_large
hf_shoulder_small
hf_middle_small
hDs_middle_small
hDs_shoulder_large
hDs_shoulder_small
hf_shoulder_medium
iSAFE_middle_large
iSAFE_middle_small
ihs_middle_medium
hf_shoulder_large
iSAFE_shoulder_medium
hDs_shoulder_medium
hf_middle_medium
iSAFE_middle_medium
iSAFE_shoulder_small
ihs_shoulder_medium
hDs_middle_medium
lf_shoulder_large
hDo_middle_large
ihs_middle_small
lf_shoulder_medium
ihs_shoulder_large
ihs_middle_large
iSAFE_shoulder_large
lf_shoulder_small
S_middle_large
S_middle_small
ihs_shoulder_small
S_shoulder_small
S_middle_medium
S_shoulder_medium
hDo_middle_small
lf_middle_large
hDo_middle_medium
hDo_shoulder_small
hDo_shoulder_large
hDo_shoulder_medium
lf_middle_small
HAF
S_shoulder_large
lf_middle_medium
DIND_middle_small
DIND_middle_medium
DIND_middle_large
DIND_shoulder_medium
DIND_shoulder_small
nsl_middle_small
nsl_middle_medium
nsl_shoulder_small
DIND_shoulder_large
nsl_middle_large
nsl_shoulder_medium
nsl_shoulder_large
H12
0.0000 0.0001 0.0002 0.0003 0.0004 0.0005 0.0006 0.0007 0.0008
mean(|SHAP value|)

C

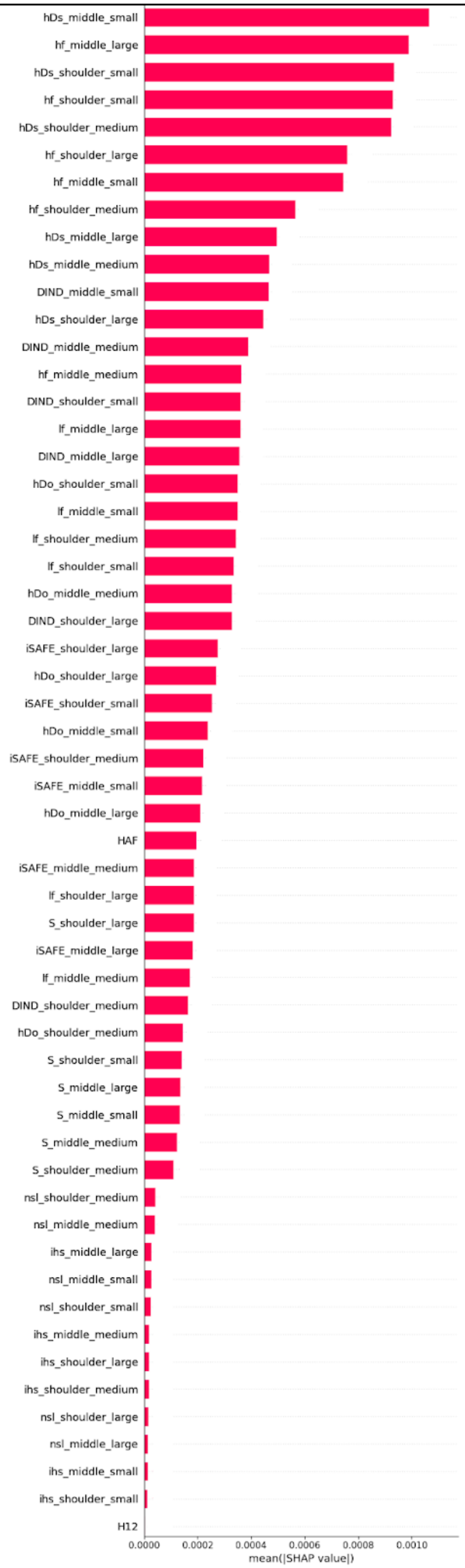


**Supplementary Figure 16.** Mean of the absolute SHAP values of each statistic grouped by window size (small - 50kb, 100kb; medium - 200kb, 500kb; large - 1Mb) in the middle (the 7 windows in the center of the locus) and shoulders (the leftmost 7 and rightmost 7 windows of the locus). Calculated for training and testing data of A. the human demographic model, B. the model with demographic declines, or C. the equilibrium demographic model.

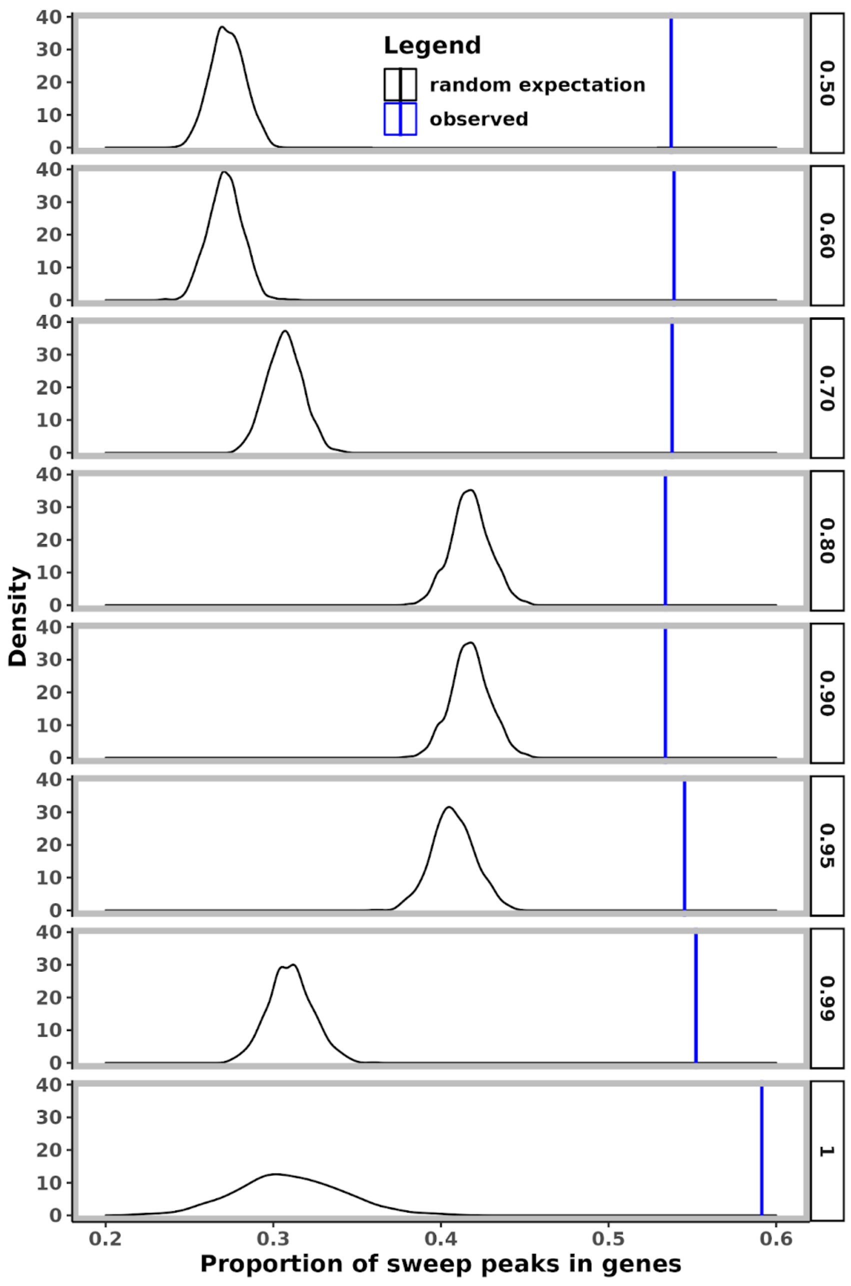


**Supplemental Figure 17** The observed proportion of selective sweeps within genes (blue line) compared to the distribution expected by randomization (black line) at different classification confidence thresholds (rows).

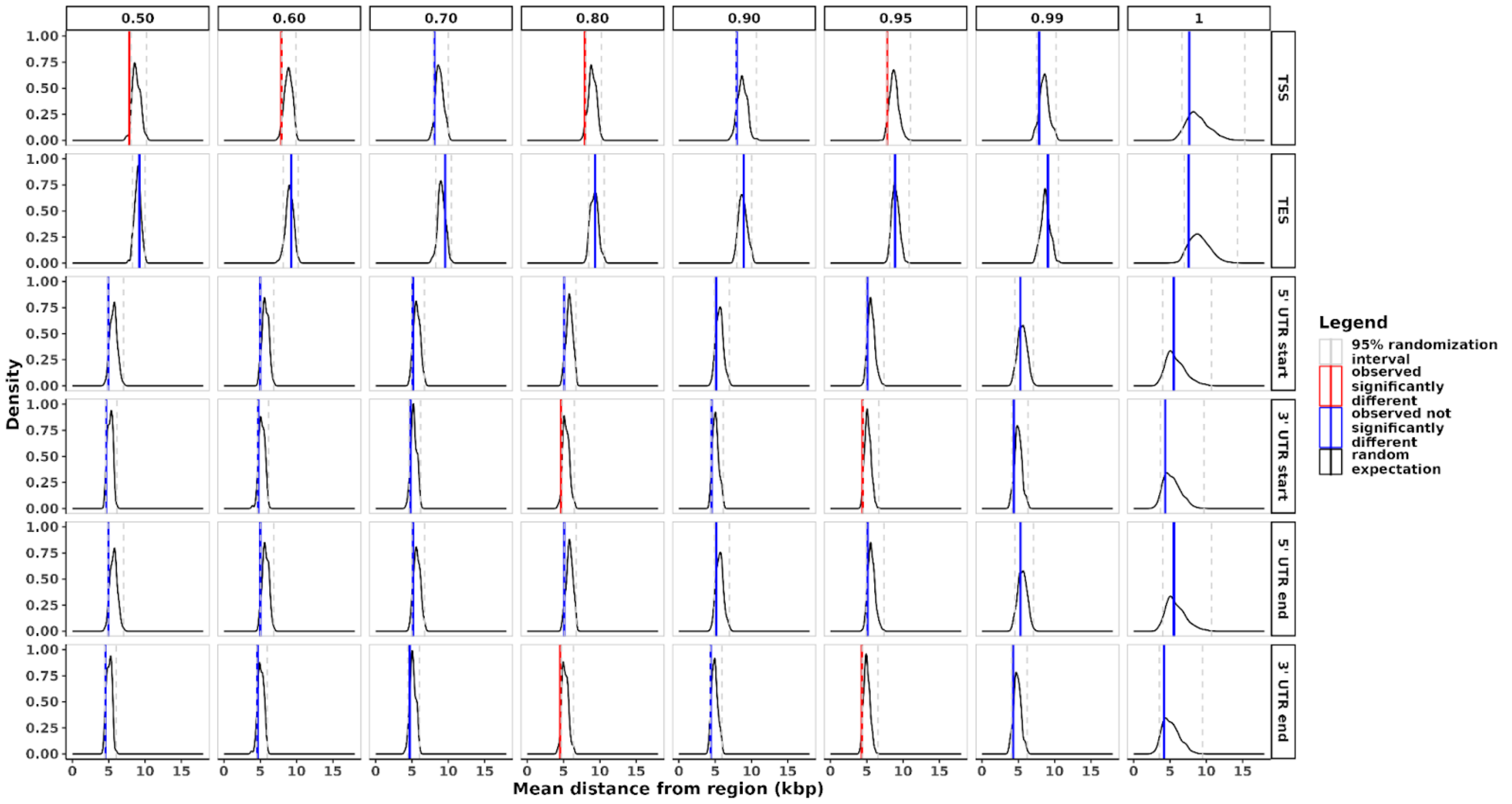


**Supplemental Figure 18** The observed mean distance (red and blue lines) of the peak of each swept region to the nearest start and end site of each type of regulatory region: transcription, 5' UTR, and 3' UTR; compared to the distribution expected by randomization (black line) at different classification confidence thresholds (columns).

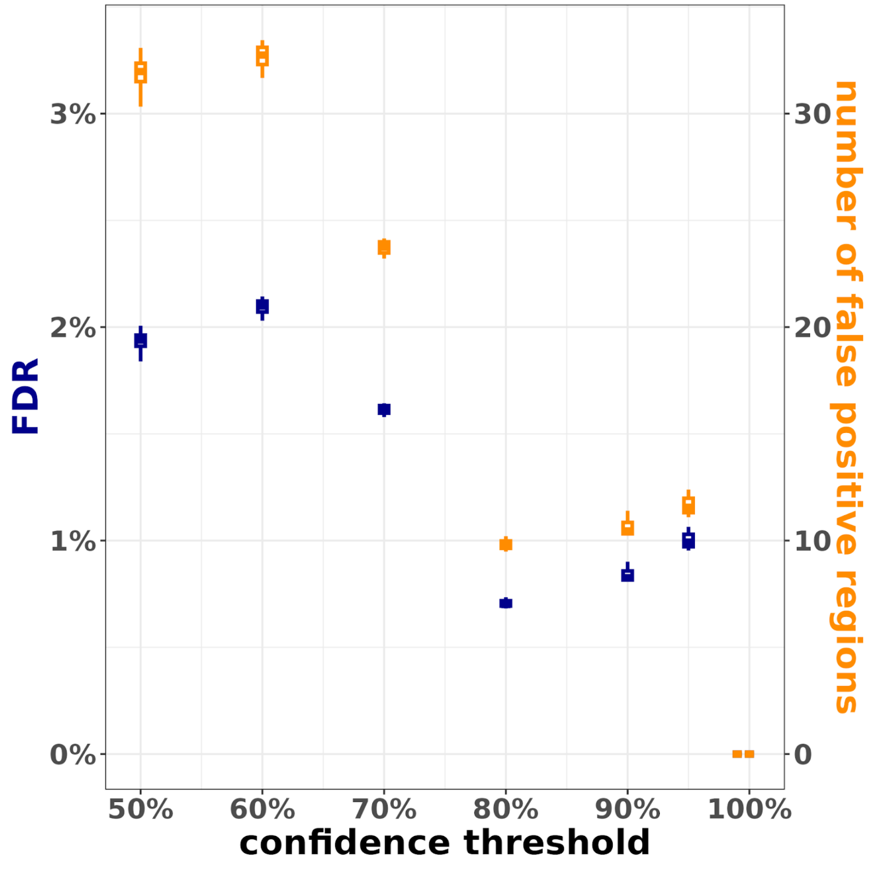


**Supplemental Figure 19** The false discovery rate can be tuned to different tolerances by increasing the confidence threshold at which a window is considered to be containing a sweep. This shows the FDR (blue) and number of false positive regions (orange) at different confidence thresholds when calculated for 100 different sets of independent windows each 1Mb apart.

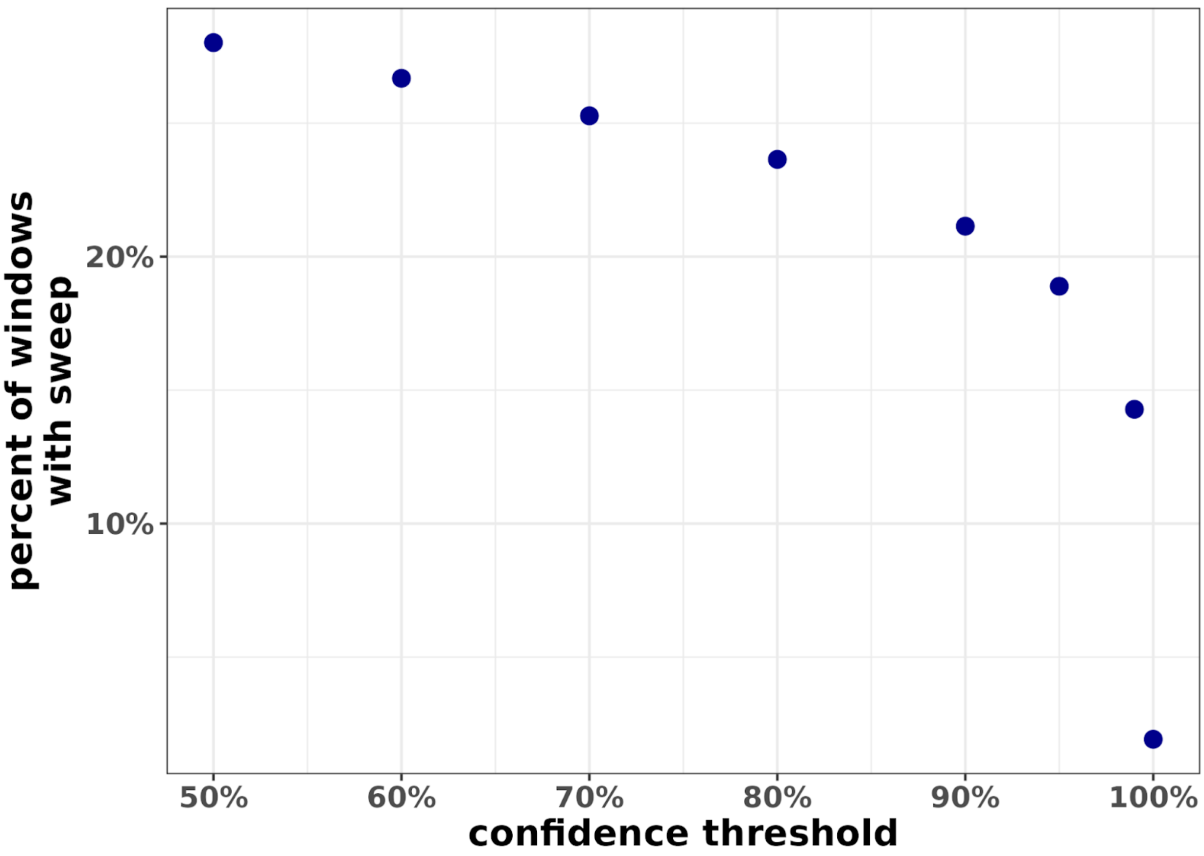


**Supplemental Figure 20** The percent of genomic windows classified as a sweep at different confidence thresholds.

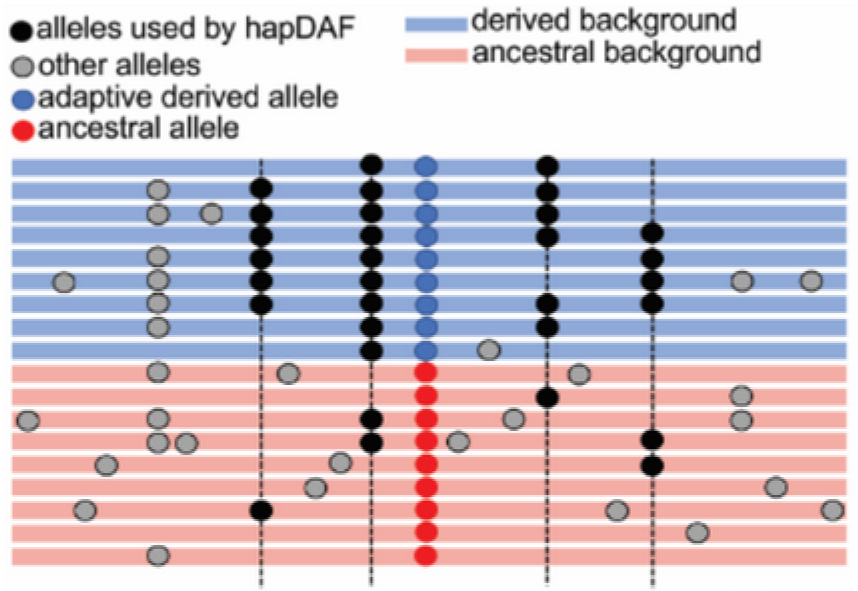


$f_{d1} = 6/9$ $f_{d2} = 9/9$ $f_{d3} = 6/9$ $f_{d4} = 4/9$
$f_{a1} = 1/9$ $f_{a2} = 2/9$ $f_{a3} = 1/9$ $f_{a4} = 2/9$

**Supplementary Figure 21**. *hapDAF* schematic. The focal variant (center, blue and red dots) determines the background of the sample (derived, blue, or ancestral, red). In a window around the focal variant, variable sites with alleles shared between derived and ancestral backgrounds are identified as shared flanking variants (black dots). Variants present in only one background (grey dots) are ignored. In this window there are four shared flanking sites across nine derived and nine ancestral backgrounds.

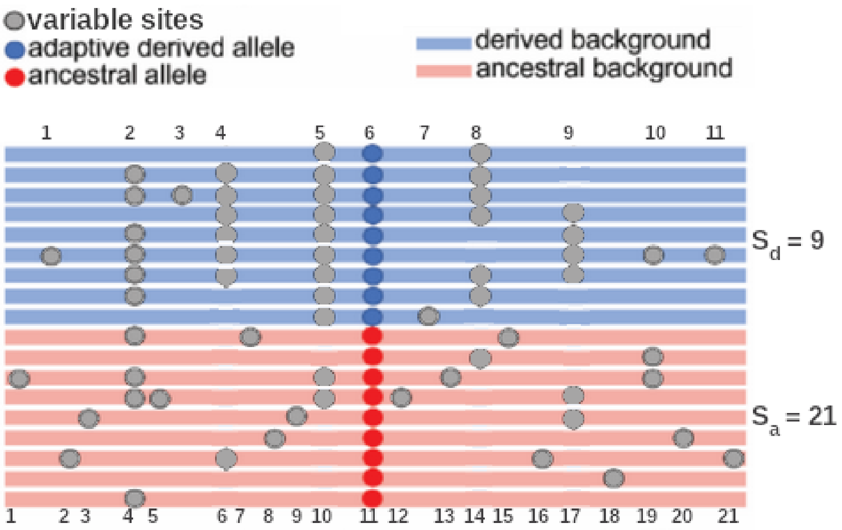


$S_d = \#variable\ sites\ on\ derived\ background$

$S_a = \#variable\ sites\ on\ ancestral\ background$

**Supplementary Figure 22**. *Sratio* schematic. The focal variant (center, blue and red dots) determines the background of the sample (derived, blue, or ancestral, red), and variable sites (grey dots) are identified in a window around the focal variant. Variable sites are identified separately on derived backgrounds and ancestral backgrounds. The number of variable sites is calculated for each background (derived, $S_d$ and ancestral, $S_a$), inclusive of the focal allele. In this window, there are nine variable sites on the derived background, and 21 variable sites on the ancestral background.

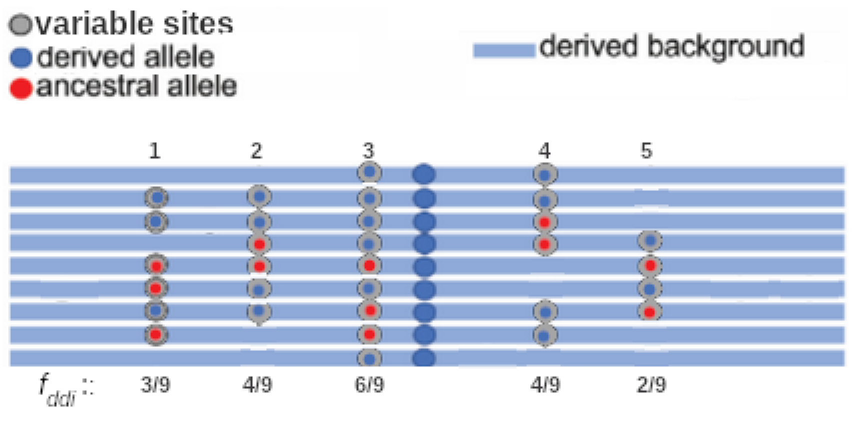


**Supplemental Figure 23**. *freq* schematic. The focal variant (center, blue dots) determines the background of the sample (derived, blue, or ancestral, red), and variable sites (grey dots with blue or red centers) are identified in a window around the focal variant. The frequency of the derived allele at each variable site ($f_{ddi}$, grey dots with blue centers) is calculated. In this window, there are five variable sites on only the derived background.

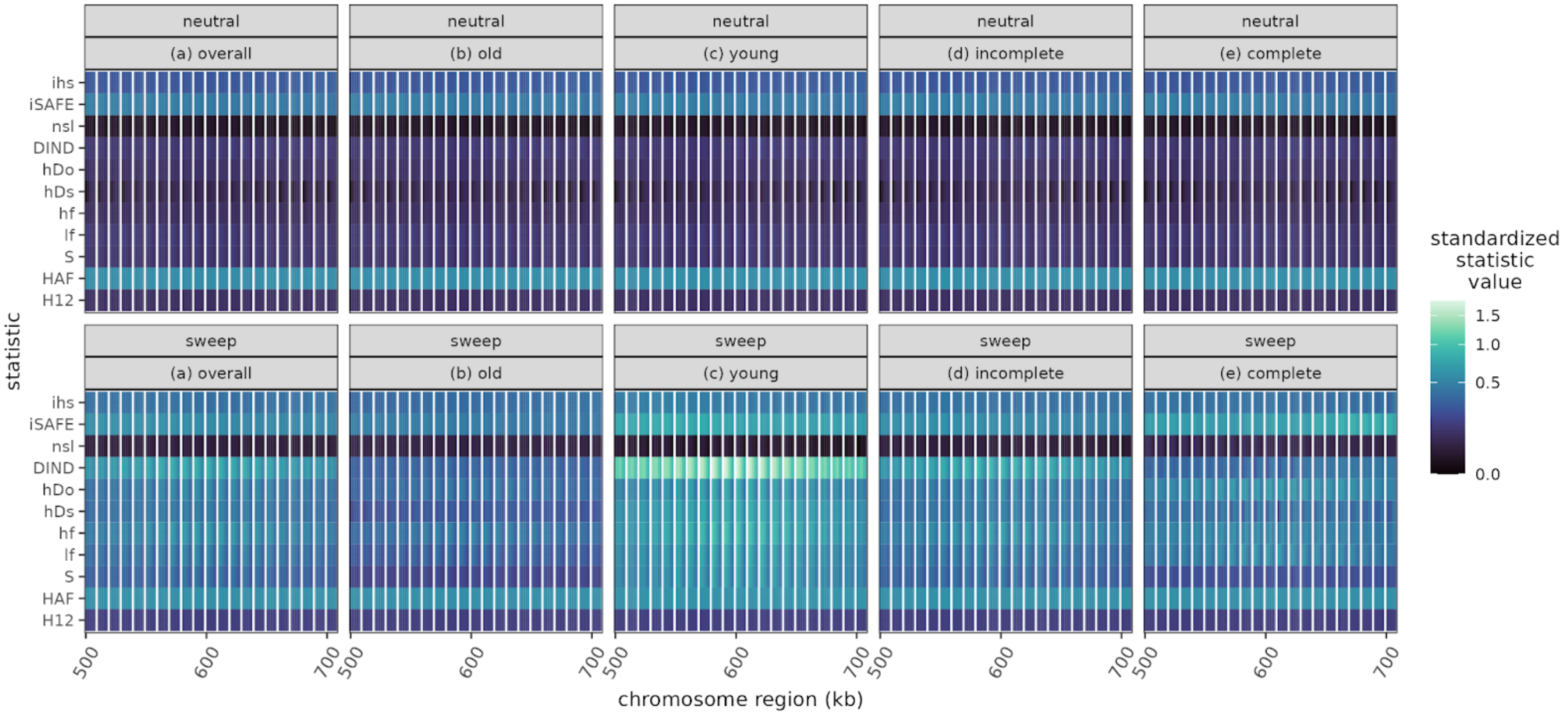


**Supplementary figure 24 Visualization of average feature vectors** Neutral (top) and sweep simulations (bottom), across (a) all combinations of parameter values, (b) sweeps 0.05 – 0.125 * $4N_e$ generations old, (c) sweeps up to 0.05 * $4N_e$ generations old, (d) incomplete sweeps, and (e) complete sweeps. Each feature vector is represented as an image, with statistics as rows and center point/window size combinations as columns.

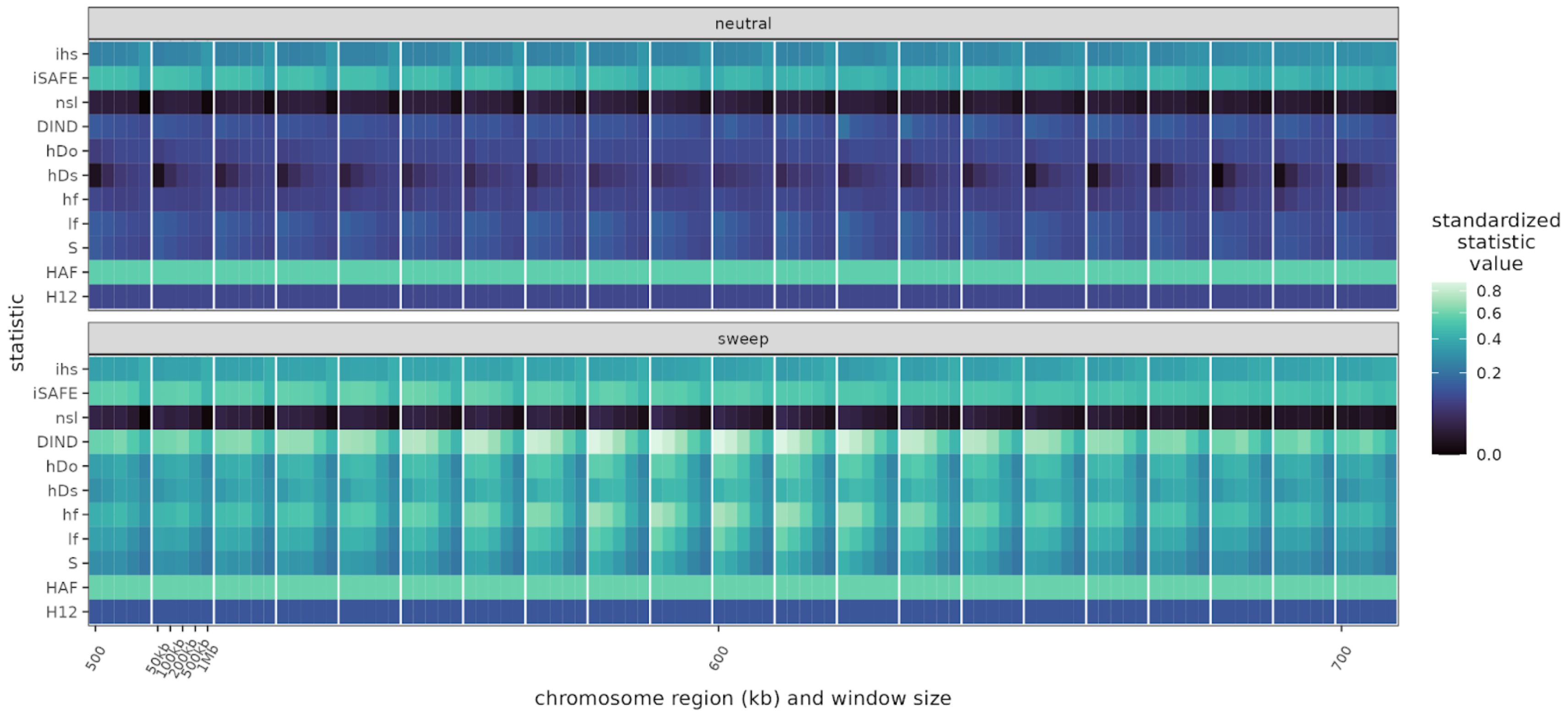


**Supplementary figure 25** Visualization of average feature vectors for neutral (top) and sweep simulations (bottom), across all combinations of parameter values. Each feature vector is represented as an image, with statistics as rows and center point/window size combinations as columns.

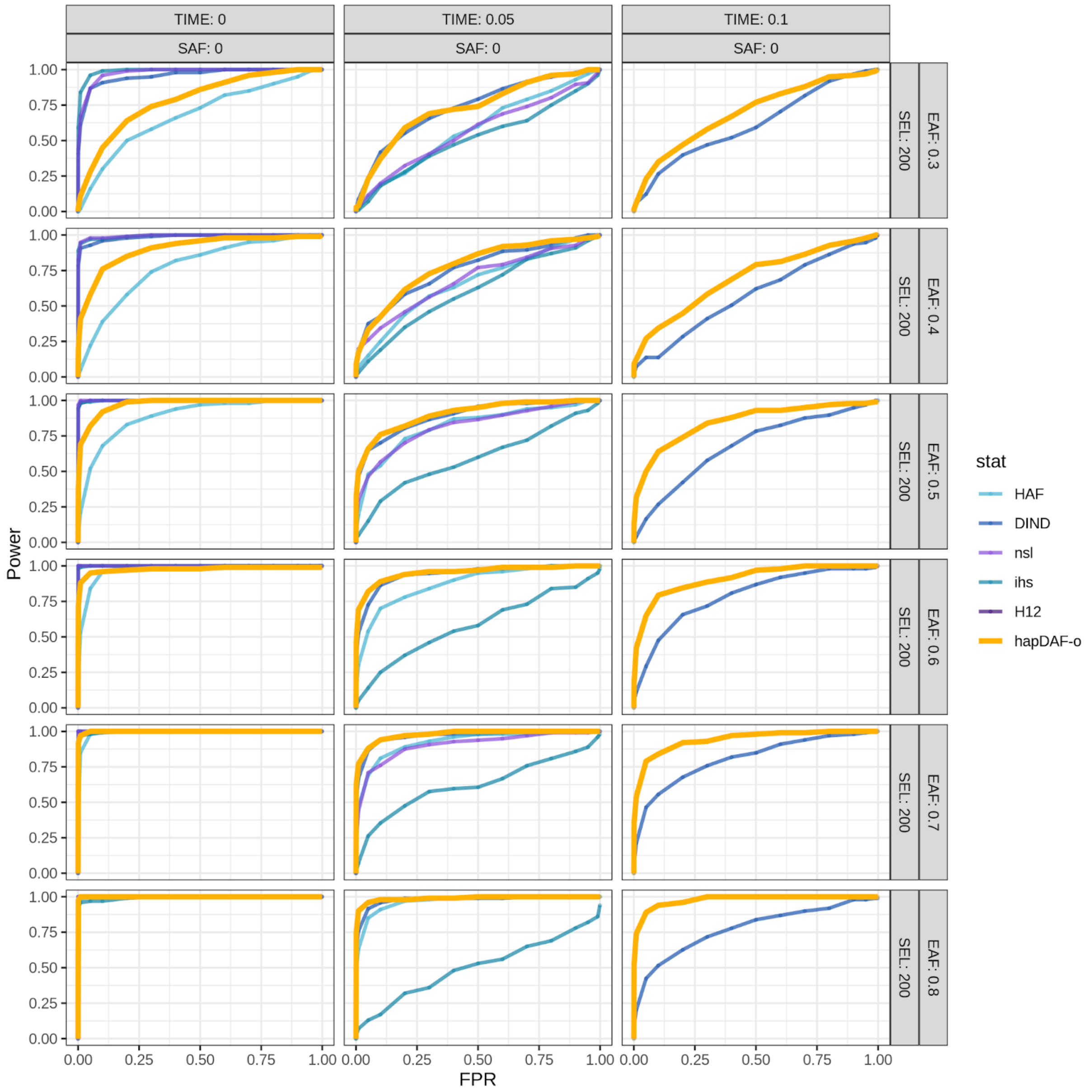


**Supplementary Figure 26** Power of *hapDAF-o* and existing statistics for incomplete sweeps from *de novo* mutation (starting allele frequency (SAF) = 0) ranging from recent (0 generations ago) to old (0.1 * $4N_e$ generations ago), at ending allele frequency (EAF) from 0.3 - 0.8 and selection strength (SEL) = 0.01 * $2N_e$.

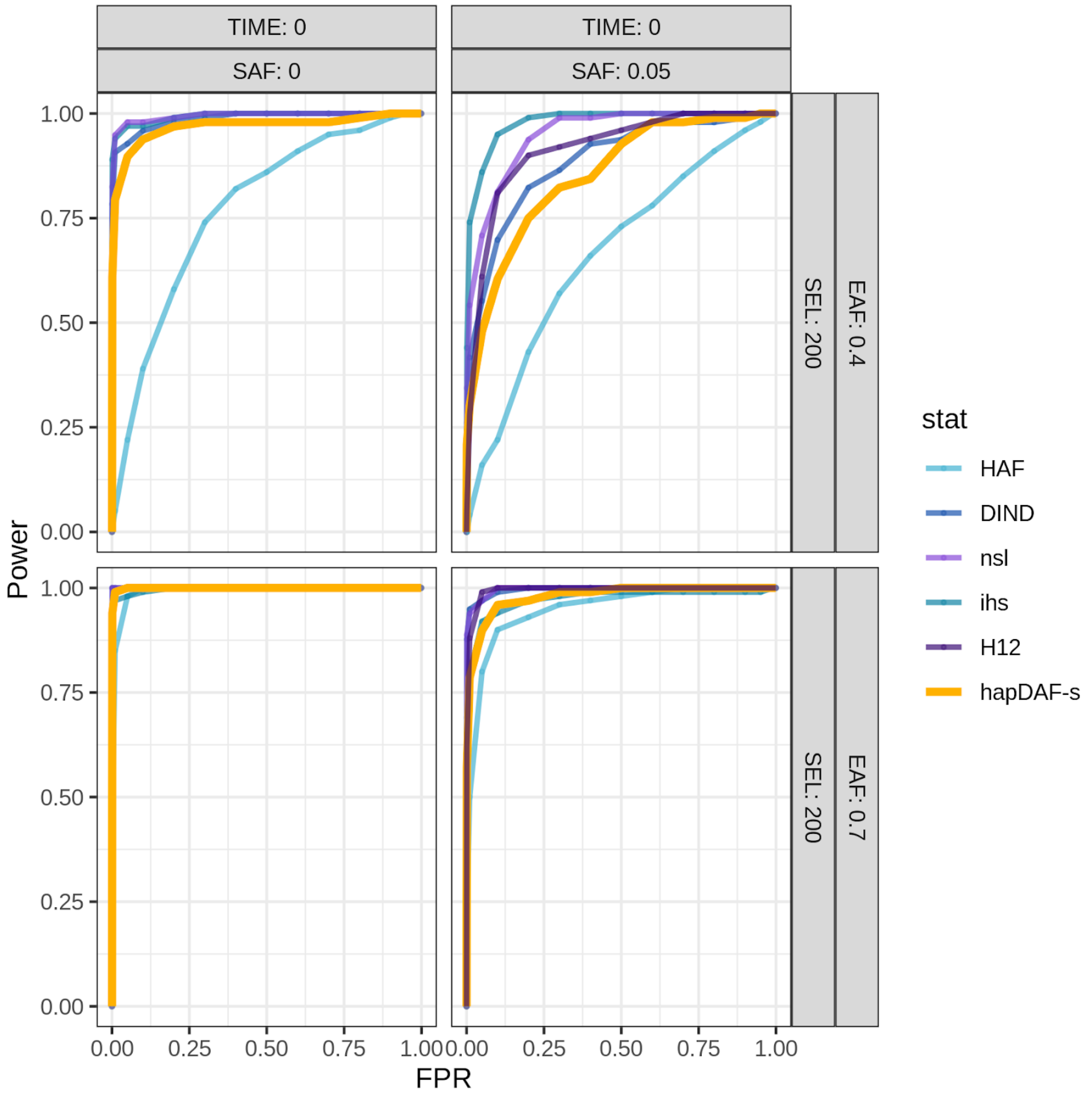


**Supplementary Figure 27** Power of *hapDAF-s* and existing statistics for incomplete (ending allele frequency (EAF) < 1), recent sweeps from *de novo* mutation (starting allele frequency SAF) = 0) and standing variation (SAF = 0.05) at ending allele frequency (EAF) of 0.4 and 0.7, at selection strength = 0.01 * $2N_e$.

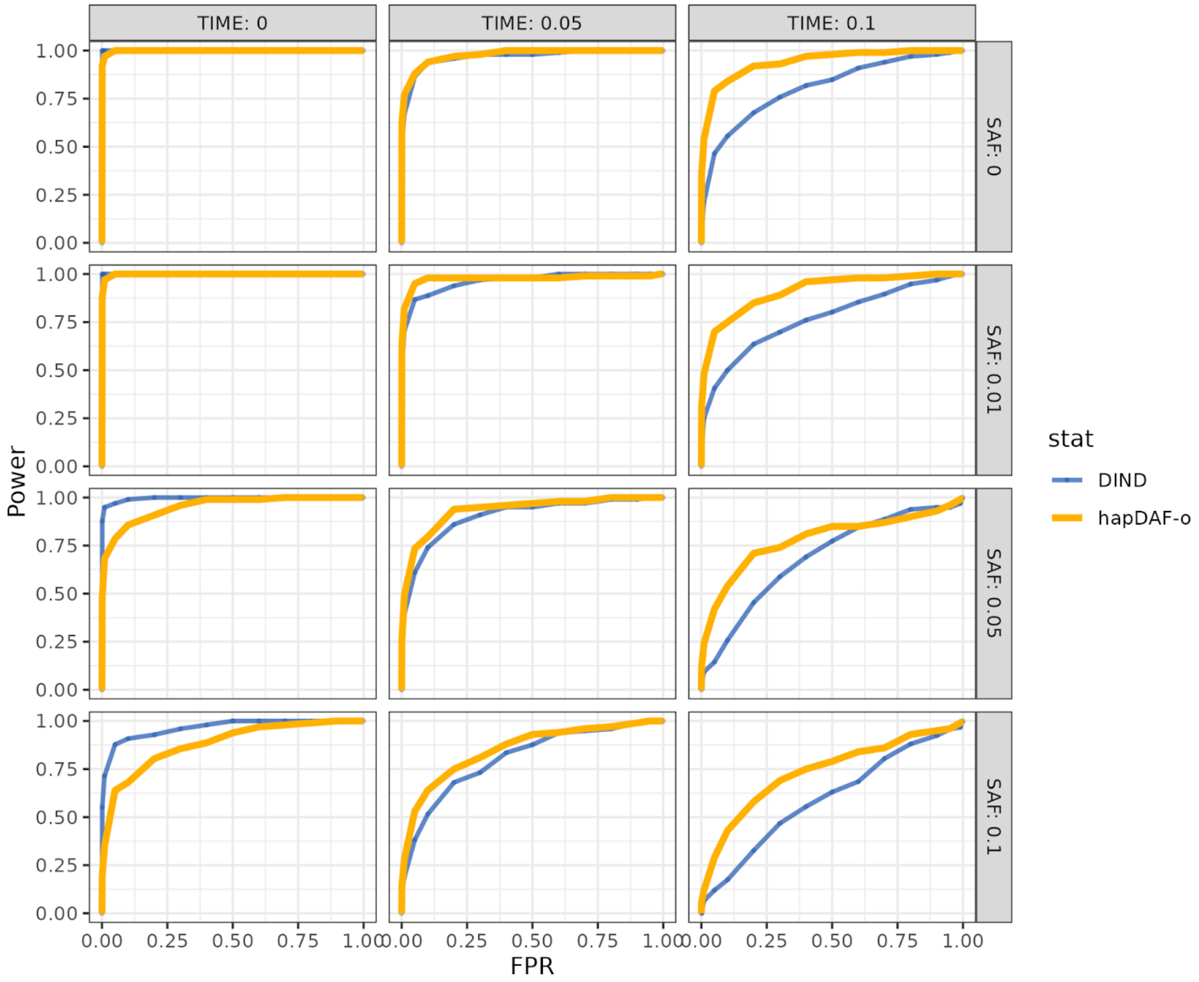


**Supplemental Figure 28** Power of *hapDAF-o* and DIND for complete sweeps (EAF = 1) of moderate selection strength = 0.01 * $2N_e$, comparing sweep time and starting allele frequency.

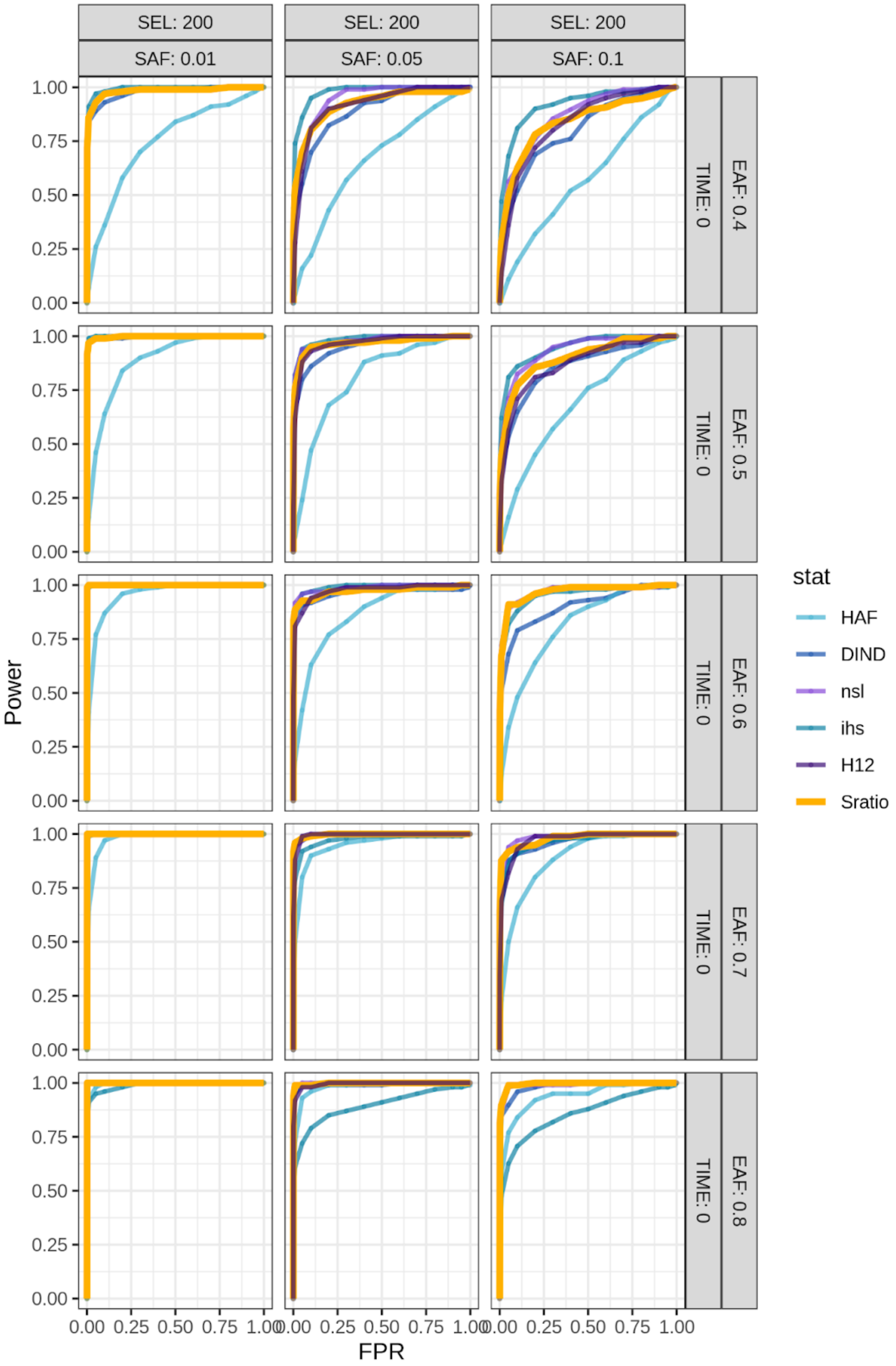


**Supplementary Figure 28** Power of *Sratio* and existing statistics for recent, incomplete (ending allele frequency (EAF) < 1), soft sweeps at starting allele frequencies (SAF) from 0 to 0.1 and final allele frequencies from 0.4 to 0.8, at selection strength (SEL) = 0.01 * $2N_e$.

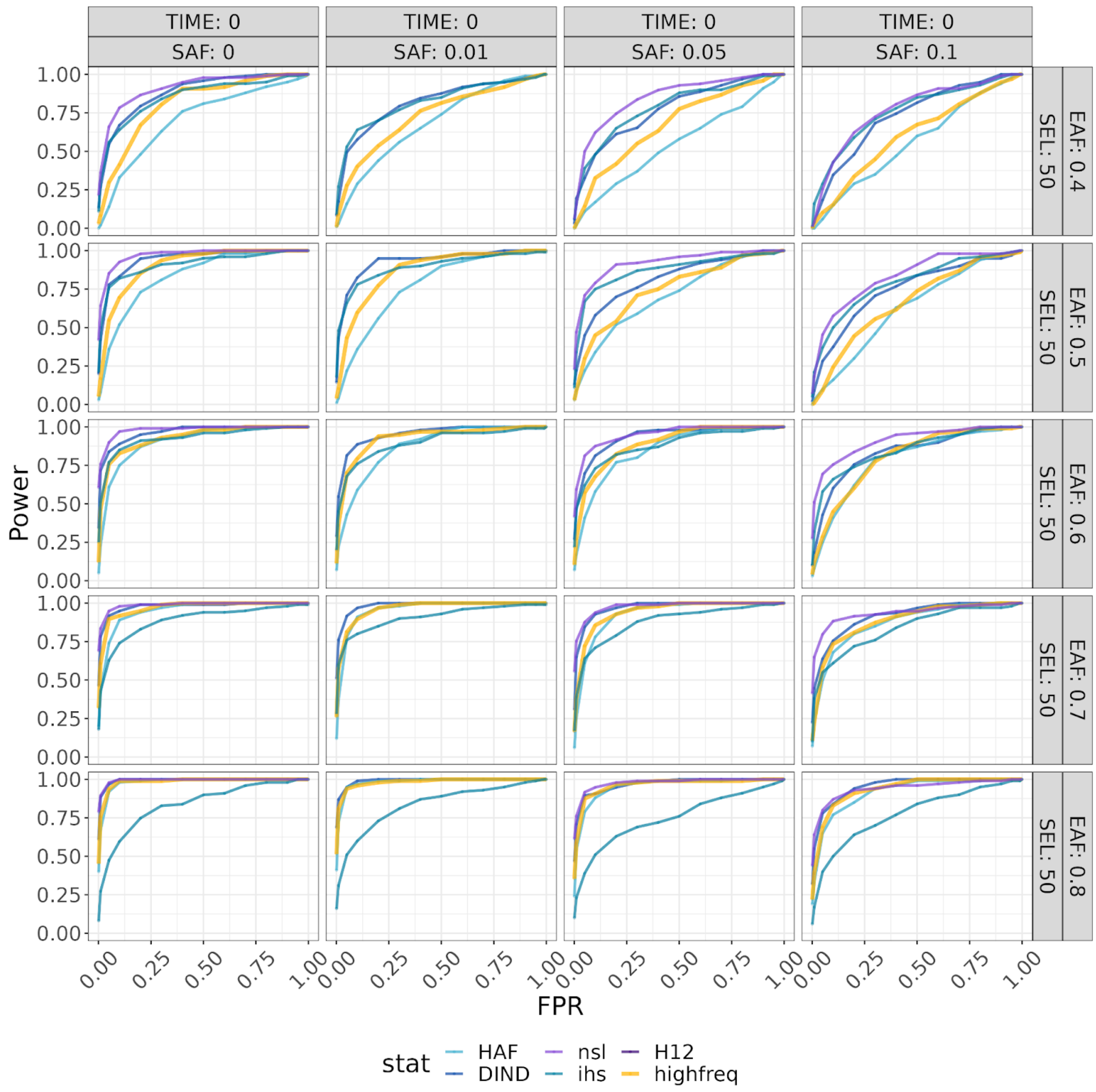


**Supplementary Figure 30** Power of existing statistics and *highfreq* (orange) for weak recent, incomplete (ending allele frequency (EAF) < 1), soft sweeps at starting allele frequencies (SAF) from 0 to 0.1, ending allele frequencies (EAF) from 0.4 to 0.8, at selection strength (SEL) = 0.005 * $2N_e$.

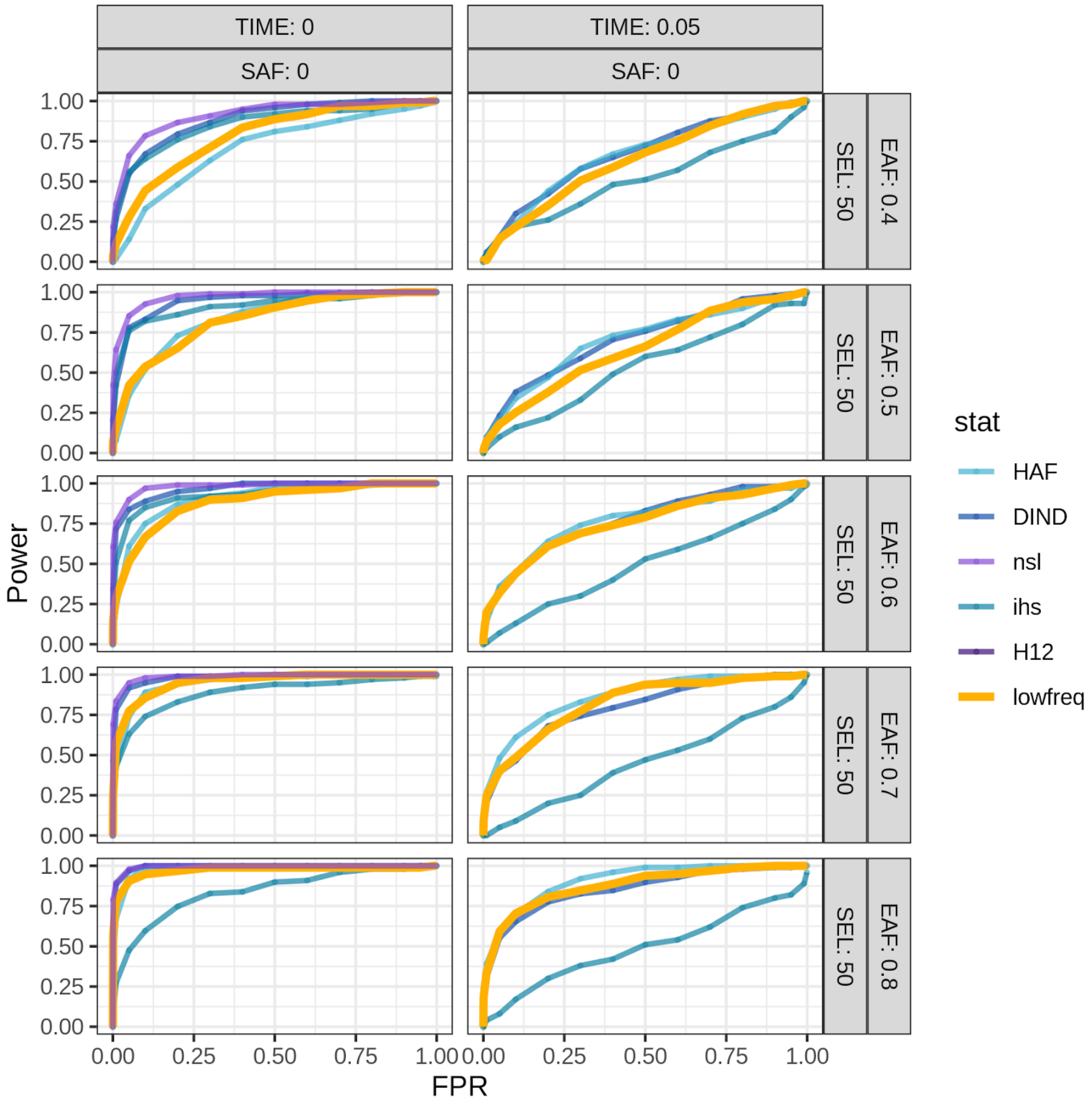


**Supplementary Figure 31** Power of existing statistics and *lowfreq* (orange) for weak, incomplete (ending allele frequency < 1) sweeps from *de novo* mutation (starting allele frequency (SAF) = 0) at ending allele frequencies (EAF) from 0.4 to 0.8 and selection strength = 0.005 * $2N_e$.

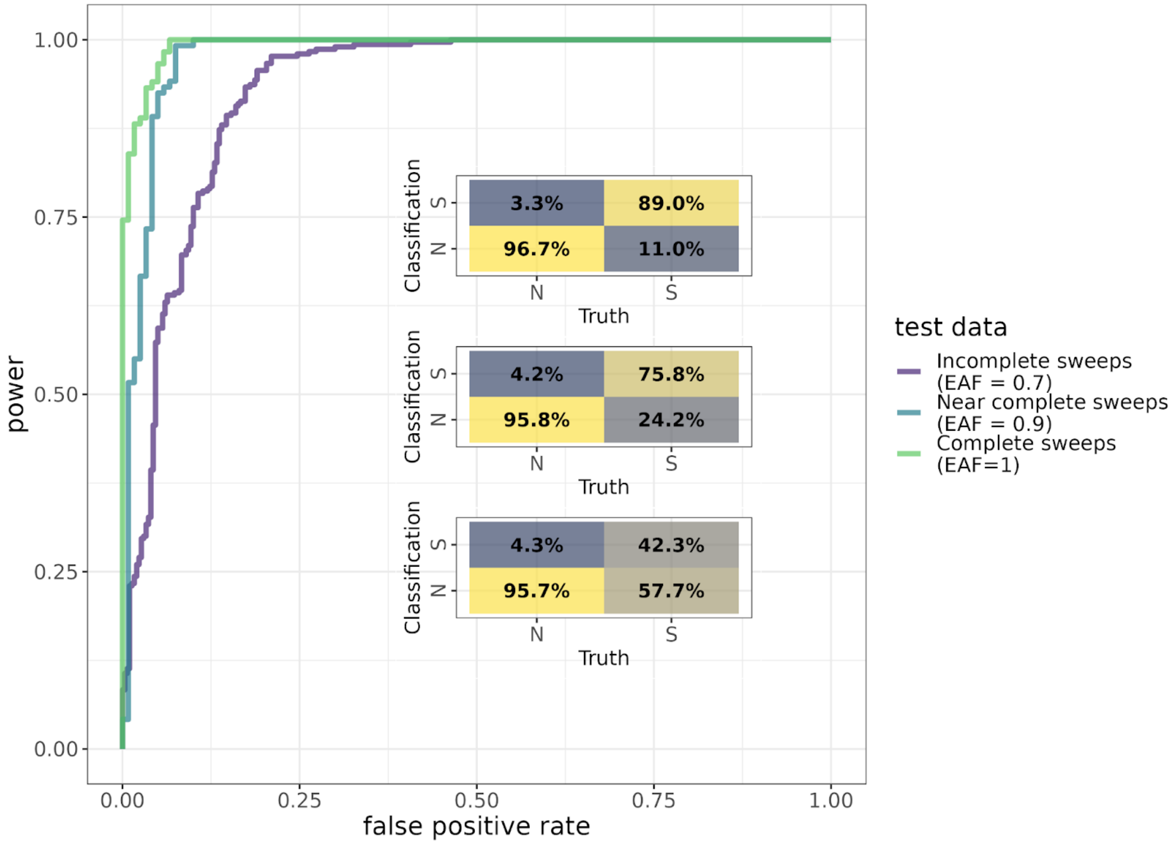


**Supplementary Figure 32** Classification of hard (SAF = 0), strong (s = 0.01) sweeps 10,000 generations old at three different levels of completeness (EAF = 0.7, purple, bottom matrix; EAF = 0.9, blue, middle matrix; EAF = 1, green, top matrix) generated with the Yoruba demographic model.

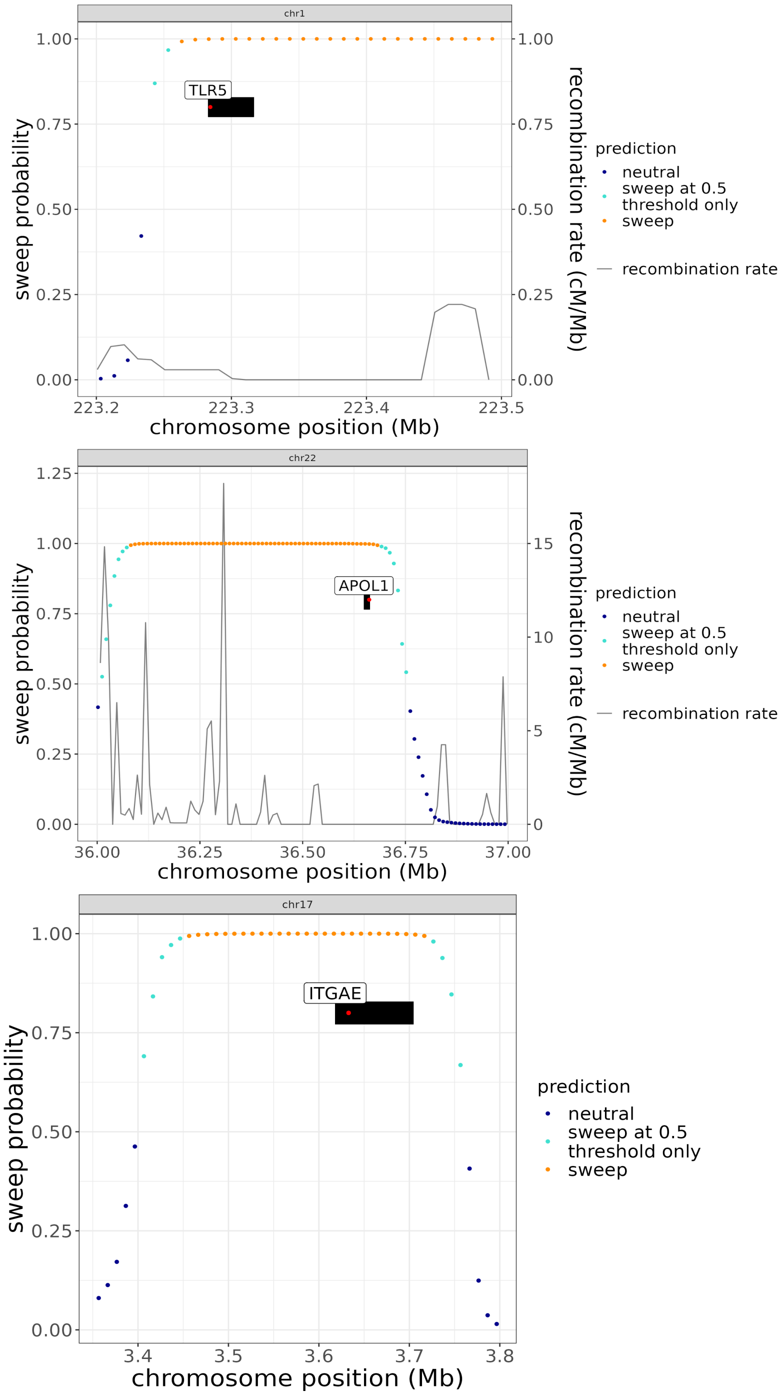


**Supplementary Figure 33** TLR5 (top), APOL1 (middle), and ITGAE (bottom) genes with the regional sweep probabilities and recombination rates. Each gene span is shown in black, with the causative polymorphism site(s) shown as a red point. The entirety of each of these genes lie within swept regions classified with high confidence by Flex-sweep. The grey lines show recombination rates in cM/Mb. No recombination rate is shown for the region including ITGAE on chromosome 17 because estimates for this region are not available in the deCODE map.

**Supplementary Table 1: csv file**

Supplementary Table 2**.** Distributions for parameters of equilibrium and human demography simulations

<table>
<tr><th>Simulation</th><th>$\mu$ (per site)</th><th>$\varrho$ (per site)</th><th>$s$</th><th>$f_{start}$ (start allele frequency)</th><th>$f_{end}$ (end allele frequency)</th><th colspan="2">$\tau$ (in generations)</th></tr>
<tr><td rowspan="3">equilibrium and simple size change</td><td rowspan="3">$\mu \sim U(2*10^{-9}, 5.2*10^{-8})$</td><td rowspan="3">$\varrho \sim exp(\beta=1*10^{-8})$</td><td rowspan="3">$s \sim U(0.001, 0.01)$</td><td rowspan="3">$f_{start}$ = {50% ~ $U(0, 0.1)$, 50% = 0}</td><td rowspan="3">$f_{end}$ = {90% ~ $U(0.2, 1)$, 10% = 1}</td><td colspan="2">$\tau \sim U(0, 5000)$</td></tr>
<tr><td>“recent”</td><td>“old”</td></tr>
<tr><td>$\tau \sim U(0, 2000)$</td><td>$\tau \sim U(2000, 5000)$</td></tr>
<tr><td>human demographic model</td><td>$\mu \sim$ truncated $N(1.25*10^{-8}, 1*10^{-8})$, minimum = $2*10^{-9}$, maximum = $5.2*10^{-8}$</td><td>$\varrho \sim$ truncated $N(5*10^{-9}, 1*10^{-8})$, minimum = $1*10^{-10}$, maximum = $1*10^{-8}$</td><td>$s \sim U(0.001, 0.01)$</td><td>$f_{start} \sim$ 50% U(0, 0.1), 50% 0</td><td>$f_{end} \sim$ 90% $U(0.2, 1)$, 10% 1</td><td colspan="2">$\tau \sim U(0, 5000)$</td></tr>
</table>

Supplementary Table 3. Demographic parameters and timings for non-equilibrium demography simulations

| Demographic event | magnitude | sampling time after last event, in generations | selection time |
|---|---|---|---|
| decline | 1x, 5x, 10x | $0.05N_e$, $0.1N_e$, $0.5\ N_e$ | $\tau$ ~ *U*(before decline, 5000) |
| expansion | 5x, 10x, 50x | $0.1N_e$ | $\tau$ ~ *U*(before expansion, 5000) |
| full recovery post bottleneck | 5x | 1, $0.05N_e$, $0.1N_e$ | $\tau$ ~ *U*(before decline, 5000) |
| human* | NA | NA | $\tau$ ~ *U*(0, 5000) |

* Human demography refers to the Yoruba population history inferred by (Speidel et al. 2019)

Appendix

Architecture of Convolutional Neural Network

- Number of samples processed simultaneously (batch size): 32
- 3 convolutional layers each with 3 convolution steps
    - Filters per layer:
        - Step 1: 64
        - Step 2: 128
        - Step 3: 256
    - Filter sizes per layer per step:
        - Layer 1: 3
        - Layer 2: 2
        - Layer 3: 2
    - Dilation:
        - Layer 1: none
        - Layer 2: 1x3
        - Layer 3: 1x5
    - 1 pooling step per layer
    - Dropout rate per layer: 0.15
- 2 fully connected layers
    - Dropout rate per layer: 0.2
    - Neurons per layer:
        - Layer 1: 512
        - Layer 2: 128
    - Activation: Rectified Linear Unit (ReLU)
- Output layer:
    - Activation: Sigmoid
    - Kernal Initialization: Glorot (Xavier) Uniform
- Optimization: AMSGrad, epsilon = $10^{-7}$
    - Learning function: Cosine Decay with Restarts, initial learning rate = 0.001, decay over 300 steps
- Loss function: binary crossentropy
- Maximum epochs: 100
    - early stopping with patience 5
