## Supplementary methods and results for "Versatile detection of diverse selective sweeps with Flex-sweep"

**TABLE OF CONTENTS**

1. **Supplementary Methods**
   1.1 New statistics
       1.1.1 *hapDAF*
       1.1.2 *Sratio*
       1.1.3 *freq*
   1.2 Convolutional neural networks for sweep detection
   1.3 Feature vectors
   1.4 Feature vector preprocessing
   1.5 Network architecture
   1.6 Architecture evaluation
   1.7 Impact of new statistics
   1.8 Statistic importance
   1.9 Impact of background selection and recombination rate heterogeneity
   1.10 Mispolarization
   1.11 Sweep parameters
   1.12 Sweep type classification power
   1.13 False discovery rate of sweep regions
   1.14 Genome-wide patterns of selective sweeps
2. **Supplementary Results**
   2.1 Power of new statistics
       2.1.1 *hapDAF*
       2.1.2 *Sratio*
       2.1.3 *freq*

###### 2.2 Model Calibration

###### 2.3 Detecting older sweeps

2.4 Comparison to state-of-the-artty
2.5 Statistic importance
2.6 Sweep type classification power
2.7 Low recombination rate
2.8 False discovery rate of sweep regions
2.9 Genome-wide patterns of selective sweeps
2.10 Previously identified genes

### Supplementary Methods

###### New statistics

These five new statistics (hapDAF-o, hapDAF-s, Sratio, lowfreq, and highfreq) take advantage of the hitchhiking to higher frequency of alleles in regions linked to an adaptive allele by comparing diversity between ancestral and derived haplotypes. This is the structure that was first used by the DIND statistic (Derived Intra-allelic Nucleotide Diversity, Barreiro et al. 2009) using π (pairwise genetic diversity) within a fixed window size around a focal variant. These new statistics instead calculate the number of shared variants between ancestral and derived backgrounds (*hapDAF-o* and *hapDAF-s*), the number of variants in a haplotype (*Sratio*), and the frequencies of derived alleles (*highfreq* and *lowfreq*). The pair of *hapDAF* statistics are similar to the Difference in Allele Frequency (DAF) statistic (Carneiro et al. 2014) in that they compare allele frequencies across two haplotypes to detect regions that are very different. While the original DAF statistic compares haplotypes between populations, *hapDAF* compares the ancestral and derived haplotypes. *hapDAF-o* is particularly important because it has power to detect old sweeps, including old, incomplete sweeps.

*hapDaf* (Supplementary figure 21)

*hapDAF-o*: This version of the *hapDAF* statistic is optimized to identify older, incomplete sweeps, by tuning the frequency thresholds of the shared flanking variants such that $f_{di}+f_{ai}>0.25$ and $f_{ai}<0.25$. Only variants that have hitchhiked to high frequency will stay informative as recombination and drift break up haplotypes and eliminate rare variants. By restricting the calculation to shared flanking variants with high combined frequency across both backgrounds, and allowing moderate frequency of shared flanking variants on the ancestral background, we increase the chance of finding focal alleles on backgrounds that have undergone considerable recombination and drift through time. While this increases the power to find old sweeps, it may reduce the power to find sweeps from standing variation that did not consistently increase the frequency of shared variants on a single derived background (as seen in comparison to eg. DIND, see below in **supplementary results**).

*hapDAF-s:* This version of the hapDAF statistic is optimized to identify recent, incomplete sweeps from standing variation, by tuning the frequency thresholds such that $f_{di}+f_{ai}>0.1$ and $f_{ai}<0.1$. An excess of rare variants provides a good signal of a selective sweep in the recent past, i.e. before they have been eliminated by drift. By allowing the calculation to include shared flanking variants with low combined frequency across both backgrounds and restricting it to alleles that are rare on the ancestral background, we increase the number of informative variants for recent sweeps, thus increasing the power of the statistic. Increasing the number of informative variants increases the chance of finding focal alleles on backgrounds that have not consistently increased the frequency of shared variants on derived backgrounds (due to incomplete sweeps that do not strongly decrease variation on the derived background, or soft sweeps that have many derived backgrounds), but have increased the frequency of shared variants that are rare on ancestral backgrounds. These thresholds were chosen based on simulation-based power analyses.

Sratio (Supplementary figure 22)

The segregating sites ratio (*Sratio*) compares the number of variable sites in a window of fixed size around a focal derived allele with the number of variable sites around the corresponding ancestral allele. Both incomplete and soft sweeps are not expected to allow a single haplotype to hitchhike to high frequency but do reduce variation on the derived background relative to the ancestral background. Therefore *Sratio* does not look for similarities in haplotypes around a derived allele but instead for small differences in the amount of variation around the focal variant, as measured by the number of variable sites. Regions around selected alleles are expected to have lower variation than those around the ancestral allele, thus a low *Sratio*.

freq (Supplementary figure 23)

The two *freq* statistics look for an excess of low or high-frequency alleles on the derived background by calculating the frequencies of derived alleles in a window of fixed size around a focal variant. These two statistics complement each other by focusing on different ends of the site frequency spectrum (SFS). That is, these statistics take advantage of the same expected excess of low and high-frequency alleles as do Tajima’s D (Tajima 1989) and Fay and Wu’s H (Fay and Wu 2000), respectively. By taking the standardized difference between two SFS estimators, these classic statistics risk losing power when the variance is high, for example due to recombination (Zeng et al. 2007) or sequencing error (Korneliussen et al. 2013). Instead, the *freq* statistics take advantage of genome-wide data to standardize the frequency of the derived allele on the derived background against the genome-wide distribution of focal variants with similar derived allele frequencies, avoiding the loss of power these classic statistics can experience under conditions with high variance

###### Convolutional neural networks for sweep detection

The use of Convolutional Neural Networks (CNNs) for sweep detection and other population genetic inference has been comprehensively described in (Kern and Schrider 2018) and reviewed in (Schrider and Kern 2018) and (Flagel et al. 2019). Briefly, CNNs are a form of supervised deep learning that take input images labeled by class, process these images to “learn” common distinguishing traits of images in each, and use these learned traits to classify novel images. At their heart is the concept that only certain traits of an image are useful to distinguish its class from another class. While they were developed to classify images, under the hood they are processing numeric pixel values. The key innovation in the application of CNNs to population genomic data was that the numeric values processed by the CNN need not have their origin in an image but could be the values of various summary statistics calculated across a sample of loci (Kern and Schrider 2018) or genome alignment (Flagel et al. 2019). These summary statistics can be ordered in a feature vector (or alignment), which is then transformed into a two (or more) dimensional array before being passed to the CNN.

###### Feature vectors

Feature vectors represent the summarized variation at each simulated locus, with one vector per simulation. We divide each simulated 1.2 Mb locus into a series of 21 center points spaced at 10kb intervals, each with nested windows of five different sizes ({50kb, 100kb, 200kb, 500kb, 1000kb}) around the center point. We then calculate 11 summary statistics to summarize the characteristics of the sample with respect to site frequency spectrum (SFS), haplotype structure, and diversity on the derived background (Table 1) in each of the five nested windows Nested windows are used because sweep signals vary in strength at different distances from the focal variant, as also described in (Caldas et al. 2022). For example, recombination reduces signal more as the distance from the selected variant increases, so large windows are expected to have more noise and lower signal for older sweeps. However, for recent and strong sweeps, large windows include useful information and increase the power to identify these sweeps. By including windows of multiple sizes, the CNN is able to explore these aspects of sweep signals, retaining information that helps it to classify sweeps and discarding information that doesn’t.

These statistics are normalized against the genome-wide distribution of focal variants with similar derived allele frequencies (as described in Voight et al. 2006) using the neutral simulations, with the exception of *HAF* and *H12*. These two statistics take a single value across the entire locus because they require a larger window to define haplotypes, so this value is repeated for each center point and window combination to maintain appropriate dimensions of the feature vector.

These normalized summary statistics are combined in a feature vector 1155 elements long, by statistic, center point (i), and window size (j), with each statistic, center point, and window represented by $stat_{i,j}$:

$\left[ iHS_{1,1},iHS_{1,2}...iHS_{2,1},iHS_{2,2}...iSAFE_{1,1},iSAFE_{1,2}...H12_{21,3},H12_{21,4},H12_{21,5} \right]$

Each feature vector is then translated into a two-dimensional image, with values of each statistic as a row, and each center point/window size combination as a column (Supplementary figures 24, 25). Feature vectors can be generated with or without a recombination map.

###### Feature vector preprocessing

Preprocessing includes strategies such as normalization and standardization (Bishop 1998). Multiple preprocessing strategies were tested to explore the impact of different types of feature vector rescaling and standardization on network success. All statistics except HAF and H12 were already of a similar order of magnitude because of the initial normalization against the genome-wide distribution of focal variants with similar derived allele frequencies. HAF could take on a much larger range of values, so feature vectors also were tested in three additional ways: with HAF/H12 normalization, with HAF scaled by a factor of 10 to bring its range of variation more in line with that of the other statistics, and without HAF and H12. This last is because HAF and H12 each take on a single value for the whole locus, repeated multiple times in the feature vector to maintain appropriate dimensions. Thus it is possible that these two statistics could overwhelm the signal from the other statistics.

The impact of rescaling feature vectors was tested in the following ways: Feature vectors without any rescaling, feature vectors rescaled proportionally such that each vector fell within the range 0-1 (both forcing all statistics to the same order of magnitude and to positive values which can be more compatible with some activation functions), and feature vectors with each statistic value rescaled by the mean and standard deviation of: 1. that statistic across all center points and window sizes; 2. that statistic across all center points, within window sizes; or 3. that statistic across all window sizes, within individual center points.

The most effective rescaling strategies across model architectures and data sets were: no rescaling, rescaling proportionately across the entire feature vector such that each vector fell within the range 0-1 (both forcing all statistics to the same order of magnitude and forcing them to positive values which can be more compatible with some activation functions), and rescaling subsetted by each statistic, proportionally within its center point, window size, or both. While the subsetted rescaling strategies gave the best performance under a few demographic scenarios, they were also more affected by demographic scenario and model misspecification (Supplementary figure 1). Not rescaling provided the most consistently good performance across all tested scenarios.

###### Network architecture

To develop the final model, we tested model architectures using the simulations described above with a variety of: number of layers (3-4), number of convolution steps per layer (2-3), number of filters per convolution step (32-256), filter sizes (2x2, 3x3, 5x5, or with dilation 1x3, 1x4, 1x5), number of pooling steps (1-2), number of fully connected layers (2-3), and number of neurons in each fully connected layer (32-512). In total, 96 different architectures were tested using these combinations. After limiting the possible architectures to the top 9 performing models on equilibrium, as well as correctly and mis-specified demographic scenarios, we tested different combinations of learning functions, activation functions, and regularization techniques. Learning optimization functions included Adam, Nadam, Cosine decay with restarts, AMSGrad, and Stochastic gradient descent (SGD) with and without Nesterov momentum. Activation functions included ReLU, Leaky ReLU, and PReLU, ELU. Addition of a squeeze and excitation layer was also tested, but this added no benefit. Regularization methods included early stopping, batch normalization, and L1 (lasso), L2 (ridge), and L1L2 regularization to improve generalization with multicollinearity.

These stratagies can interact in unpredictable ways (Liao et al. 2022), so combinations were tested comprehensivtly to find those that produced the best classification results under multiple scenarios. Strategies that were found to improve the network, as determined by comparisons of AUC, accuracy, and false positive rate, were then tested with adjustments to one or more tuning parameters, including learning rate. Tensorflow’s default learning rate of 0.01 provided inconsistent performance among runs, suggesting that it struggled to find optima. Learning rates below 0.001 performed less well, suggesting that they fell into local optima. A learning rate of 0.001 performed better than 0.01 or 0.0001, and all further tests using a constant learning rate (ie. not a learning rate scheduler) used this value. Models implementing early stopping with patience set to 5 avoided overfitting (Supplementary figure 2).

The best model architectures had a few things in common: three (rather than four) convolutional layers, an increasing number of filters per step, constant filter sizes for all steps, and one pooling step for each convolutional layer. Models with two or three fully connected layers both worked well.

###### Architecture evaluation

We calculated true and false positive rates, accuracy, and precision for each combination of model architectures and strategies and constructed receiver-operating characteristic (ROC) curves for visualization and area under the curve (AUC) calculations. Initial evaluation was done by comparing ROC curves, and strategy combinations that consistently performed worse were discarded.

Comparison to state-of-the-art methods: We also compared the success of Flex-sweep to diploS/HIC, one of the state-of-the-art machine learning methods designed to detect selective sweeps (Kern and Schrider 2018). diploS/HIC uses a similar CNN-based architecture to detect and distinguish among hard sweeps, soft sweeps, and regions linked to each. In order to maintain the most power to detect ancient sweeps, we do not attempt to distinguish between sweep types or linked regions and thus do not include separate simulations with regions linked to (but not including) sweeps, so for appropriate comparison to diploS/HIC we modified its original classification structure from five categories (neutral, hard sweep, soft sweep, linked hard, linked soft) to binary classification (neutral or sweep). For this comparison, we used the same simulations generated for the rest of the analyses, with an additional set of sweep simulations in the same parameter space except with only complete sweeps.

###### Impact of new statistics

To understand how much influence the new statistics (*highfreq, lowfreq, Ratio, hapDAF-o, hapDAF-s*) have on the quality of the predictions, we compared the classifications results of the network using only the new statistics vs. only the previously published statistics. In addition, we tested if the order of the statistics in the input tensor affects performance. The original order is shown in Supplementary figures 24 and 25. We tested two additional orders, one interspersing statistics by the type of data they summarize, and one grouping them by the type of data they summarize (Table 1, new order: DIND, HAF, hapDAF-o, iSAFE, highfreq, hapDAF-s, nSL, Sratio, lowfreq, iHS, H12).

###### Statistic importance

To explore how each statistic contributes to Flex-sweep classification, we used Deep SHAP (SHapley Additive exPlanations) to interpret the contribution of each feature (a statistic calculated at a region in the locus over a specific window size) (Lundberg and Lee 2017). The Shapley value of a feature in a feature vector is the average of its marginal contribution to the loss function over permutations of that feature, holding constant the other feature values. In other words, each time the feature values are permuted, the model re-classifies the feature vector and we measure the difference between the classification and the true state. This marginal contribution is then averaged over permutations and, in Deep SHAP, aggregated across feature vectors (Lundberg and Lee 2017; Molnar 2022). One advantage of SHAP is the game theoretic foundation of the underlying Shapely values, which ensures that importance values are sensitive to both the model and the data (that is, that the result is not dependent only on large differences in feature values regardless of their contribution to the model output). In addition, the contribution of each feature is additive and sums to the model loss (allowing averaging over feature contributions), and the contributions are distributed across features without bias.

However, as with most permutation-based interpretation methods, Deep SHAP is not able to incorporate information about feature correlations, which could lead to a single feature absorbing importance from other, correlated features. We calculated SHAP values (Lundberg and Lee 2017) for the equilibrium-trained, YRI demography-trained, and decline-trained models and interpreted the results in their corresponding testing data following two approaches: 1. clustering features by similarity, and 2. averaging the importance across each statistic at small (50kb-100kb), medium (200kb-500kb) and large (1Mb) window sizes in the middle (the 7 windows in the center of the locus) and shoulders (the leftmost and rightmost 7 windows of the locus) of each region.

###### Impact of background selection and recombination rate heterogeneity

Background selection simulations were performed with three different structures of “coding” sites: a generic structure in which randomly selected nucleotides making up 5% of the central 100kb of the locus were considered coding; a single gene structure in which the central 50kb of the locus was divided into 10 evenly-spaced, 100 bp long “exons” making up 5% of the region; a multi-gene structure in which the central 500kb of the locus was divided into 5 evenly-spaced “genes,” each made up of 10 evenly-spaced, 100 bp long “exons.” In each case, 75% of mutations in coding sites had the deleterious DFE and 25% were neutral, following Schrider 2020. We found the distribution of coding sites to have little impact on predictive capabilities, so the “generic” structure was used for analyses unless otherwise specified. To generate simulations with recombination rate heterogeneity similar to that observed in humans, we subsetted 100 random 1.2 Mb regions from chromosome 1 of the sex-averaged deCODE map (Halldorsson et al. 2019)) and provided these to SLiM 3.

###### Mispolarization

To understand the effect of mispolarization of alleles, we classified the equilibrium demographic simulation data after switching the ancestral and derived states in 0.1%, 1%, 5%, and 10% of SNPs, using the estimated mispolarization rate in human data of 1-4% (Glémin et al. 2015) as a guide and exceeding those estimates to account for the possibility of more uncertainty in non-model species data.

###### Sweep parameters

We explored the what simulation parameter values are associated with false positives and false negatives. We classified each neutral simulation as a false positive or true negative, and each sweep simulation as a false negative or true positive, and used t-tests to compare the parameter values in each case.

###### Sweep type classification power

To test whether Flex-sweep could classify the types of sweeps without losing power, we generated additional simulations in the same parameter space described previously, except fixing the starting allele frequency and ending allele frequency to generate sets of hard complete, hard incomplete, soft complete, and soft incomplete sweeps. The resulting data set included 50,000 simulated regions, 10,000 each of neutral, hard complete, hard incomplete, soft complete, and soft incomplete. We modified the CNN architecture to use softmax activation rather than sigmoid in the output layer to classify these five categories: the four types of sweeps, and neutral. Training and classification were otherwise run as previously described.

###### False discovery rate of sweep regions

We developed a false discovery rate (FDR) metric to demonstrate its actual performance in this real-world data set by comparing the proportion of windows classified as sweeps in the Yoruba data to the false positive rate from the validation tests using the Yoruba demographic model. We take a set of non-overlapping windows each separated by 1Mb and use the false positive rate (FPR = # false positives / (# false positives + # true negatives) ) and false negative rate (FNR = # false negatives / (# false negatives + # true positives)) from training to calculate the number of these independent windows falsely classified as swept. Since these windows are 1Mb apart, they more closely approximate the structure of the training and testing data. The FDR is then calculated from the number of these false positive windows and the total number of windows classified as a sweep in this set of independent windows as: FDR = # false positive windows / total # windows classified as swept. We repeat these calculations for 100 sets of windows 1Mb apart to generate an average FDR for independent regions. Because Flex-sweep provides a classification confidence score, we can additionally tune the FDR to correspond to a tolerance for a specific FPR (false positive rate) by increasing the confidence score at which a window is classified as a sweep. The default confidence threshold is 0.5. This FDR is then applied to the number of swept regions to determine how many were likely to be spurious signals. We consider this to be conservative, since with a sliding window size of 1.2 Mb and a step size of 10kb, 120 consecutive windows would have to be false positives for a single 10kb stretch of a sweep region to not be included in at least one true positive window.

###### Genome-wide patterns of selective sweeps

To understand the patterns of selective sweeps across the genome, we calculated the proportion of swept regions localized within genes, as well as the distances of swept regions from transcription start and sites and 5’ and 3’ UTR start and end sites. These serve as proxies for regions affecting regulatory and expression patterns. A swept region was considered to be within a gene if the site of the peak of the sweep confidence was within the interval bounds of any gene as defined by Ensembl (release 69) (Cunningham et al. 2022). Similarly, the site with the highest sweep confidence within a swept region was taken as the point from which to measure the distance to transcription and UTR start and end sites. To test for statistical significance, we randomized the positions of the swept regions we identified while maintaining their internal features (length, location of peak relative to 5’ and 3’ ends) and the position of the unclassifiable regions, and recalculated the proportions and distances of the shuffled swept regions to these genomic features 1,000 times. For each, we conducted one-tailed tests with 𝛼 = 0.05 - if the observed genic proportion was larger than 95% of the genic proportions calculated with the randomized regions, the difference was considered to be significant. Similarly, if the observed distance of a swept region to a transcription, 5’, or 3’ start or end site was smaller than 95% of the distances calculated with the randomized regions, the difference was considered to be significant. We repeated these analyses at sweep confidence thresholds of 0.5 (default), 0.6, 0.7, 0.8, 0.9, 0.95, and 0.99.

### Supplementary Results

###### Power of new statistics

hapDAF

The haplotype-derived allele frequency (*hapDAF*) statistic compares the frequency of shared flanking variants on ancestral and derived backgrounds in a fixed window around a focal variant (see **Methods** for detailed description, Supplementary figure 21). Flanking variants that are more common on the derived background are selected that are present on at least one ancestral background, and meet certain frequency thresholds. For *hapDAF-o*, the frequency of the flanking variant on the ancestral background is less than 25% and its total frequency on both backgrounds is 25%. For *hapDAF-s* these thresholds are 10% and 10%. These thresholds were tuned at 5% intervals to focus the statistic on the goal to either identify older, incomplete sweeps (*hapDAF-o*), or more recent, incomplete sweeps from standing variation (*hapDAF-s*; see **Methods**). *hapDAF-o* has good power at selection times back to at least 𝜏 = 0.1 * 4N_e_ generations ago at *s* = 0.01 (Supplementary figure 26. This includes incomplete and soft sweeps. In comparison, *hapDAF-s* has more power to detect recent sweeps (𝜏 = 0) (Supplementary figure 27), but less to find sweeps from standing variation that did not consistently increase the frequency of shared variants on a single derived background (as seen in comparison to eg. DIND, Supplementary figure 28).

Sratio

The segregating sites ratio (*Sratio*) compares the number of variable sites around a focal derived allele with the number of variable sites around the corresponding ancestral allele (see **Methods** for detailed description, Supplementary figure 22). This is designed to detect incomplete and soft sweeps. *Sratio* has comparable power to other statistics designed to detect sweeps under similar scenarios, including HAF, DIND, iHS, and nsl (Supplementary figure 29).

freq

The two *freq* statistics are designed to detect incomplete sweeps by looking for an excess of low or high-frequency alleles on the derived background (see **Methods** for detailed description, Supplementary figure 23). They complement each other by focusing on different ends of the site frequency spectrum (SFS). Frequency thresholds for derived backgrounds with the derived variant were tested at 5% intervals. For *lowfreq*, the frequency threshold with the highest power to detect recent, incomplete sweeps is a ceiling at 25%, while for *highfreq* it is a floor at 25%.

*highfreq* and *lowfreq* have the power to detect a variety of incomplete sweeps. This includes some power at weak (*s* = 0.005), older sweeps (𝜏 = 0.05) that is comparable to that of existing statistics. Each statistic captures a known aspect of selective sweeps, respectively an excess of rare alleles (*lowfreq*) and and excess of high frequency derived (*highfreq*) and therefore should add power in the synergistic context of a neural network or applications such as Approximate Bayesian Computation (Supplementary figures 30, 31).

###### Model calibration

To determine how well calibrated the model is, we assessed whether the observed rate of detecting sweep regions was similar to the model’s confidence score in its prediction of sweep regions. We find that the model is better calibrated when trained with the equilibrium demography than the Yoruba demography, with the model underestimating its confidence in sweep regions particularly at higher false positive rates. In the equilibrium-trained model, the confidence score is 0.55 at a 2% FPR, 0.67 at a 1% FPR, and 0.98 at a 0.1% FPR, while in the Yoruba-trained model these values are 0.13, 0.40, and 0.96. This systematic bias toward the confidence score being too low means that the model is conservative, and may therefore be under-reporting sweep regions even at the default confidence threshold of 0.5. This is, in part, why we have chosen to refer to the values output by the CNN as “confidence scores” rather than probability scores, because while their relative magnitudes reflect the likelihood that region contain a sweep, these may be underestimates of the true probabilities. This suggests that there is additional room for improving the power of such methods without sacrificing FPR.

###### Detecting older sweeps

With training as described in the main text, Flex-sweep is capable of detecting sweeps up to 10,000 generations old under the Yoruba demographic model, but its scope is limited (Supplementary figure 32). In particular, sweeps must be complete, or nearly so. At a false positive rate of 0.1%, its power to detect strong (s = 0.01), hard (starting allele frequency = 0; hereafter SAF), complete sweeps is 75%, which drops to <10% for incomplete sweeps (ending allele frequency = 0.7 (EAF) and EAF = 0.9). At a more relaxed false positive rate of 1%, its power increases to 84% for complete sweeps and 52% for nearly complete sweeps (EAF = 0.9), but remains very low (11%) for less complete sweeps (EAF = 0.7), (Supplementary figure 32).

###### Comparison to state-of-the-art

We also compare Flex-sweep to diploS/HIC, another powerful CNN-based method (Kern and Schrider 2018) and one of the first to popularize CNNs for detecting selective sweeps. In this case, we include training and testing both diploS/HIC and our method using a data set with only complete sweeps because they are what diploS/HIC is designed to detect (see **Methods**). In all other respects the training and testing data sets are the same as the equilibrium data described above. In addition, because diploS/HIC is designed to detect and classify sweeps into four types (hard sweeps, soft sweeps, and regions linked to those respective types), we modified it to instead classify only the binary state - neutral or swept region. Flex-sweep has greater power and lower false positive rates than this modified, binary (sweep or neutral) diploS/HIC and, as expected because it was not diploS/HIC’s purpose, this difference is more pronounced in data sets that include incomplete sweeps (Figure 4, Supplementary figure 12).

###### Statistic importance

To explore how each statistic contributes to Flex-sweep classification, we used Deep SHAP (SHapley Additive exPlanations) to interpret the contribution of each feature (a statistic calculated at a region in the locus over a specific window size) (Lundberg and Lee 2017). We calculated SHAP values (Lundberg and Lee 2017) for the equilibrium-trained, YRI demography-trained, and decline-trained models and interpreted the results in their corresponding testing data following two approaches: 1. clustering features by similarity, and 2. averaging the importance across each statistic at small (50kb-100kb), medium (200kb-500kb) and large (1Mb) window sizes in the middle (the 7 windows in the center of the locus) and shoulders (the leftmost and rightmost 7 windows of the locus) of each region. Both produced similar interpretations, which were similar to the ungrouped results. For ease of interpretation we present the results of the importance values averaged over each statistic at different window sizes in the middle vs shoulders of the locus (Supplementary figure 16). This results in 56 features (H12 and HAF only have one importance value each because they are calculated over the entire locus).

Both *highfreq* and *hapDAF*-s, calculated for the middle as well as the shoulders of the locus at various window sizes, make up most of the top 10 most important features for models trained and tested with simulations generated under the human demographic model, equilibrium demographic model, and demographic model with a population decline. For the human demographic model (Supplementary figure 16a), the top 10 most important features also include DIND and hapDAF-o; for the decline model, the top 10 most important features also include iSAFE (Supplementary figure 16b); and for the equilibrium demographic model, the top 10 most important features include only highfreq and hapDAF-s (Supplementary figure 16c). iHS, nSL, and H12 were consistently among the least important features for these three data sets. This is likely because these statistics only have good power to detect recent sweeps, and our simulated data sets also use very old sweeps for which they have little to no power. It is important to note, however, that this does not mean that these three statistics do not contribute to the overall model, nor that they are not potentially important with other training and testing data. In particular, they likely still contribute to detecting very recent sweeps, and may be more important in data sets focusing on these.

###### Sweep type classification power

We test the effect of classifying the type of sweep on power, grouping sweep types into hard complete, hard incomplete, soft complete, or soft incomplete. We find that the false positive rate (identifying a neutral region as any kind of sweep) is somewhat increased (to 9.6% under the equilibrium demographic scenario and 1.8% under the YRI demographic model), and sweep types are frequently misclassified. Unsurprisingly, hard sweeps are most frequently classified correctly. Hard incomplete sweeps are often misclassified as soft incomplete, and vice versa, and soft incomplete sweeps are often misclassified as neutral regions (Supplementary figure 13).

###### Low recombination rate

The mean recombination rate of the training simulations was 1 cM, which could influence the range of recombination rates at which the CNN can distinguish a neutral region from a sweep, so we also trained a low recombination rate model with simulations generated under the same parameter distributions as the original equilibrium demographic model, but with the mean of the exponential recombination rate an order of magnitude lower (0.1 cM). We find that training with low recombination rates drastically improves the false positive rate of testing data with low recombination rates, but decreases power for testing data with higher recombination rates (Supplementary figure 11). Because the exponential distribution of recombination rates with a mean of 0.1 cM in the training simulations still includes values > 1 cM, further testing could be conducted to determine if limiting the upper bound or further reducing the mean of the recombination rate in training simulations could improve the false positive rate in low recombination rate regions.

###### False discovery rate

72,749 of 258,665 windows (28%) were classified as sweeps at the default confidence threshold of 0.5, and 11,659 were unclassifiable. Windows are unclassifiable when there are too few SNPs to calculate one or more of the statistics in the feature vector over one or more window sizes. Most of those that were unclassifiable were near the centromeres of the chromosome. 1,654 swept regions were found, quantified as the number of unbroken streaks of windows classified. These regions, and the genes they contain, are provided in Supplementary table 1, and make up 23% of the total genome (with the length of a region calculated from the peak of the first window classified as a sweep to the peak of the last). During training on the human demographic model, Flex-sweep had a 0.7% false positive rate (FPR = number of false positives / total number of positive classifications). Averaging over 100 sets of independent windows 1 Mb apart in the Yoruba data, the average FDR is 1.9% (see **Methods** and **supplementary methods**). Obtaining a false discovery rate below 1% would require increasing the confidence threshold to approximately 0.8 (Supplementary figure 19).

###### Genome-wide patterns of selective sweeps

To understand the patterns of selective sweeps across the genome, we calculated the proportions of swept regions in genes and their distances from sites associated with regulatory activity. We find that selective sweeps are significantly more likely to be found within genes than would be expected by chance at all tested confidence thresholds (one-tailed randomization test (see **Methods**), all p < 0.01, Figure 5, Supplementary figure 17). Across the genome, 54% of sweeps are found within genes, while the expected mean by randomization is 28% at the default confidence threshold (Figure 5, see **Methods**). Between 54% and 59% of sweeps are found within genes at higher confidence thresholds (Supplementary figure 17). In addition, the peaks of each sweep region (see **Methods**) are more likely to be closer to the transcription start site (Figure 6, one-tailed randomization test p = 0.02, see **Methods**). While this pattern holds true for all confidence thresholds below a sweep confidence of 1 (Supplementary figure 18), it is only significant at the 0.5, 0.6, 0.8, 0.95, and 1 confidence thresholds (p_60_= 0.03, p_70_ = 0.07, p_80_ = 0.03, p_90_ = 0.12, p_95_ = 0.048, p_99_ = 0.11, p_100_ < 0.01). Given the reduced number of sweeps at higher thresholds and the sub-200bp differences between the observed means and the lower threshold of the expected means, it may require a higher resolution run of Flex-sweep (i.e. using a smaller step size between windows) to appropriately resolve the significance of these patterns. The peaks of each sweep region trend toward being closer to the 5’ and 3’ UTR start and end sites at confidence thresholds up to 0.99 as well, though these are not significant (0.05 < p <= 0.13), with the exception of the 3’ UTR start and end sites at the 0.8 threshold (p = 0.03 and p = 0.03), 0.95 threshold (p = 0.032 and p = 0.209), and 1 threshold (p < 0.001 and p < 0.001). Sweeps are not more likely to be close to transcription end sites than expected by chance (all p > 0.05 Figure 6, Supplementary figure 18).

###### Previously identified genes

In addition, we find evidence of sweeps with high probability at multiple genes with previous genomic and functional evidence of selection. Those supported by both lines of evidence include genes associated with responses to infectious diseases and pathogens, in particular TLR5 (Toll Like Receptor 5) (Hawn et al. 2003; Abu-Maziad et al. 2010; Grossman et al. 2013), ITGAE (Integrin Subunit Alpha E) (Grossman et al. 2013; Triska et al. 2015; Ravenhall et al. 2018; Harris and DeGiorgio 2020), and APOL1 (Apolipoprotein L1) (Thomson et al. 2014; Mizuno et al. 2010; Ko et al. 2013) (Supplementary figure 33). We also found evidence of sweeps at genes previously suggested to be sweep targets by genomic evidence alone, including CADM3 (de Magalhães and Matsuda 2012; Grossman et al. 2013; Sugden et al. 2018), CTNS (Grossman et al. 2013), DOCK3 (Higasa et al. 2009; Sugden et al. 2018), P2RX5 (Mizuno et al. 2010; Sugden et al. 2018), and SHPK (Kudaravalli et al. 2008; Sugden et al. 2018; Grossman et al. 2013; Barreiro et al. 2008), among others (Supplementary table 1).
